# Huib32: A Potent and Selective USP32 Inhibitor Modulating Endosomal Processes and Advancing Cell-Permeable USP32 Probes

**DOI:** 10.1101/2025.04.19.649632

**Authors:** Stephan Scherpe, Vito Pol, Raymond Kooij, Adán Pinto-Fernández, Rayman T.N. Tjokrodirijo, Darragh P. O’Brien, Iolanda Vendrell, Dharjath S. Hameed, Peter A. van Veelen, Benedikt M. Kessler, Paul P. Geurink, Aysegul Sapmaz

## Abstract

Deubiquitinating enzymes (DUBs) are pivotal regulators of ubiquitination, a vital post-translational modification essential for cellular processes. Dysregulated DUB activity disrupts cellular homeostasis, driving diseases like cancer and neurodegeneration. Ubiquitin-specific protease 32 (USP32) has emerged as a promising therapeutic target due to its role in endosomal and autophagosomal dynamics and its association with breast, ovarian, and lung cancers. Here, we describe **Huib32** (<u>H</u>uman de<u>U</u>biquitinase Inhi<u>b</u>itor 32) as a USP32 inhibitor. Cyanimide-containing **Huib32** potently and selectively inhibits USP32 by covalently binding to the active site Cys743 *in vitro* and in cells, enhancing substrate ubiquitination, altering endosomal morphology, and mimicking USP32 depletion. Additionally, we present two activity-based probes (ABPs), **Huib32*1** and **Huib32*2**, which enable precise detection of USP32 activity and confirm probe selectivity via mass spectrometry. Together, **Huib32** and its probes represent a unique approach for targeting USP32, offering new research tools and potential therapeutic avenues for cancer and disorders involving endocytic trafficking.

## INTRODUCTION

Ubiquitination, a post-translational modification, regulates many cellular events, including protein degradation, autophagy, endocytosis, cell signaling, DNA repair, and cell cycle progression.^1–3^ A key component of this system is deubiquitinases (DUBs), enzymes that reverse the ubiquitination process and thereby regulate the stability, localization, and function of various proteins. As such, they are an indispensable component of the spatiotemporal control of the complex ubiquitin-dependent cellular processes^4^. Dysregulation of DUBs has been shown to be involved in numerous diseases, such as cancer,^5^ neurodegenerative diseases,^6^ infectious diseases,^7,8^ and autoimmune diseases.^9^ In recent years, the development of selective DUB inhibitors has become an exciting avenue in drug discovery,^10,11^ with the aim of targeting specific members of the DUB family, which encompasses ∼100 members, for therapeutic benefit. Despite multiple challenges in developing DUB-specific inhibitors, including off-target effects due to the high similarity in DUB active sites, the complexity of ubiquitin signaling pathways, or the lack of structural information,^12^ selective inhibitors have been described for numerous DUBs,^13^ including UCHL1, USP14, USP7, USP30 and BAP1. These inhibitors enable the exploration of the specific roles of individual DUBs in cellular processes and diseases, including cancer, neurodegenerative disorders, and immune diseases, with some even advancing to clinical trials, such as USP1 and USP30 inhibitors^14^.

Ubiquitin-specific protease 32 (USP32) is a DUB from the cysteine protease DUB family that has emerged as a key regulator of cellular trafficking, particularly in the context of endo-lysosomal and auto-lysosomal systems.^15,16^ USP32 has been implicated in a range of cancers, including breast cancer,^17^ lung cancers,^18,19^ gastric cancer,^20^ epithelial ovarian cancer,^21^ and others.^22–24^ Despite its essential role in regulating these pathways and participation in cancer pathogenesis,^25^ the development of specific and potent inhibitors targeting USP32 remains an unmet challenge. Therefore, the discovery of a selective inhibitor for USP32 would represent a significant step forward in the field of DUB-targeted therapies, enhancing our understanding of its biological roles and revealing its potential as a therapeutic target. Here, we report the discovery of **Huib32**, a potent and selective small-molecule inhibitor of USP32. Mechanistic studies demonstrate that **Huib32** irreversibly covalently inhibits USP32 by binding to the active site Cys743 residue through its cyanimide warhead, with no cross-reactivity to other DUBs. *In vitro* and cell-based assays confirm the specificity of **Huib32** for USP32. Functional and biochemical assays and proteome-wide analyses revealed the ability of **Huib32** to modulate the ubiquitination of key USP32 substrates, such as RAB7, RAB11, and ARL8A/B, and its impact on endosomal trafficking. To further advance the potential for visualizing USP32 activity, we synthesized two activity-based probes (ABPs) based on an analog of **Huib32**: fluorescent probe **Huib32*1** (with the installment of SulfoCy5) and **Huib32*2** (with the installment of biotin), to enable activity-based visualization and enrichment of USP32 in cells. **Huib32*1** effectively labels active USP32, while **Huib32*2** facilitates efficient USP32 pull-down from cellular lysates, validated by quantitative LC-MS/MS analysis. These findings position **Huib32** and its derivatives as invaluable tools for exploring the role of USP32 in physiological and pathological conditions and providing the foundation for future therapeutic development targeting USP32.

## RESULTS

### Discovery and properties of Huib32

To identify a small-molecule inhibitor of USP32, we performed a high-throughput screen (HTS) of small molecules, measuring the activity of USP32 using fluorogenic ubiquitin-rhodamine-morpholine (UbRhoMP) as a substrate.^26^ We screened an in-house library containing crudes of 7,536 compounds (Figure 1A), which was previously synthesized in a 1,536-well plate by performing an amidation reaction in DMSO with 16 cyclic cyanimide-containing amine building blocks (BB-codes) and 471 carboxylic acids (CA-codes) using *N*,*N*’-diisopropylcarbodiimide (DIC) and hydroxybenzotriazole (HOBt)^27^ (Supplementary Data 1). The recombinant USP32 full-length (USP32FL) protein,^15^ along with a panel of 16 DUBs and SENP1 (SUMO protease), was pre-incubated with the compounds (1.25 µM final concentration), followed by incubation with UbRhoMP (for DUBs) or SUMO2RhoMP (for SENPs) substrate. We identified 105 compounds that showed >50% inhibition of USP32 activity. Seven of these compounds, all sharing the same carboxylic acid moiety (CA282), selectively inhibited USP32 activity among a panel of 16 DUBs from the different families and SENP1 (Figure 1B, Supplementary Data 1). As we experienced undesirably high cysteine reactivity of compounds derived from BB013 and BB015,^27^ we continued with 5 out of 7 identified BB’s for further investigation (Figure 1C).

**Figure 1.**
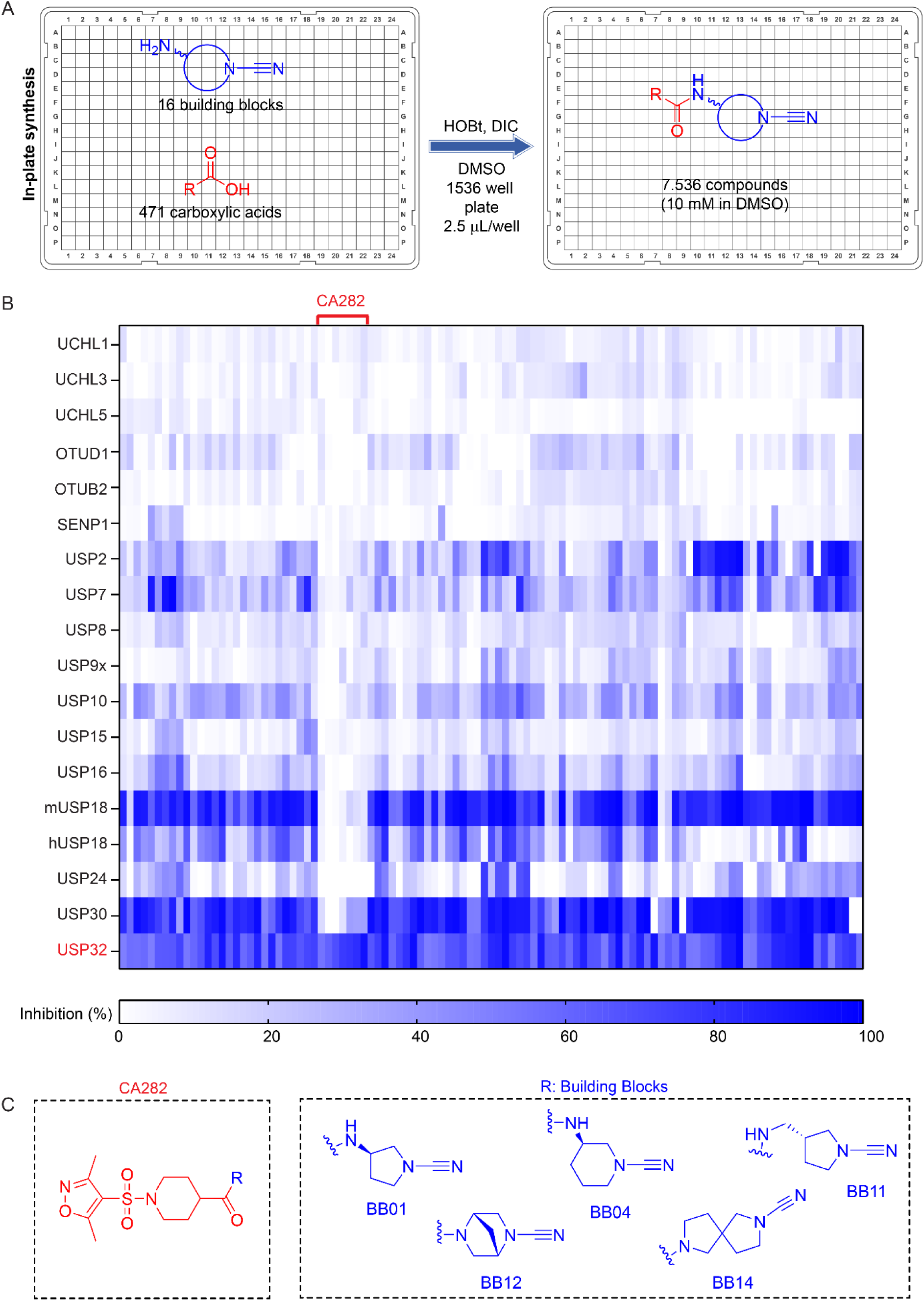
Discovery of USP32 inhibitors via high-throughput screening of a cyanamide library. **A.** Schematic illustration of in-plate synthesis to build a compound library. A total of 7,536 compounds were synthesized in a 1536-well plate by performing an amidation reaction in DMSO with 16 amine building blocks (BB-codes) and 471 carboxylic acids (CA-codes) using DIC and HOBt. **B.** Heatmap of 105 compounds showing > 50% inhibition of USP32 activity. Seven compounds carrying the same carboxylic acid (CA282) selectively inhibited USP32 in a panel of 21 DUBs from different DUB families. **C.** Chemical structure of resynthesized and purified top 5 compounds that showed selectivity towards USP32. See also Supplementary Figure 1.

To improve the potency towards USP32, we designed and synthesized (in-plate) a small library containing 125 compounds by coupling each of the five BB’s (BB01, BB04, BB11, BB12, and BB14) to CA282 and 24 analogues of CA282. (Supplementary Figure 1, Supplementary Data 2). Inhibition of USP32FL was assessed for all compounds at a 1 and 10 µM final concentration. Modifications to the piperidine ring, such as altering the carboxylic acid position (CA470 and CA471) or substituting the piperidine ring with other groups (CA472-CA476), led to a complete loss of activity towards USP32. Changing the sulfonyl group to a carboxyl (CA477) or substituting the isoxazole with an aliphatic moiety (CA485, CA486, and CA489) also resulted in a decrease in potency. While aromatic substitutions of the isoxazole moiety largely retained inhibitory activity, compounds with 2,4,6-trimethyl-phenyl (CA481) or phenyl (CA483) showed significant reductions. Notably, substituting the isoxazole for 4-methoxyphenyl (CA480) mostly preserves potency towards USP32, while also allowing for further structural manipulation of the methoxy group (vide infra). These findings highlight the importance of the 1-sulfonyl-4-carboxyl-substituted piperidine ring, as well as the preference for an aromatic sulfonyl substituent, in preserving inhibitory efficacy. However, none of the compounds outperformed the original hits containing the CA282 carboxylic acid (Supplementary Figure 1, Supplementary Data 2). Based on these results, we decided to continue with CA282 and resynthesized and purified CA282 coupled to each of the 5 BBs for further analyses (Figure 1C).

### Properties of the USP32 inhibitors

To assess the inhibition efficiency of the compounds towards USP32 FL protein, we determined half-maximal inhibitory concentration (IC_50_) values of the five pure compounds using our UbRhoMP-based fluorescent intensity assay. Three compounds, **BB01CA282**, **BB12CA282**, and **BB14CA282**, exhibited submicromolar inhibition efficiency (IC_50_ values of 21.2, 94, and 518 nM, respectively) towards USP32 (Figure 2A, B, Supplementary Figure 2A, and Table S1). Further analysis of these compounds revealed that **BB01CA282** inhibits the enzyme with enzyme inactivation rate (*k*_inact_) to inhibition constant (*K*_i_) ratio *(k*_inact_/*K*_i_) of 18,800 s^-1^M^-1^ while **BB12CA282** and **BB14CA282** inhibit USP32 with lower *k*_inact_/*K*_i_ ratio of 2,130 and 219 s^-1^M^-1^ respectively, indicating higher covalent inhibition potency of **BB01CA282** towards USP32 (Figure 2C, Supplementary Figure 2B and Table S1). We performed a jump dilution assay to investigate the reversibility of inhibition of USP32 for inhibitors **BB01CA282**, **BB12CA282** and **BB14CA282**. In this assay, USP32 at a concentration of 80 nM and inhibitors at a concentration of 10 times their IC_50_ values were mixed and incubated for 30 min, which was followed by 100-fold dilution with a buffer containing the UbRhoMP substrate and measurement of fluorescence intensity. Following the dilution of USP32-inhibitor complexes, rapid recovery of USP32 activity was observed for all three inhibitors, comparable to DMSO or low inhibitor concentration (0.1 times of IC_50_), suggesting a fast reversible covalent bond formation between enzyme and inhibitors (Figure 2D and Supplementary Figure 2C). These results demonstrate the highest inhibition potency of **BB01CA282** towards USP32, compared to the other inhibitors. Therefore, we decided to continue with **BB01CA282** for further analysis and henceforth referred to the molecule as **Huib32** (<u>H</u>uman de<u>U</u>biquitinase Inhi<u>b</u>itor 32). The synthesis route of this compound is shown in the Supplementary Information.

**Figure 2.**
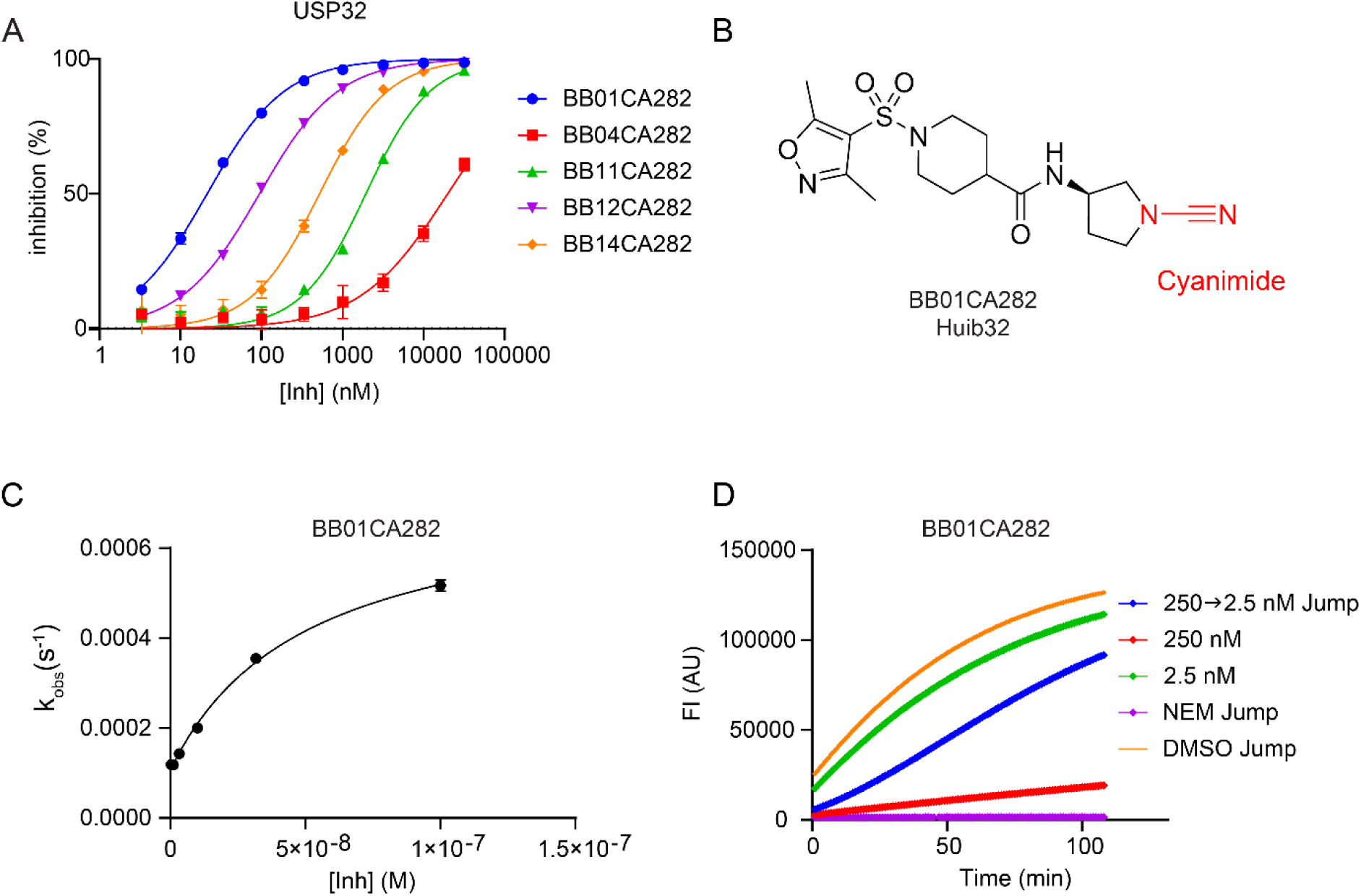
Properties of USP32 inhibitors. **A.** IC_50_ determination of the indicated compounds for USP32 using UbRhoMP-based fluorescent intensity assay. **B.** Structure of **BB01CA282**/**Huib32**. **C.** Kinetics plot of *k_obs_* versus different concentrations of **BB01CA282**/**Huib32**. **D.** Jump dilution progress curve for USP32 proteolytic activity for **BB01CA282**/**Huib32**. See also Supplementary Figure 2.

### Huib32 selectively targets USP32 *in vitro* and in cells, among other DUBs

To gain insight into the specificity of **Huib32**, we first tested **Huib32** with a panel of 46 DUBs from distinct families of DUBs, including USP32FL and USP32 catalytic domain (USP32CD). **Huib32** selectively inhibited both USP32FL and USP32CD (Figure 3A). The only other DUB inhibited by Huib32 was USP6CD (Figure 3A), which is not surprising since *USP6* was reported to be a chimeric fusion gene with 97% nucleotide similarity to the catalytic domain of *USP32.*^28^ To further test the selectivity of **Huib32** in a cellular context, we incubated either lysate from MelJuSo cells for 1 h or live MelJuSo cells for 24 h with an increasing concentration of **Huib32** (0-50 µM), followed by incubation of cell lysates with rhodamine-ubiquitin-propargylamine (RhoUbPA) probe for 5 min at 37°C. USP32 inhibition is reflected by the disappearance of the probe-labelled USP32 band, as observed through fluorescence scanning of SDS-PAGE and immunoblotting against USP32. We observed a decrease in the probe labeling of endogenous USP32 in a Huib32 dose-dependent manner, starting from **Huib32** final concentration of 0.1 µM (Figure 3B and Supplementary Figure 3A). Remarkably, the activity of other DUBs was not affected by **Huib32** treatment up to a final concentration of 50 µM, as indicated by the fluorescent scan of the rhodamine signal, which represents the active DUBs labeled with RhoUbPA probe (Supplementary Figure 3A). Depletion of USP32 by specific siRNA treatment confirmed that the disappearing band indeed corresponded to USP32. (Supplementary Figure 3B). These results confirmed not just selectivity but also cell permeability and cellular target engagement of **Huib32**. We further tested USP32 inhibition by **Huib32** in HEK293T cell line, which yielded similar results to those observed in MelJuSo cells (Supplementary Figure 3C). To quantitatively measure the specificity of **Huib32**, we performed pull-down experiments in which MelJuSo cells were treated with **Huib32** (10 µM final concentration) or DMSO, followed by cell lysis and incubation with a dual functionalized DUB probe conjugated with both rhodamine and biotin at the N-terminus of ubiquitin (RhoK(Biotin)UbPA). The synthesis route of this probe is shown in the Supplementary Information. Taking advantage of the dual functionalized probe, we first tested the pull-down efficiency via a gel-based analysis of fluorescent signal representing the activity of DUBs in MelJuSo cell extract, again resulting in the disappearance of the band corresponding to USP32 in the samples treated with **Huib32** without any effect on other DUBs (Supplementary Figure 3D). The pull-down samples were then subjected to quantitative mass spectrometry analysis (Figure 3C). A total of 21 endogenous DUBs were identified in the mass spectrometry analysis, 17 of which had adequate abundance to carry out quantitative analysis (Supplementary Data 3). **Huib32** solely affected USP32 while showing no effect on the other 16 DUBs detected (Figure 3D). Our results conclusively indicated that **Huib32** is a USP32-specific inhibitor among the cysteine DUBs.

**Figure 3.**
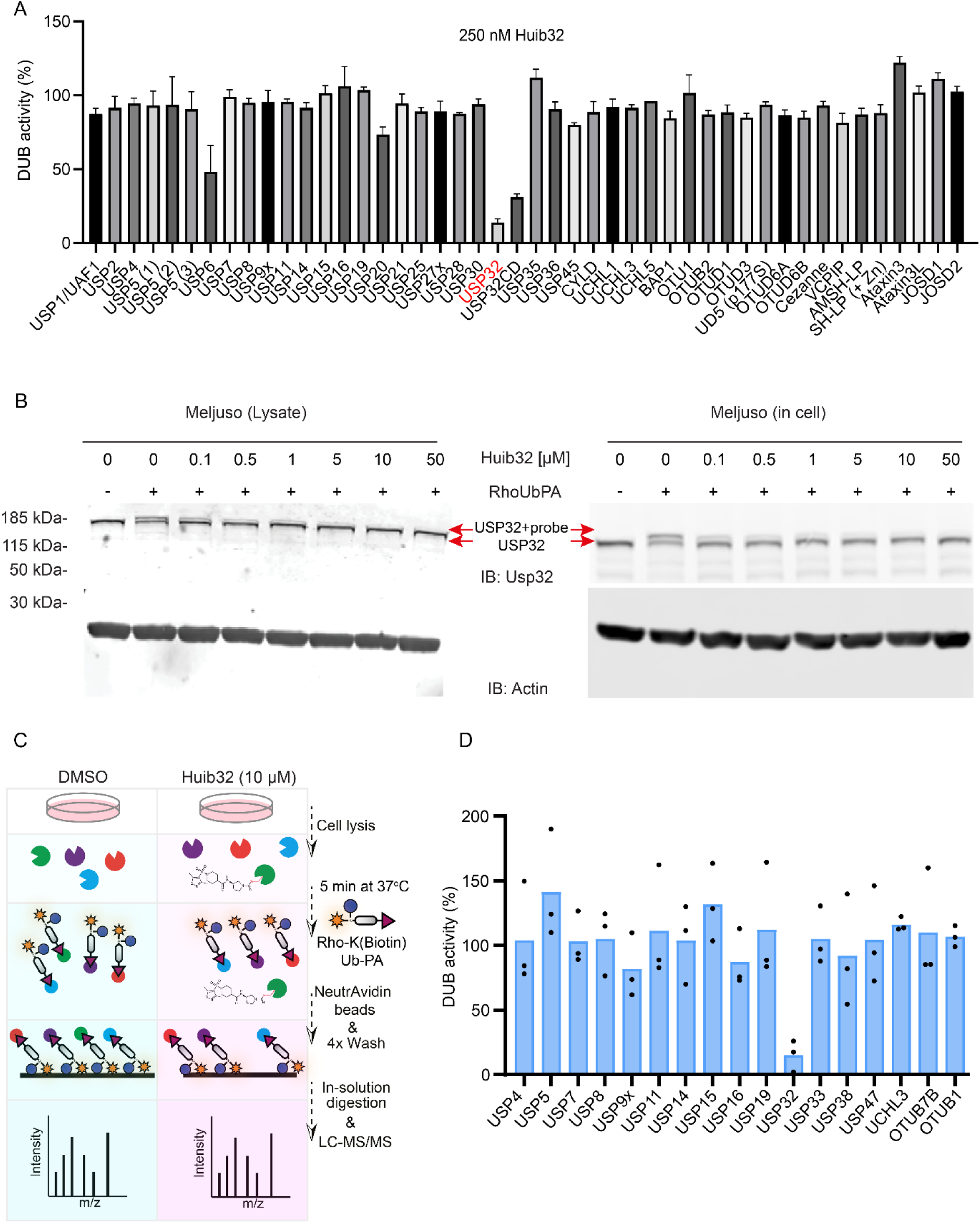
*In vitro* and cellular specificity of USP32 inhibitors. **A.** Specificity screen with purified recombinant DUBs (commercial or in-house) of remaining DUB activity after treatment with **Huib32** inhibitor. **B.** Target engagement of **Huib32** in cell lysate of MelJuSo cells or intact MelJuSo cells. Cell lysates (left panel) were incubated with the indicated concentration of Huib32 for 1 h, while intact MelJuSo cells (right panel) were incubated with Huib32 for 24 h. Following incubation, both cell lysates were treated with the RhoUbPA probe for 5 min. The samples were subjected to SDS-PAGE, followed by immunoblotting against USP32 and β-actin as a loading control. See also Supplementary Figure 3A. **C.** Schematic representation of the competition assay workflow to determine **Huib32** specificity. Intact MelJuSo cells were treated with 10 µM **Huib32** for 24 h, lysed, and incubated with a RhoK(Biotin)UbPA probe for 5 min. DUBs modified by the probe were pulled down using NeutrAvidin beads, and the samples were analyzed using mass spectrometry in three technical replicates. **D.** Quantification of remaining DUB activity in MelJuSo cells treated with **Huib32**. The remaining activity was calculated by comparing DUB probe-binding in DMSO-treated versus **Huib32**-treated samples, showing the inhibitor’s effect on DUB activity.

### Huib32 recapitulated the catalytic function of USP32 in the endocytic pathway

To gain more insight into the biological impact of **Huib32**, we investigated the dispersion and enlargement of the late endosomal compartment, where a striking phenotype was observed with the depletion of USP32. To test this, we incubated MelJuSo cells with the indicated concentration of **Huib32** for 72 h, during which we observed stable target engagement of **Huib32** (Supplementary Figure 4A and 4B), and previous USP32 depletion experiments were performed.^15^ The dispersion and swelling of late endosomal compartments positive with major histocompatibility complex II (MCHII) were observed in **Huib32**-treated cells (Figure 4A), once again implying that the catalytic activity of USP32 is essential for the functionality of the endocytic pathway. Furthermore, we investigated the impact of Huib32 on the ubiquitination of RAB7, a key substrate of USP32.^15^ Treatment with **Huib32** strongly increased the ubiquitination of RAB7 after 4 h of incubation in MelJuSo cells stably expressing GFP-RAB7, which steadily stayed up to 72 h of incubation (Supplementary Figure 4C). To test whether this modification increase by **Huib32** is indeed at the lysine residue targeted by USP32, the ubiquitination pattern of RAB7 was followed on affinity-isolated GFP-RAB7 WT and GFP-RAB7 2KR (carrying lysine to arginine mutation on K191 and K194) from **Huib32** (non)treated cells ectopically expressing HA-tagged ubiquitin. Approximately 2-fold increased RAB7 ubiquitination was observed in GFP-RAB7 WT from the cells treated with **Huib32** compared to nontreated cells, but not in ubiquitination mutant GFP-RAB7 2KR (Figure 4B).

**Figure 4.**
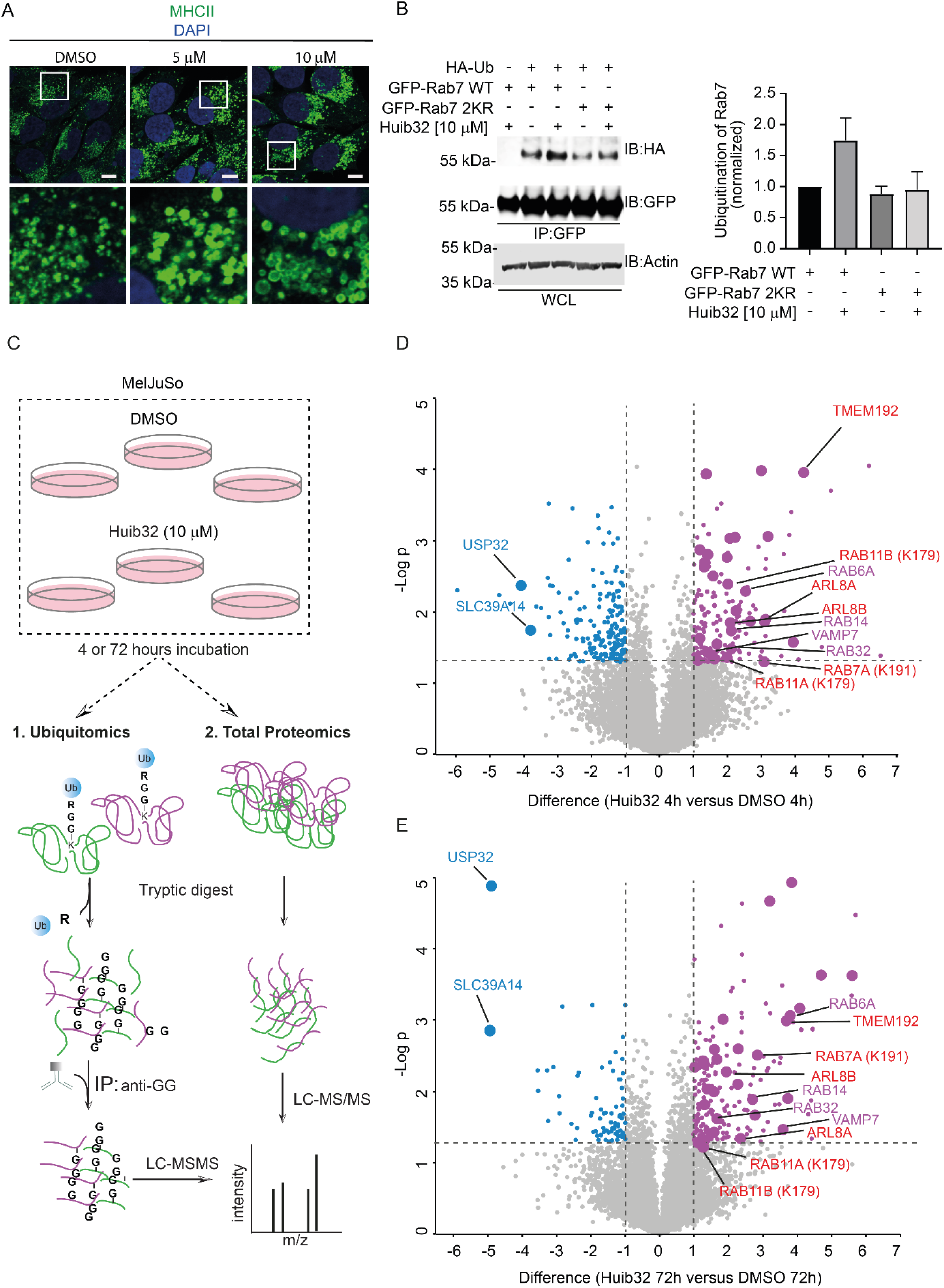
Recapitulation of the USP32 catalytic function in the endocytic pathway via Huib32. **A.** Effect of **Huib32** treatment on the size and distribution of endosomes. Representative confocal overlays of fixed MelJuSo cells that were treated with either DMSO or an indicated concentration of **Huib32** for 72 h. Fixed MelJuSo cells were immunostained against MHC-II (green) and stained with DAPI (blue). **B.** Ubiquitination status of GFP-Rab7 as an effect of **Huib32** treatment. GFP-Rabs, immunoprecipitated (IP) from HEK293T cells coexpressing HA-Ub and either GFP-RAB7 WT or GFP-RAB7 2KR, which were treated with Huib32 (10 µM) or DMSO for 72 h, were assessed by immunoblot against HA. WCL: whole cell lysate. β-actin was used as a loading control. See also Supplementary Figure 4C. **C.** Schematic representation of label-free quantitative proteomics workflow used to compare total proteome (see the Volcano plots in Supplementary Figure 6) and ubiquitinated substrates of MelJuSo cells that are treated with **Huib32** (10 µM) or DMSO for 4 h or 72 h. **D, E.** Volcano plots comparing the abundance of detected peptides carrying a GlyGly (GG) modification expressed as log_2_ ratios of MelJuSo cells treated with 10 µM final concentration of **Huib32** versus DMSO for 4 h **(D)** or 72 h **(E).** Dashed lines indicate a significance cutoff at a p-value of 0.05 (-log10 = 1.3) in the y-axis and –1 ≥ log_2_ ≥ 1 cutoff at fold change in the x-axis, n=3 independent experiments. All modified peptides that are enriched with the **Huib32** treatment are highlighted in purple, while the modified peptides downregulated by the **Huib32** treatment are highlighted in blue. The enriched or depleted unique peptides modified at the same lysine residue in the cells treated with **Huib32** for 4 h and 72 h are represented with larger circles. The peptides marked with corresponding protein names in red color were identified in the previous ubiquitomics studies as USP32 substrates.^15,16^ See also Supplementary Figure 5.

Having established that cells treated with **Huib32** displayed similar effects as observed by the knockdown of USP32, we performed a proteome-wide analysis with MelJuSo cells treated with **Huib32** or DMSO. Following 4 h and 72 h of incubation with 10 µM of **Huib32** or DMSO, harvested cells were either used for total proteome analysis or followed by enrichment using anti-Gly-Gly antibody recognizing Lys-ε-Gly-Gly remnants of ubiquitylated proteins upon trypsin digestion to profile the changes in **Huib32**– or DMSO-treated samples (Figure 4C and Supplementary Figure 4D). A total of 8,060 unique proteins in the total proteomics analysis (Supplementary Data 4 and 5) and 11,488 unique Lys-ε-Gly-Gly remnant peptides in ubiquitomics analysis from MelJuSo cells treated with **Huib32** or DMSO for 4 h and 72 h (Supplementary Data 6 and 7) were identified. Comparison of the ubiquitomes of **Huib32**-treated and DMSO-treated samples revealed significant differences in the abundance of Lys-ε-Gly-Gly remnant peptides. In **Huib32**-treated samples, 179 and 188 unique Lys-ε-Gly-Gly remnant peptides were significantly enriched (≥ 1.0 fold change in the log2 intensities), while 177 and 76 were depleted greatly (≤ –1.0 fold change in the log2 intensities), at 4 h and 72 h, respectively, compared to DMSO-treated controls (Figures 4D and 4E, Supplementary Figures 5A and 5B, Supplementary Data 6, 7). Notably, 33 unique Lys-ε-Gly-Gly remnant peptides, originating from 30 distinct proteins, were consistently enriched in both the 4 h and 72 h **Huib32** treatments (Supplementary Figures 5A and 5B). Many of these 30 proteins are involved in membrane trafficking, including small GTP-binding proteins such as RAB6A, RAB32, Rab14, ARL8A/B and late endosomal proteins such as TMEM192 and VAMP7 (Figures 4D and 4E). Interestingly, unique Lys-ε-Gly-Gly remnant peptides detected in TMEM192, and ARL8A/B were previously found to be targeted by USP32.^15,16^ Lysine 191 residue of RAB7, the validated substrate of USP32,^15^ and lysine residue 179 of RAB11A/B, a potential substrate identified previously,^16^ were also enriched in both 4 h or 72 h **Huib32**-treated samples (Figures 4D and 4E). Interestingly, one of the two peptides depleted at 4 h and 72 h treatment by **Huib32** was a peptide containing the lysine residue 983 of USP32, suggesting potential activity-dependent but indirect regulation of USP32 by itself (Figures 4D and 4E, Supplementary Figures 5A and 5B). Additionally, total proteomics data revealed that proteins carrying the enriched unique Lys-ε-Gly-Gly remnant peptides in **Huib32**-treated samples did not show any changes in their protein level, suggesting ubiquitination on these substrates regulated by USP32 function is not a degradation or stabilization signal for the cell (Supplementary Figures 6A and 6B). Overall, these findings not only confirmed the established function of USP32 in membrane trafficking, implying the efficient inhibition of USP32 via **Huib32** in cells, but also provided additional insights into uncovered substrates of USP32.

### Rational Desing of USP32 activity-based probes, Huib32*1 and Huib32*2

Given the high potency and in vitro and in-cell selectivity of **Huib32** towards USP32 and the covalent nature of the compound, we hypothesized that this compound could serve as an attractive starting point for generating an activity-based probe for USP32. However, the **Huib32** structure does not provide a convenient site to install a reporter group, such as a fluorescent dye or biotin. Therefore, we decided to revisit the screening data from the small library, which contains 125 compounds (Supplementary Figure 1 and Supplementary Data 2). Our attention was drawn to compound **BB01CA480**, having the same core structure as **Huib32**, but with the 3,5-dimethylisoxazole moiety replaced with *p*-methoxybenzene. Compared to **Huib32**, **BB01CA480** showed a slight reduction in USP32 inhibitory potency at 1 µM, but the *p*-methoxy group provided a convenient site for the installment of a reporter group. Based on this observation, we first synthesized an azide-containing analog of this compound, named **Huib32\*** (Scheme 1). First, bromoethoxy-benzene (**4**) was reacted with chlorosulfuric acid and the resulting chlorosulfonate was coupled to methyl piperidine-4-carboxylate (**5**) to generate the sulfonamide **6**. Next, the bromide was substituted for an azide (**7**) and after saponification of the methyl ester, carboxylic acid **8** was amidated with 3-aminocyanopyrrolidine (obtained after Fmoc deprotection of **1**) resulting in **Huib32\***. Two probes were generated by reacting the azide moiety of **Huib32\*** in a copper(I)-catalyzed alkyne-azide cycloaddition (CuAAC) with SulfoCy5 alkyne to obtain fluorescent probe **Huib32*1**, and biotin-alkyne to obtain **Huib32*2** for enrichment of USP32 (Scheme 1 and Figure 5A). IC_50_ determination of **Huib32\*** and its probe derivatives, **Huib32*1** and **Huib32*2**, indicated that these compounds were less potent than **Huib32** towards USP32, which was expected based on the initial screening data (Supplementary Figure 1). Interestingly, the installation of SulfoCy5 or biotin resulted in a slight increase in inhibitory potency, with IC_50_ values of 412 nM and 266 nM, respectively, compared to their azide precursor (IC_50_, 670 nM) (Figure 5B).

**Figure 5.**
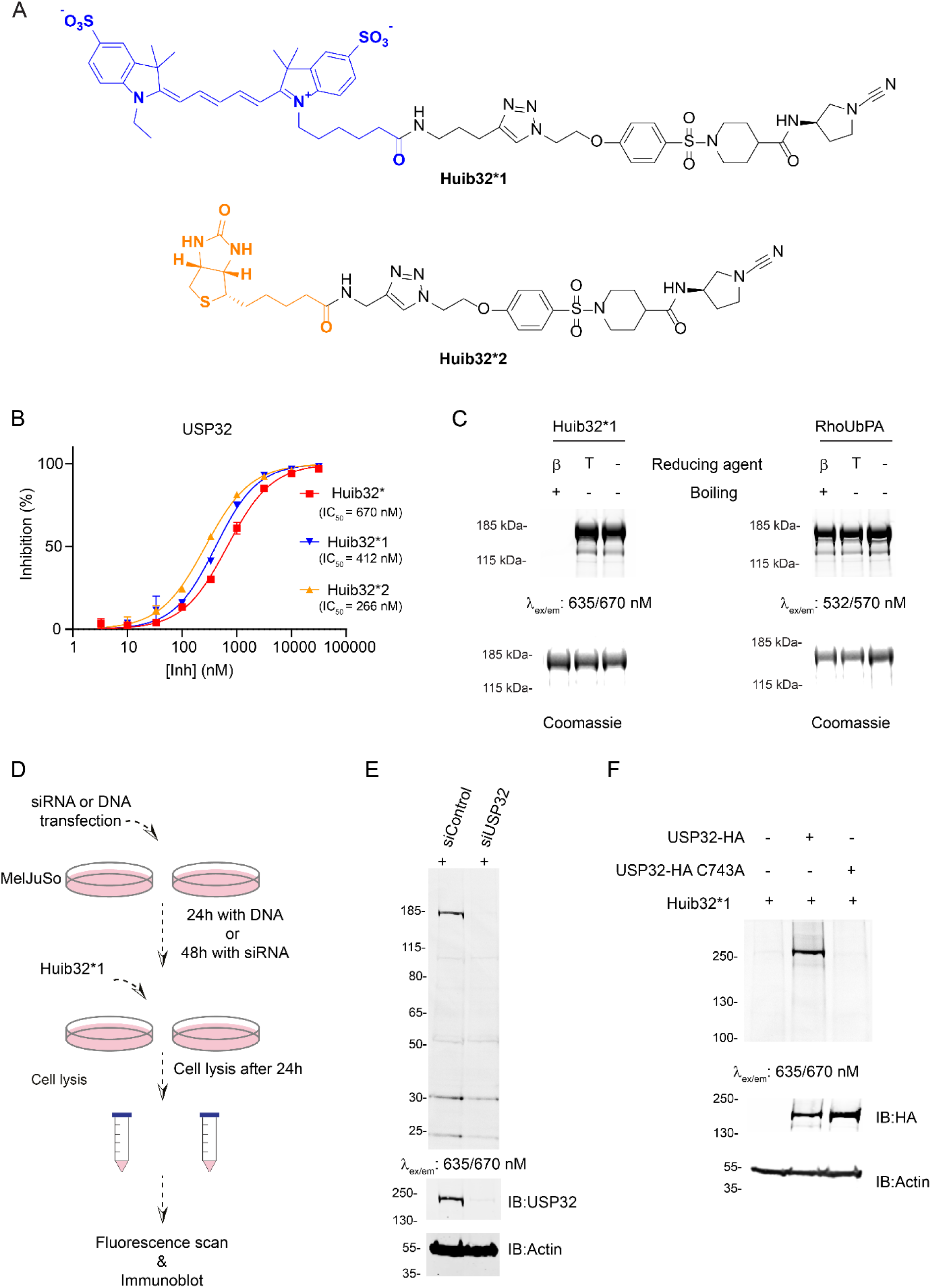
Properties of Huib32*, Huib32*1, and Huib32*2, and Characterization of the Fluorescent Probe Huib32*1. **Chemical structures of** Huib32*1 **and** Huib32*2 probes. **B**. IC_50_ determination of **Huib32\***, **Huib32*1,** and **Huib32*2** for USP32 **C.** Labeling of purified recombinant human USP32 with **Huib32*1**. Recombinant human USP32 was incubated with **Huib32*1** and RhoUbPA for 1h at 37°C, followed by SDS-PAGE, fluorescence scanning, and Coomassie staining. β: β-Mercaptoethanol and T: TCEP. **D.** Schematic overview of fluorescent probe labeling of endogenous USP32 in siRNA-treated cells or USP32 overexpressed cells with **Huib32*1**. **E.** Fluorescent probe labeling of endogenous USP32 in control (siControl) and USP32 knockdown (siUSP32) MelJuSo cells with **Huib32*1**. **F.** Fluorescent probe labeling of ectopically expressed USP32-HA wildtype (WT) or catalytically inactive mutant (CA) in HEK293T cells with **Huib32*1.**

### Fluorescent probe Huib32*1 can label active USP32 *in vitro* and in cells

To investigate whether the SulfoCy5 probe **Huib32*1** can label and visualize USP32 activity, we incubated purified recombinant USP32 with **Huib32*1** or RhoUbPA (control for USP32 activity) for 1 h at 37°C. As we previously experienced, the binding of compounds with cyanamide warheads to their target is susceptible to denaturing conditions; thereby, we resolved the samples by SDS-PAGE under denaturing boiling in the presence of β-mercaptoethanol) and non-denaturing conditions (no boiling and the absence of β-mercaptoethanol). Similar to our previous findings using a UCHL1 cyanimide probe,^26^ a clear band corresponding to probe-labeled USP32 was observed under non-denaturing conditions, whereas no labeling was detected under denaturing conditions (Figure 5C).

We further evaluated the cell permeability and in-cell activity of SulfoCy5 probe **Huib32*1**. For this purpose, MelJuSo cells treated with siControl (WT) or siUSP32 (USP32 knockdown) were incubated with 10 µM of **Huib32*1** for 24 h, followed by cell lysis and SDS-PAGE under non-denaturing conditions (Figure 5D). A clear band was observed in WT MelJuSo cells, which was not visible in the USP32 knockdown MelJuSo cells, indicating that probe **Huib32*1** could enter the cells and selectively label USP32 (Figure 5E). Encouraged by these results, we further investigated whether this interaction is activity-based binding to USP32. HEK293T cells expressing either HA-tagged USP32 (USP32-HA) WT or catalytically inactive mutant USP32 (USP32-HA C743A), or neither of them, were incubated with **Huib32*1** for 24 h followed by cell lysis and SDS-PAGE under non-denaturing conditions. USP32 labeling was clearly observed in samples expressing USP32 WT but not in those expressing catalytically inactive USP32, indicating that **Huib32*1** binds USP32 via its active-site cysteine (Figure 5F).

### Assessment of DUB Selectivity and Potential Off-Targets for Huib32*2

We further assessed the selectivity of biotinylated probe **Huib32*2**(Figure 5A), by incubating MelJuSo cell lysate with either **Huib32*2**, biotin-PEG4-alkyne or DMSO, followed by a pulldown using NeutrAvidin beads under optimized washing condition (lysis buffer containing 1% of SDS) and probe concentration (1 µM) (Supplementary Figure 7A). The samples were then analyzed by immunoblotting, silver staining, and quantitative LC-MS/MS analysis (Figure 6A). Immunoblot analysis using antibodies against USP32 confirmed efficient enrichment of USP32 by **Huib32*2** but not by DMSO or biotin-PEG4-alkyne controls (Figure 6B). Silver staining revealed a prominent band corresponding to the molecular weight of USP32, along with several additional bands. Some of these bands were similarly enriched in control samples, indicating nonspecific binding to NeutrAvidin beads. In contrast, others appeared to bind exclusively to the **Huib32*2** probe, suggesting potential off-target interactions of the biotinylated probe (Supplementary Figure 7B). From the quantitative LC-MS/MS data, we calculated the intensity-based absolute quantification (iBAQ) values for the pulldown samples. The iBAQ values of the pull-down samples from **Huib32*2**-treated lysate were then compared to those of the DMSO or biotin-PEG4-alkyne-treated controls. Identified proteins were ranked based on their iBAQ values to determine the most highly enriched targets in **Huib32*2**-treated samples (Supplementary Data 8). A detailed examination of the list of ubiquitin-related enzymes, including deubiquitinating enzymes (DUBs), E1, E2, and E3 enzymes, revealed a high degree of specificity of the probe for USP32 within the ubiquitin system, as illustrated in Figure 6C. Only a few of these enzymes were detected in the pulldown experiment, with iBAQ values significantly lower than those of USP32. Since endogenous protein levels can influence pulldown efficiency, a total proteome analysis of MelJuSo cells used in the experiment was performed to investigate endogenous protein levels (Supplementary Figure 8A and Supplementary Data 9). Despite the relatively low abundance of USP32 in MelJuSo cells compared to other DUBs, USP32 was the most highly enriched enzyme in the pull-down. This observation suggests a strong preference of the probe for USP32 over other enzymes.

**Figure 6.**
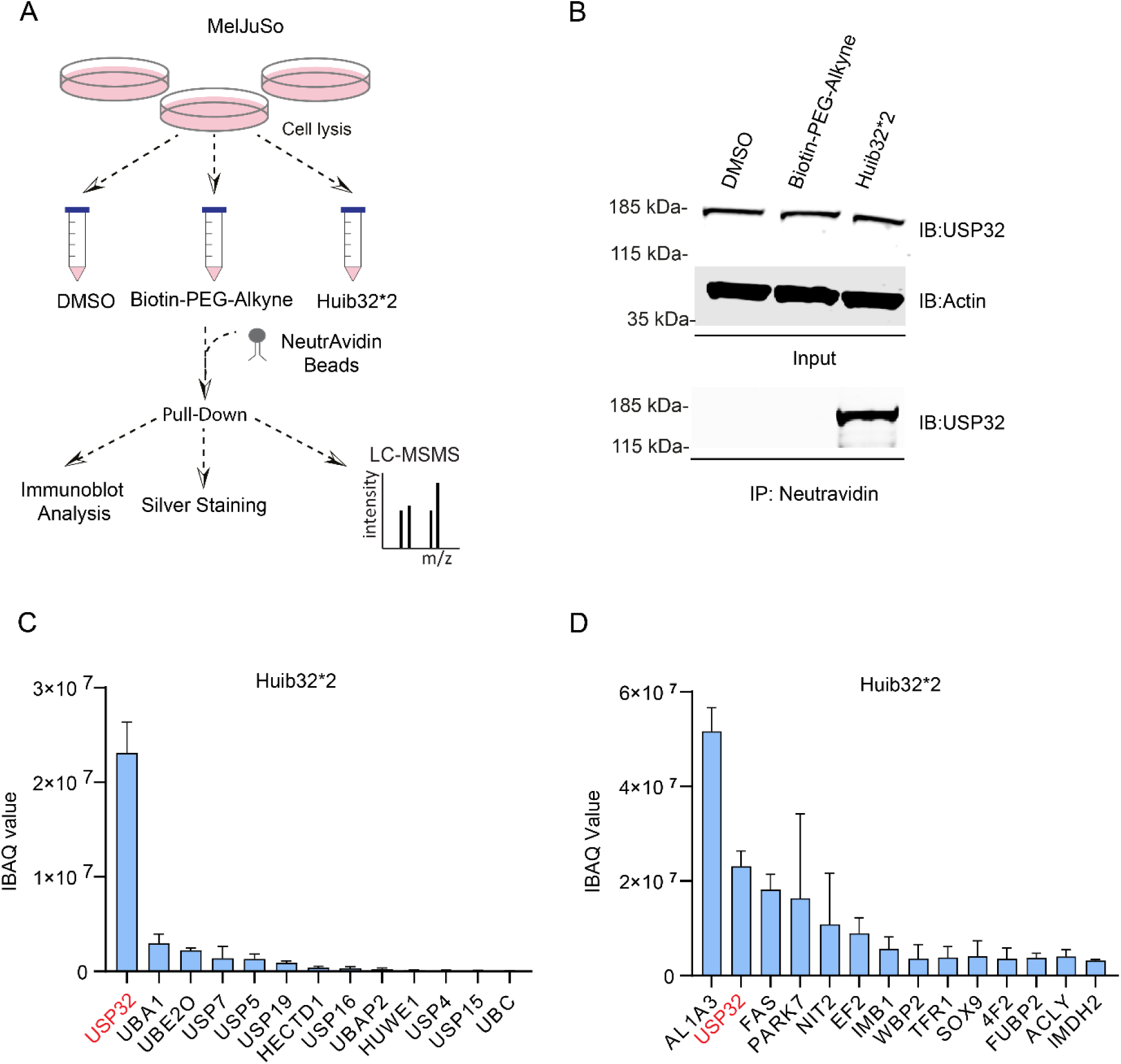
Assessment of DUB Selectivity and Identification of Potential Off-Targets for Huib32*2. **A.** Schematic overview of the pull-down experiment to identify **Huib32*2** probe binding proteins. MelJuSo cells were lysed, and cell lysates were treated with either 1 µM **Huib32*2**, biotin-PEG4-alkyne (negative control), or DMSO (vehicle control) for 1 h. Treated lysates were incubated with NeutrAvidin beads for pulldown. Enriched proteins were processed for immunoblotting, silver staining, or analyzed by LC-MS/MS. **B.** Immunoblot of USP32 protein of DMSO, biotin-PEG4-alkyne, and **Huib32*2**-treated samples. Input samples were analyzed by immunoblotting against USP32 and β-actin. **C, D.** iBAQ values of the enriched proteins via **Huib32*2** were averaged across the three replicates, and the resulting values were used to create the relevant bar graphs. Proteins were filtered (iBAQ value > 0 in all probe-treated samples and unique peptides > 3, and iBAQ value = 0 in both DMSO and Biotin-alkyne control samples) and ranked by iBAQ values. Enzymes related to the Ub system **(C)** and the top-15 highest-ranking proteins **(D)** were then selected for display and Source data are provided in Supplementary Data 8. Corresponding iBAQ values of these proteins, obtained from the total proteomics of MelJuSO cells used for enrichment via **Huib32*2**, were also shown in Supplementary Figure 8, and Source data are provided in Supplementary Data 9.

To extend the analysis beyond ubiquitin-related proteins, we examined the iBAQ values of the top 15 hits that were enriched exclusively in probe-treated samples but not in DMSO or Biotin-PEG4-alkyne controls. While USP32 ranked as the second most enriched protein, the probe also enriched several other proteins. Notably, the most enriched protein, aldehyde dehydrogenase 1 family member A3 (AL1A3), contains an active-site cysteine. Other active-site cysteine-containing enzymes, including fatty acid synthase (FAS), Parkinson’s disease protein 7 (PARK7), and nitrilase family member 2 (NIT2) (Figure 6D) were also identified. Interestingly, PARK7, NIT2 and ALDH1A2 (isoform of AL1A3) were previously identified as off-targets of UCHL1 activity-based probes containing cyanimide warhead^26,29^, suggesting a potential preference for this warhead by these proteins. However, given the significantly higher abundance of these proteins in the MelJuSo proteome compared to USP32 (Supplementary Figure 8B, Supplementary Data 9), these findings still support the strong preference of **Huib32*2** for USP32 over other proteins.

## DISCUSSION

Given the centrality of the ubiquitin-proteasome system in regulating various biological processes, DUBs are increasingly recognized as promising targets for therapeutic intervention, particularly in cancer, neurodegenerative diseases, and immune disorders. Therefore, the discovery of selective inhibitors for deubiquitinases (DUBs) has become a significant area of interest in drug discovery. Despite their importance, the lack of highly specific and potent DUB inhibitors and probes has been a bottleneck in DUB-targeted drug discovery. Recent advancements in the development of specialized libraries,^27,30^ HTS-compatible reagents^31–34^, and ABPs^35,36^ have overcome some of these challenges in the discovery of selective DUB inhibitors, offering valuable tools for research and potential therapeutic leads^37–40^.

Here we have developed the first highly potent and selective inhibitor for USP32, a member of the USP family of DUBs, which is primarily implicated in regulating intracellular trafficking and oncogenic pathways^25^. The development of the USP32 inhibitor, referred to as **Huib32**, was initiated through a high-throughput screening (HTS) campaign using a DUB-targeted, in-house-prepared cyanimide library. **Huib32** emerged as a highly potent inhibitor with a sub-micromolar IC_50_ value, effectively blocking USP32 activity in both in vitro and cellular assays. Remarkably, **Huib32** exhibited high selectivity for USP32 over other closely related DUBs. This specificity is crucial for ensuring that observed cellular effects are due to on-target inhibition of USP32 rather than off-target actions. The cellular effects observed upon **Huib32** treatment are consistent with USP32’s known involvement in cellular trafficking. For instance, **Huib32** induced the dispersion of late endosomes, a hallmark of disrupted endosomal trafficking, and increased the ubiquitination of RAB7, a key substrate of USP32 ^15,16^, and other potential substrates, including RAB6, RAB32, RAB11A/B, and TMEM192. These phenotypic changes provide compelling evidence that USP32 activity directly regulates membrane dynamics and that inhibition of the enzymatic activity of USP32 can disrupt these processes. Therefore, selective inhibition of USP32 may offer a novel therapeutic approach to conditions characterized by defects in cellular trafficking. Specifically, targeting USP32 could hold promise in addressing diseases such as Alzheimer’s and Parkinson’s Diseases, where disruptions in protein degradation and misfolding are central to disease progression.^6^ Building on recent advances in developing small molecule-based DUB-specific ABPs,^26,29,41,42^ in addition to the development of **Huib32**, we further extended its utility through the design of functional probes. However, such a design was not possible for **Huib32**. Therefore, we revisited the screening data from the small library to investigate the structure-activity relationship of Huib32 and its analogs as a starting point to generate a probe precursor (**Huib32\***) containing an azide handle. **Huib32\*** can be rapidly equipped with reporter groups through a click reaction, generating **Huib32*1** and **Huib32*2**, which contain SulfoCy5 and Biotin, respectively. The fluorescent probe **Huib32*1** enables the visualization of USP32 activity in cells, while the biotinylated probe **Huib32*2** facilitates the enrichment of USP32 for quantitative proteomics. Despite a slight reduction in potency, these probes effectively capture USP32 in biochemical and cellular assays, providing valuable tools for mapping USP32-regulated protein networks and investigating its role in cellular trafficking. Furthermore, they offer a foundation for screening and developing selective USP32 inhibitors, expanding their potential applications in therapeutic targeting.

In conclusion, the discovery of **Huib32** as a selective inhibitor of USP32 represents a significant step forward in the field of DUB-targeted drug discovery. The high potency and selectivity of **Huib32** and its ability to modulate USP32 activity in a cellular context make it an invaluable tool for advancing our understanding of USP32’s biological functions and its potential as a therapeutic target. While the development of **Huib32** marks a significant advancement in DUB-targeted drug discovery, several future directions should be explored. Further optimization of **Huib32** and the probes **Huib32*1** and **Huib32*2** that have been discovered in our study could enhance their potency and selectivity, enabling more comprehensive studies of USP32 and its role in cellular processes. Additionally, investigating the therapeutic potential of USP32 inhibition in disease models is a crucial next step. This may lead to the identification of novel treatment strategies for diseases associated with impaired cellular trafficking and ubiquitin signaling.

## MATERIAL AND METHODS

### Echo-mediated high-throughput in-plate synthesis

Echo synthesis was performed as indicated previously.^27^ Briefly, stock solutions of the cyanimide amines in DMSO (200 mM), carboxylic acids in DMSO (100 mM), and a mixture of HOBt and DIC in DMSO (111.1 mM) were prepared in Echo-ready 384PP source plates (Labcyte P-5525). Each well of an Echo-ready 1536LDV plate (Labcyte LP-0400) was charged with the mixture of HOBt and DIC (1125 nL, final concentration 50 mM) using a Thermo Multidrop Combi nL liquid dispenser. Next, the stock solutions of the carboxylic acids (1250 nL, final concentration 50 mM) and the amines (125 nL, final concentration 10 mM) were transferred to the wells using a Labcyte Echo550 acoustic dispenser. The plate was sealed and incubated overnight at room temperature (RT). In total, 7,536 compounds were synthesized as crude reaction mixtures in a total volume of 2.5 µL per well with a maximum concentration of 10 mM, assuming 100% reaction conversion. LC-MS analysis was used to confirm compound formation and purity. A 1 mM daughter plate was prepared in an Echo-ready 1536LDV plate (Labcyte LP-0400) by diluting the compounds 10x in DMSO (300 nL compound solution plus 2700 nL DMSO) using a Labcyte Echo550 acoustic dispenser (compounds) and a Thermo Multidrop Combi nL liquid dispenser (DMSO). Compounds were transferred from the daughter plate into assay plates for HTS.

### Expression and purification of recombinant DUBs and SENPs

Commercially available DUBs used in this study were USP2 (Ubiquigent, #64-0014-050), USP8 (Ubiquigent, #64-0053-050), USP9xCD (Ubiquigent, #64-0017-050), USP10 (Biotechne, E-592), USP30CD (Ubiquigent, #64-0057-050), USP24 (Biotechne, #E-616-050), UCHL5 (Novus biochemicals, #NBP1-72315). In-house purified enzymes included SENP1 (419–644),^27^ USP7 FL, USP15, USP16 FL,^43^ USP32 FL (1-1604), and USP32 CD (735-1604),^15^ UCHL1 FL and UCHL3 FL,^44^ OTUD1 (290-481),^45^ OTUB2 FL,^46^ which were expressed and purified as previously described. mUSP18 FL and hUSP18 FL were produced as described.^47^ For USP6 CD, a new expression construct (pFastNKI-his3C-LIC-USP6-CD) was generated and expressed following a protocol adapted from USP32. In brief, Sf9 cells were cultured in SF900 II SFM media supplemented with 1% penicillin-streptomycin and maintained at 27°C. Cells were transfected with bacmid DNA encoding USP6-CD using Cellfectin II. Baculovirus stocks (P1 and P2) were generated, and protein expression was induced by infecting Sf9 cells with P2 virus. After 72 h, cells were lysed in a buffer containing 50 mM Tris (pH 8.5), 300 mM NaCl, 2 mM TCEP, protease inhibitors, and 0.01% Tween20, followed by sonication and centrifugation. Proteins were purified via Ni2+ affinity chromatography, anion exchange chromatography, and size exclusion chromatography (S200 column). All protein stocks were aliquoted and stored at –80°C

### High-throughput screening

The screen was performed as described previously.^27^ The reaction buffer contained 50 mM Tris-HCl, 100 mM NaCl, 2 mM TCEP, pH 7.5, 1 mg/mL 3-[(3-cholamidopropyl) dimethylammonio] propanesulfonic acid (CHAPS) and 0.5 mg/mL γ-globulins from bovine blood (BGG). The screen was conducted in a 1536-well plate (Corning 3724) with a reaction volume of 8 μL per well. Stock solutions of DUBs and UbRhoMP, ISG15ct-RhoMP or SUMO2RhoMP were prepared. Using a Labcyte Echo550 acoustic dispenser, 10 nL of the 1 mM DMSO stock solutions of the library compounds were transferred from the source plates into the empty 1536-well screening plates to obtain a 1.25 μM final compound concentration. DMSO was used as a negative control and 10 mM of the general DUB inhibitor *N*-Ethylmaleimide (NEM) as a positive control. Next, DUBs (6 μL, the final concentrations of each DUB indicated in Supplementary Data 1) were dispensed using a Biotek MultiflowFX liquid dispenser and incubated for 30 min, followed by dispensing the substrate (2 μL, final concentration 400 nM). After 1-2 h incubation, the fluorescence intensity signal was recorded on a BMG Labtech PHERAstar plate reader (λex/em 480/520 nm). The percentage inhibition of each compound was calculated from the fluorescence intensity values, normalized to the positive (10 mM NEM, 100% inhibition) and negative (DMSO, 0% inhibition) controls.

### IC_50_ determination

The assays were conducted in black 384-well “nonbinding surface flat bottom low flange” plates (Corning 3820) at RT. The reaction buffer contained 50 mM Tris·HCl, 100 mM NaCl (pH 7.6), 1 mg/mL CHAPS, 0.5 mg/mL BGG and 1 mM TCEP. Each well contained a final volume of 20.4 μL. Compounds were dissolved in DMSO as 2, 0.2, and 0.02 mM stock solutions, and appropriate volumes were transferred from these stocks to the empty plate using a Labcyte Echo550 acoustic dispenser and accompanying dose–response software to obtain a 9-point serial dilution (3 replicates) of 3.33 nM to 31.7 µM. A DMSO back-fill was performed to obtain equal volumes of DMSO (400 nL) in each well. NEM (10 mM) was used as a positive control (100% inhibition), and DMSO served as a negative control (0% inhibition). Buffer dispensing steps were performed on a Biotek MultiflowFX liquid dispenser. A total volume of 10 μL of buffer was added, and the plate was vigorously shaken for 20 sec. Next, 5 μL of 3.2 μM recombinant USP32 (4x final concentration) was added and incubated for 30 min. Substrate UbRhoMP (5 μL, final concentration 500 nM) was then added, and the increase in fluorescence over time was recorded using a BMG Labtech PHERAstar plate reader (excitation 487 nm, emission 535 nm). Initial enzyme velocities were determined from the slopes of the fluorescence increase and normalized to the positive and negative controls. IC_50_ values were calculated by plotting normalized values against inhibitor concentrations using the “[inhibitor] vs. response – variable slope (four parameters)” curve fitting method with constraints “Bottom = 0” and “Top = 100” in GraphPad Prism 7 software.

### k_inact_/K_I_ determination

The assays were conducted in black 384-well “nonbinding surface flat bottom low flange” plates (Corning 3820) at RT. The reaction buffer contained 50 mM Tris·HCl, 100 mM NaCl (pH 7.6), 1 mg/mL CHAPS, 0.5 mg/mL bovine γ-globulins (BGG) and 1 mM TCEP. Each well contained a final volume of 20.4 μL. Compounds were dissolved in DMSO as 2, 0.2, 0.02, and 0.002 mM stock solutions, and appropriate volumes were transferred from these stocks to the empty plate using a Labcyte Echo550 acoustic dispenser and accompanying dose–response software to obtain a 10-point serial dilution (3 replicates) of 0.33 nM to 31.7 µM. A DMSO back-fill was performed to obtain equal volumes of DMSO (400 nL) in each well. *N*-Ethylmaleimide (NEM, 10 mM) was used as a positive control (100% inhibition), and DMSO served as a negative control (0% inhibition). Buffer dispensing steps were performed on a Biotek MultiflowFX liquid dispenser. A total volume of 10 μL of buffer was added, and the plate was vigorously shaken for 20 sec. Next, substrate UbRhoMP (5 μL, final concentration 500 nM) was added and incubated for 5 min. Then, 5 μL of 1 μM recombinant USP32 (4x final concentration) was added and the increase in fluorescence over time was recorded using a BMG Labtech PHERAstar plate reader (excitation 487 nm, emission 535 nm). All data fitting and calculations were done using GraphPad Prism 7 software. The fluorescence intensities were plotted against time (in seconds) after a baseline correction using the DMSO control for each inhibitor concentration. The data were fitted to the equation:

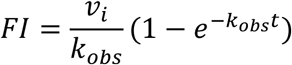

Where FI is the fluorescence intensity (in AU), v_i_ is the initial enzyme velocity (in AU/sec), k_obs_ is the observed rate constant (in sec^-1^), and t is the time (in sec). Thus, obtained k_obs_ values were plotted against the inhibitor concentration ([inh]) and the data were fitted to the equation below to determine the values of k_inact_ and K_I_.

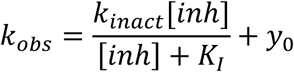

### Jump dilution assay

The assay was performed in a buffer containing 50 mM Tris·HCl, 100 mM NaCl at pH 7.6, 2.0 mM cysteine, 1 mg/mL CHAPS, and 0.5 mg/mL BGG. The DUB activity assay was performed in triplicate with USP32 FL at a final concentration of 0.8 nM, Ub-Rho-morpholine at 500 nM, and **BB01CA282** (**Huib32**) at 250 nM and 2.5 nM, **BB12CA282** at 1 µM and 10 nM, **BB14CA282** at 5 µM and 50 nM, or a jump dilution from highest to the lowest indicated concentrations for each compound. Samples of 20 μL containing 160 nM USP32 and 20 μL containing 500 nM **BB01CA282**, 2 μM **BB12CA282**, 10 μM **BB14CA282** (2% DMSO), 2% DMSO, or 20 mM *N*-

Ethylmaleimide (NEM) were incubated for 30 min at RT. Each sample (3 μL) was then diluted into a 300 μL solution containing 500 nM Ub-Rho-morpholine. After brief mixing, 20 μL of each of these solutions was quickly transferred to a non-binding-surface, flat-bottom, low-flange, black 384-well plate (Corning), and the increase in fluorescence over time was recorded using a BMG Labtech PHERAstar plate reader (excitation 485 nm, emission 520 nm). As a control, samples were taken in which 40 μL of 5 nM and 500 nM of **BB01CA282**, 20 nM and 2 μM of **BB12CA282** or 100 nM and 10 μM of **BB14CA282** solution in buffer (2% DMSO) was added to 35 μL of a 1.83 nM USP32 solution. After 30 min of incubation, 5 μL of 8 μM Ub-Rho-morpholine solution was added, after which 20 μL of each solution was transferred to the same 384-well plate mentioned above, and the increase in fluorescence intensity was measured concomitantly. Fluorescence intensities were plotted against time using GraphPad Prism 7.

### *In vitro* DUB specificity measurement

DUB specificity for **Huib32** at 250 nM was tested by Ubiquigent (DUBprofiler assay) using 100 nM ubiquitin-rhodamine110 as substrate. Additionally, due to the absence of USP32CD and USP32WT in the Ubiquigent panel, they were measured in triplicate by performing the DUB activity assay in-house, together with selected DUBs from the Ubiquigent panel (OTUB2: 25 nM, UCHL1: 1 nM, USP6CD: 0.1 nM, USP7: 0.5 nM, USP16: 2 nM, USP32CD: 0.5 nM, USP32WT 0.4 nM) with **Huib32** at a concentration of 250 nM.

### Cell lines and Cell culture

HEK293T (Cat# ATCC® CRL-3216™) cells were cultured in Dulbecco’s modified Eagle medium (DMEM) (Gibco). MelJuSo cells (human melanoma), kindly provided by Prof. G. Riethmuller (LMU, Munich), were cultured in Iscove’s modified Dulbecco’s medium (IMDM) (Gibco), supplemented with 7.5% fetal bovine serum (FBS) at 37°C and 5% CO_2_. All cell lines have been authenticated and routinely tested for mycoplasma.

### Transfection

For siRNA transfection, sense-siUSP32 GCCUCAGUUACGUGAAUAC and siControl (#Cat) were pursued from Dharmacon. Silencing was performed in MelJuSo as follows: 200 µl siRNA (500 nM stock) were incubated with 4 µl Dharmafect reagent 1 (Dharmacon) diluted in 200 µl medium without supplements with gentle shaking for 20 min at RT. A total of 60 x 10^3^ MelJuSO cells resuspended in 1.6 mL of growth medium without antibiotics from 37.5 x 10^3^ cells per mL suspension were added to transfection mixes to a total volume of 2 mL per well in a 6-well plate and cultured for 3 days prior to further analysis. For DNA transfection, HEK293T cells were seeded into 6-well plate to achieve 50–60% confluence the following day and transfected using PEI (polyethyleneimine, Polysciences Inc., Cat# 23966) as follows: 500 μL DMEM medium without supplements was mixed with 18 μL PEI and 6 μg DNA, incubated at RT for 20 min, and added dropwise to the cells for culturing for 18–24 h prior to further analysis.

### SDS-PAGE and Immunoblotting

Proteins were separated by the indicated percentage of SDS-PAGE for each experiment. After proteins were transferred to a nitrocellulose membrane at 300 mA for 2.5 h, the membranes were blocked with 5% skim milk in PBS and incubated with the indicated primary antibody diluted in 5% skim milk in 0.1% PBS-Tween 20 (PBST) for 1 h at RT. After washing with 0.1% PBST three times for 10 min, proteins were incubated with the indicated secondary antibodies diluted in 0.1% PBST for 30 min and washed three times in 0.1% PBST. The signal was detected using direct imaging by the Odyssey Classic imager (LI-COR).

### DUB Competition assay in cell lysate and living cells

In competition assay using cell lysate, MelJuSo cells were suspended in the cell lysis buffer (50 mM Tris,150 mM NaCl, 0.5% Triton X-100, and 2 mM TCEP at pH 7.5), supplemented with protease inhibitor cocktail (11836145001, Roche). The samples were kept on ice and sonicated (Fisher Scientific FB120 Sonic Dismembrator, 3 pulses, amplitude 40%). The cell lysate was centrifuged at 21,500 g with Eppendorf Centrifuge 5430 R for 20 min at 4°C, and supernatant fractions were collected. Cell lysates were incubated with the indicated concentrations of **Huib32** for 1 h, followed by the addition of Rho-Ub-PA and incubation for 5 min at 37°C. Competition assay in living cells was performed in MelJuSo or HEK293T cells, in which they were treated with **Huib32** at specified concentrations and time points prior to lysis. Cell lysates were incubated with 0.5 µM final concentration of Rho-Ub-PA for 5 min at 37°C. for both assays, For both assays, reactions were stopped by boiling in NuPAGE LDS sample buffer containing β-mercaptoethanol and resolved by SDS-PAGE (4-12% Bis-Tris gel, MOPS SDS running buffer). Gels were visualized by fluorescence scanning (Typhoon FLA 9500, Rhodamine channel, λex/em 473/530 nm) and proteins were transferred to nitrocellulose membranes. Immunoblotting was performed with mouse anti-USP32 (Santa Cruz Biotechnology, Cat# sc-374465; 1:1000) and anti-β-actin (Sigma-Aldrich, Cat# A544; 1:10000) primary antibodies, followed by IRDye 800CW secondary antibodies (Li-COR, Cat# 926-32210; 1:5000).

### In-cell target engagement and selectivity of Huib32 via RhoK(Biotin)UbPA probe

MelJuSo cells were seeded into 10 cm dishes to achieve 50-60% confluence by the following day. Cells were then treated with either 10 µM **Huib32** or an equivalent volume of DMSO as a control and incubated for 24 h. The experiment was performed in triplicate, with one 10 cm dish prepared per sample. Following treatment, cells from each dish were suspended in 300 µL of lysis buffer (50 mM Tris, 150 mM NaCl, 0.5% Triton X-100, and 2 mM TCEP, pH 7.5) supplemented with a protease inhibitor cocktail (Roche, 11836145001). Samples were kept on ice, sonicated using a Fisher Scientific FB120 Sonic Dismembrator (3 pulses, amplitude 40%), and centrifuged at 21,500 x g for 20 min at 4°C using an Eppendorf Centrifuge 5430 R. The resulting supernatants were collected as cell lysates. Cell lysates were incubated with a final concentration of 1 µM RhoK(Biotin)UbPA probe for 5 min at 37°C. The reactions were stopped by diluting the lysates to a final volume of 1 mL with lysis buffer containing 1% SDS. The samples were then incubated with NeutrAvidin beads (Thermo Fisher, Cat# 29204) overnight at 4°C. The beads were washed five times with lysis buffer containing 1% SDS to remove nonspecifically bound proteins. Subsequently, the bound proteins were eluted by boiling the beads in NuPAGE LDS sample buffer supplemented with β-mercaptoethanol. Eluted proteins were resolved by SDS-PAGE using a 4-12% Bis-Tris gel in MOPS SDS running buffer for fluorescence scanning. For LC/MSMS analysis eluted proteins were separated for a migration distance of 2 cm, followed by staining of the gel with Instant Blue Coomassie protein stain (Expedeon). The stained protein bands were excised and submitted for LC-MS/MS analysis to identify and quantify the proteins. See the Mass Spectrometry section below for details of mass spectrometry analysis

### Confocal microscopy

MelJuso cells (7.5 x 10³) were seeded into 24-well plates (Costar, Cat# 3524) containing glass coverslips (Menzel Gläser, Cat# MENZCB00130RAC) and, the next day, treated with the indicated final concentration of **Huib32** or DMSO (0 µM) for 72 h. Following incubation, cells were washed with Phosphate Buffer Saline (PBS), fixed with 3.7% formaldehyde (acid-free, Merck Millipore) in PBS for 20 min, and permeabilized with 0.1% Triton X-100 (Sigma-Aldrich, Cat# T8787) in PBS for 10 min. After blocking with 5% skim milk powder (Oxoid, Cat# LP0031) in PBS for 30 min, cells were incubated with anti-MCHII antibody^48^ (1:300 dilution) for 1 hour, washed in PBS (three times for 5 min), and subsequently incubated with anti-rabbit Alexa-dye-coupled secondary antibodies (Invitrogen) for 30 min. After further washing in PBS (three times for 5 min), cells were mounted with ProLong Gold antifade Mounting medium with DAPI (Life Technologies, Cat# P36941) and imaged using a Leica SP8 microscope equipped with solid-state lasers, HCX PL 63x oil immersion objectives, and HyD detectors. Data were collected with digital zoom (2.5x) in a 1024 x 1024 scanning format, followed by post-processing in Fiji software.

### Ubiquitination assays

The ubiquitination status of GFP-tagged proteins was assessed by using the ubiquitination assay described previously.^15^ HEK293T cells and transfected with HA-ubiquitin, GFP-RAB7 WT, or GFP-RAB7 2KR or MelJuSo cells stably expressing GFP-RAB7 WT incubated with **Huib32** or DMSO were harvested in 300 µL lysis buffer containing 50 mM Tris-HCl, pH 7.5, 150 mM NaCl, 0.5% Triton X-100, 10 mM N-methyl maleimide (NMM) (general DUB inhibitor diluted in DMSO, freshly added) and protease inhibitor cocktail (Roche Diagnostics, EDTA-free, freshly added) by scraping. Then, 100 µL buffer containing 100 mM Tris-HCl, pH 8.0, 1 mM EDTA, and 2% SDS was added to the crude lysates; samples were sonicated (Fisher Scientific FB120 Sonic Dismembrator, 3 pulses, amplitude 40%) followed by bringing sample volume to 1 mL with lysis buffer 1. After centrifugation (20 min, 4°C, 20,817× g), lysates were incubated with 6 µL GFP_Trap_A beads (Chromotek) overnight at 4°C. Beads were washed four times with lysis buffer and all liquid was removed before the addition of SDS sample buffer (containing 10 mM DTT). Proteins were denatured by heating at 95°C for 15 min, subjected to 8% SDS-PAGE, and detected by immunoblotting, as indicated above, using mouse anti-ubiquitin (Cat# sc-8017, 1:1000), mouse anti-HA (HA.11 (16B12), Covance, Cat# MMS-101R; 1:1000), rabbit anti-mGFP^49^ (1:1000) primary antibodies and IRDye 680LT goat anti-rabbit IgG (H + L) (Li-COR, Cat# 926-68021, 1:20,000) and IRDye 800CW goat anti-mouse IgG (H + L) (Li-COR, Cat# 926-32210, 1:5000) secondary antibodies.

### Identification of USP32 enzymatic activity-dependent ubiquitome and total proteome

MelJuSo cells were seeded into 10-cm plates in 10 mL of medium. A day after seeding, cells were incubated with 10 µM **Huib32** or DMSO for 4 or 72 h. The cells were washed twice with cold PBS (1 mL per 10-cm plate) and collected by scraping them with a cell scraper. GlyGly-containing peptide immunoprecipitation was performed on MelJuSo cell lysates using the PTMScan Ubiquitin Remnant Motif Kit (Cell Signaling #5562), according to the manufacturer’s protocol as described previously. Briefly, 3 mg of extracts were solubilised and denatured in 10 mL of lysis buffer (20 mM HEPES, pH 8.0, 9 M urea, 1 mM sodium orthovanadate, 2.5 mM sodium pyrophosphate, 1 mM β-glycerophosphate), reduced using dithiothreitol (4.5 mM final concentration) for 30 min at 55 °C. This was followed by alkylation using iodoacetamide (100 mM final concentration) for 15 min at RT in the dark. Samples were subsequently diluted fourfold in 20 mM HEPES, pH 8.0 (∼2 M urea final), followed by digestion with trypsin-TPCK (Worthington, LS003744, 10 mg/mL final) overnight at RT. Peptide samples were then acidified using trifluoroacetic acid (1% final concentration) and desalted using C-18 SepPak cartridges (Waters) according to the manufacturer’s protocol. At this point, 20 µg of digested protein was removed from each sample for matching total proteome control analysis. Peptides were lyophilised and resuspended in 1.4 mL immunoprecipitation IAP buffer (PTMScan), and the remaining insoluble material cleared by centrifugation. Anti-GlyGly antibody beads were then added to the solution, which was then rotated end-over-end for 2 h at 4 °C. Beads were subsequently washed twice using 1 mL IAP buffer, followed by three water washes. Immunoprecipitated material was eluted twice in 55 and 50 μL 0.15% trifluoroacetic acid in water. Purified GlyGly-modified peptide eluates and matching proteome material were dried by vacuum centrifugation and re-suspended in buffer A (98 % MilliQH20, 2% CH3CN and 0.1% TFA). LC-MS/MS analysis was performed using a Dionex Ultimate 3000 nano-ultra high-pressure reverse-phase chromatography coupled online to a Fusion Lumos as described previously,^40,50^ using a data-independent acquisition (DIA) method.^51^

MS raw files were searched against the UniProtKB human sequence database, and label-free quantitation was performed using DIA-NN Software (v1.8) in library-free mode. Search parameters include carbamidomethyl (C) as a fixed modification, and oxidation (M), acetylation (N-term), and K-GG (for GG-peptidome) as variable modifications. A maximum of two missed cleavages were allowed, and the matching between runs (MBR) function was enabled. Label-free interaction data analysis was performed in Perseus (v1.6.2.3), and volcano and scatter plots were generated using a t-test with a permutation FDR = 0.01 for multiple-test correction and s0 = 0.1 as cut-off parameters.

### Labeling of purified recombinant USP32 with Huib32*1 and Rho-Ub-PA probes

The assay was conducted in Tris buffer (50 mM Tris-HCl, 150 mM NaCl, 2 mM TCEP, pH 7.5). Recombinant human USP32 FL protein (4 μM final concentration) was incubated with 10 μM final concentration of **Huib32*1** or Rho-Ub-PA probes for 1 h at 37°C. After the incubation, all of the reactions were stopped by adding NuPAGE LDS sample buffer containing β-mercaptoethanol or TCEP. Samples containing β-mercaptoethanol were heated at 95°C for 5 min. Proteins were then separated by SDS-PAGE using 4-12% Bis-Tris precast gels (Invitrogen, NuPAGE) with MOPS SDS running buffer (Novex, NuPAGE). The gels were visualized using a Typhoon FLA 9500 (GE Healthcare Life Sciences) fluorescence scanner, where USP32 adducts were detected in the Cy5 channel (λex/em 635/655 nm) or Rhodamine channel (λex/em 473/530 nm). Gels were subsequently stained with Instant Blue Coomassie protein stain (Expedeon) and scanned on an Amersham Imager 600 (GE Healthcare Life Sciences) for further analysis.

### Probe labeling of endogenous and ectopically expressed USP32 in cells

For endogenous USP32 labeling, MelJuSo cells were transfected with siControl or siUSP32 in a 6-well plate for 72 h. the day before the harvesting, the cells were incubated with 10 μM final concentration of **Huib32*1** probe for 24 h, followed by cell lysis in the buffer containing 50 mM Tris-HCl, pH 7.5, 150 mM NaCl, 0.5% Triton X-100, and protease inhibitor cocktail (Roche Diagnostics, EDTA-free, freshly added) by sonication (Fisher Scientific FB120 Sonic Dismembrator, 3 pulses, amplitude 40%) and adding loading buffer (4x). Proteins were then separated by SDS-PAGE using 4-12% Bis-Tris precast gels (Invitrogen, NuPAGE) with MOPS SDS running buffer (Novex, NuPAGE). For labeling of ectopically expressed USP32 WT and USP32 CA mutant, HEK293T cells were seeded into a 6-well plate the day before transfection of USP32 constructs. Cells were transfected with 2xHA-N1 empty vector, USP32-HA WT or USP32-HA C743A mutant using PEI transfection reagent (polyethylenimine, Polysciences Inc., Cat# 23966) as follows: 250 μL DMEM medium without supplements was mixed with 18 μL PEI and 6 μg DNA, incubated at RT for 20 min, and added drop-wise to the cells and **Huib32*1** probe added to cells 4 h after transfection for culturing for 24 h before further analysis. Samples were resolved by SDS-PAGE using a 4-12% Bis-Tris gel with MOPS SDS running buffer. The gels were visualized using a Typhoon FLA 9500 (GE Healthcare Life Sciences) fluorescence scanner, where USP32 adducts were detected in the Cy5 channel (λex/em 635/655 nm), followed by protein transfer to nitrocellulose membrane. Immunoblot analysis was performed as described above by using mouse anti-USP32 antibody (Santa Cruz Biotechnology, Cat# sc-374465; 1:1000), mouse anti-β-actin (Sigma-Aldrich, Cat# A544; 1:10000) primary antibodies and IRDye 800CW goat anti-mouse IgG (H + L) (Li-COR, Cat# 926-32210, 1:5000) secondary antibody.

### Protein enrichment with biotinylated Huib32*2 probe

MelJuSo cells were seeded into 10 cm dishes and collected when they reached to 80-90% confluency. Cells from each dish were suspended in 300 µL of lysis buffer (50 mM Tris, 150 mM NaCl, 0.5% Triton X-100, and 2 mM TCEP, pH 7.5) supplemented with a protease inhibitor cocktail (Roche, Cat# 11836145001). The experiment was performed in triplicate, with one 10 cm dish prepared per sample. Samples were kept on ice, sonicated using a Fisher Scientific FB120 Sonic Dismembrator (3 pulses, amplitude 40%), and centrifuged at 21,500 x g for 20 min at 4°C using an Eppendorf Centrifuge 5430 R. The resulting supernatants were collected as cell lysates. Cell lysates were incubated with a final concentration of 1 µM **Huib32*2** probe or negative controls (Biotin (PEG4) alkyne or DMSO) for 1 h at 37°C. The reactions were diluted to a final volume of 1 mL with lysis buffer. The samples were then incubated with NeutrAvidin beads (Thermo Fisher, Cat# 29204) overnight at 4°C. The beads were washed five times with lysis buffer containing 1% SDS to remove nonspecifically bound proteins. Subsequently, the bound proteins were eluted by boiling the beads in NuPAGE LDS sample buffer supplemented with β-mercaptoethanol. Eluted proteins were resolved by SDS-PAGE using a 4-12% Bis-Tris gel in MOPS SDS running buffer, with proteins separated for a migration distance of 2 cm. The gel was stained with Instant Blue Coomassie protein stain (Expedeon). The stained protein bands were excised and submitted for LC-MS/MS analysis to identify and quantify the proteins. Separately, untreated MelJuSo cells were also analyzed by LC-MS/MS for total proteomics profiling. See the Mass Spectrometry section below for details of mass spectrometry analysis

### Mass Spectrometry

For liquid chromatography tandem mass spectrometry (LC-MS/MS), gel slices were washed, subjected to reduction with dithiothreitol, alkylation with iodoacetamide and in-gel trypsin digestion using a Proteineer DP digestion robot (Bruker). Tryptic peptides were extracted from the gel slices and lyophilized.

Peptides from gel bands were dissolved in 95/3/0.1 v/v/v water/acetonitril/formic acid and analyzed by on-line C18 nanoHPLC MS/MS with a system consisting of an Easy nLC 1000 gradient HPLC system (Thermo, Bremen, Germany), and a LUMOS mass spectrometer (Thermo). Samples were injected onto a homemade precolumn (100 μm × 15 mm; Reprosil-Pur C18-AQ 3 μm, Dr. Maisch, Ammerbuch, Germany) and eluted via a homemade analytical nano-HPLC column (30 cm × 50 μm; Reprosil-Pur C18-AQ 3 μm). The gradient was run from 10% to 40% solvent B (20/80/0.1 water/acetonitrile/formic acid (FA) v/v) in 30 min. The nano-HPLC column was drawn to a tip of ∼10 μm and acted as the electrospray needle of the MS source. The LUMOS mass spectrometer was operated in data-dependent MS/MS mode for a cycle time of 3 seconds, with a HCD collision energy at 30% and recording of the MS2 spectrum in the orbitrap. In the master scan (MS1) the resolution was 120,000, the scan range 400-1500, at an AGC target of ‘standard’. A lock mass correction on the background ion m/z=445.12003 was used. Dynamic exclusion after n=1 with exclusion duration of 10 s. Charge states 2-5 were included. For MS2 precursors were isolated with the quadrupole with an isolation width of 1.2 Da. The MS scan range was set to ‘auto’. The MS2 scan resolution was 30,000 with an AGC target of ‘standard’ with a maximum fill time of ‘auto’.

Alternatively, peptide fractions were dissolved in water/formic acid (100/0.1 v/v) and analyzed by on-line C18 nanoHPLC MS/MS with a system consisting of an Ultimate3000nano gradient HPLC system (Thermo, Bremen, Germany), and an Exploris480 mass spectrometer (Thermo). Fractions were injected onto a cartridge precolumn (300 μm × 5 mm, C18 PepMap, 5 μm, 100 A, and eluted via a homemade analytical nano-HPLC column (50 cm × 75 μm; Reprosil-Pur C18-AQ 1.9 μm, 120 A (Dr. Maisch, Ammerbuch, Germany). The gradient was run from 2% to 36% solvent B (20/80/0.1 water/acetonitrile/formic acid (FA) v/v) in 120 min at 250 nL/min. The nano-HPLC column was drawn to a tip of ∼10 μm and acted as the electrospray needle of the MS source. The mass spectrometer was operated in data-dependent MS/MS mode with a cycle time of 3 s, with a HCD collision energy at 30% and recording of the MS2 spectrum in the orbitrap, with a quadrupole isolation width of 1.2 Da. In the master scan (MS1) the resolution was 120,000, the scan range of *m/z* 400-1500, at standard AGC target and a maximum fill time of 50 ms. A lock mass correction on the background ion m/z=445.12003 was used. Precursors were dynamically excluded after n=1 with an exclusion duration of 10 sec, and with a precursor range of 30 ppm. Included charge states were 2-5. For MS2 the first mass was set to 120 Da, and the MS2 scan resolution was 30,000 at an AGC target of 75% at a maximum fill time of 60 ms.

RAW data were processed using MaxQuant (v 2.5.1) and mapped to the minimal Homo sapiens UniProt proteome database (20,596 entries). For all analyses, the default Maxquant settings were used, with trypsin as the enzyme. Additionally, the iBAQ^52^ was selected in the global parameters tab.

## ASSOCIATED CONTENT

### Supporting Information

Supporting Information containing table and figures, synthesis schemes, NMR and LC-MS spectral data is provided as a separate file (PDF)

Supplementary Data 1–9, including IC₅₀ values, high-throughput screening results, compound SMILES, and proteomics data analyses, are available upon request.

## ACKNOWLEDGEMENTS

This project is funded by the Institute for Chemical Immunology (grant no. ICI00026 to A.S.) and the Innovative Medicines Initiative 2 (IMI2) Joint Undertaking under grant agreement no. 875510 (EUbOPEN project). A.P.F. and B.M.K. were supported by the Chinese Academy of Medical Sciences (CAMS) Innovation Fund for Medical Science (CIFMS), China [grant number: 2018-I2M-2-002] and by Pfizer. Work in the A.P.F. lab was also funded by Boehringer Ingelheim and Ono Pharma UK Ltd.

## AUTHOR CONTRIBUTIONS

S.S. and A.S. designed the research. P.P.G. and R.K. conducted the initial high-throughput screen. V.P. synthesized the chemical compounds identified in the first screen and Huib32*. S.S. synthesized the Huib32*1 and Huib32*2 probes. P.P.G. performed kinetics experiments. D.S.H. designed and synthesized the RhoK(Biotin)UbPA probe. A.P.F., B.M.K., D.O.B., and I.V. performed the mass spectrometry analysis for ubiquitomics and total proteomics data. P.A.V. and R.T.N.T. performed mass spectrometry analysis of competitive DUB selectivity pulldown, Huib32*2 pulldown, and Meljuso cells total proteomics data. S.S., P.P.G., and A.S. interpreted the data and wrote the manuscript.

---

**Scheme 1.**
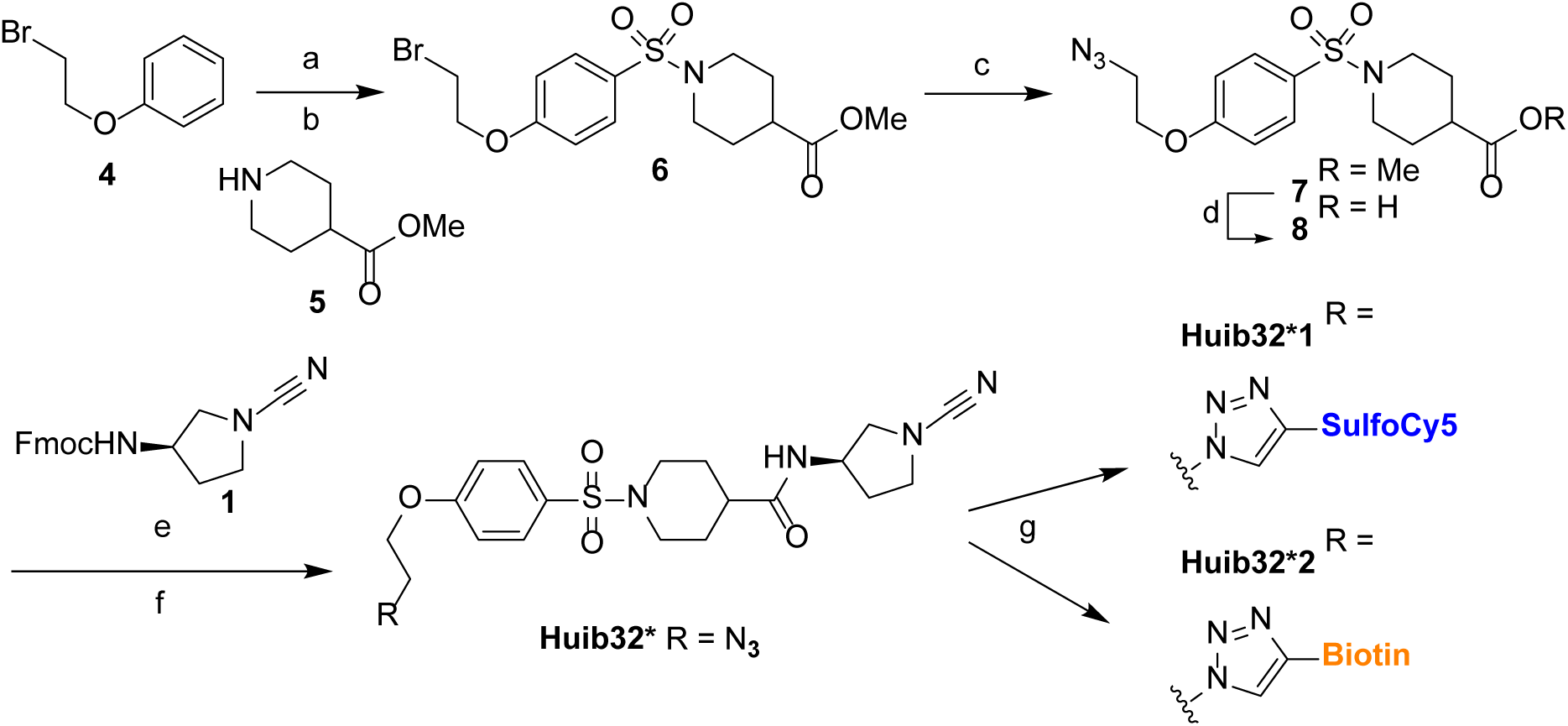
Synthesis of Huib32*1 and Huib32*2. (a) HSO_3_Cl, DCM, −20°C; (b) Et_3_N, 0°C to rt; (c) NaN_3_, DMF, 80°C; (d) LiOH, THF; (e) DBU, DMF; (f) EDC*HCl, DiPEA; (g) Biotin-alkyne or SulfoCy5-alkyne, CuSO_4_, sodium ascorbate, tris((1-benzyl-4-triazolyl)methyl)amine (TBTA) ester, DMSO/H_2_O/ACN.

