## Supplementary Information for "Huib32: A Potent and Selective USP32 Inhibitor Modulating Endosomal Processes and Advancing Cell-Permeable USP32 Probes"

Dedication: In memory of Huib Ovaa

NMR & LC-MS analysis of synthesized compounds…………………………………...19

Supplementary Figure 1

A

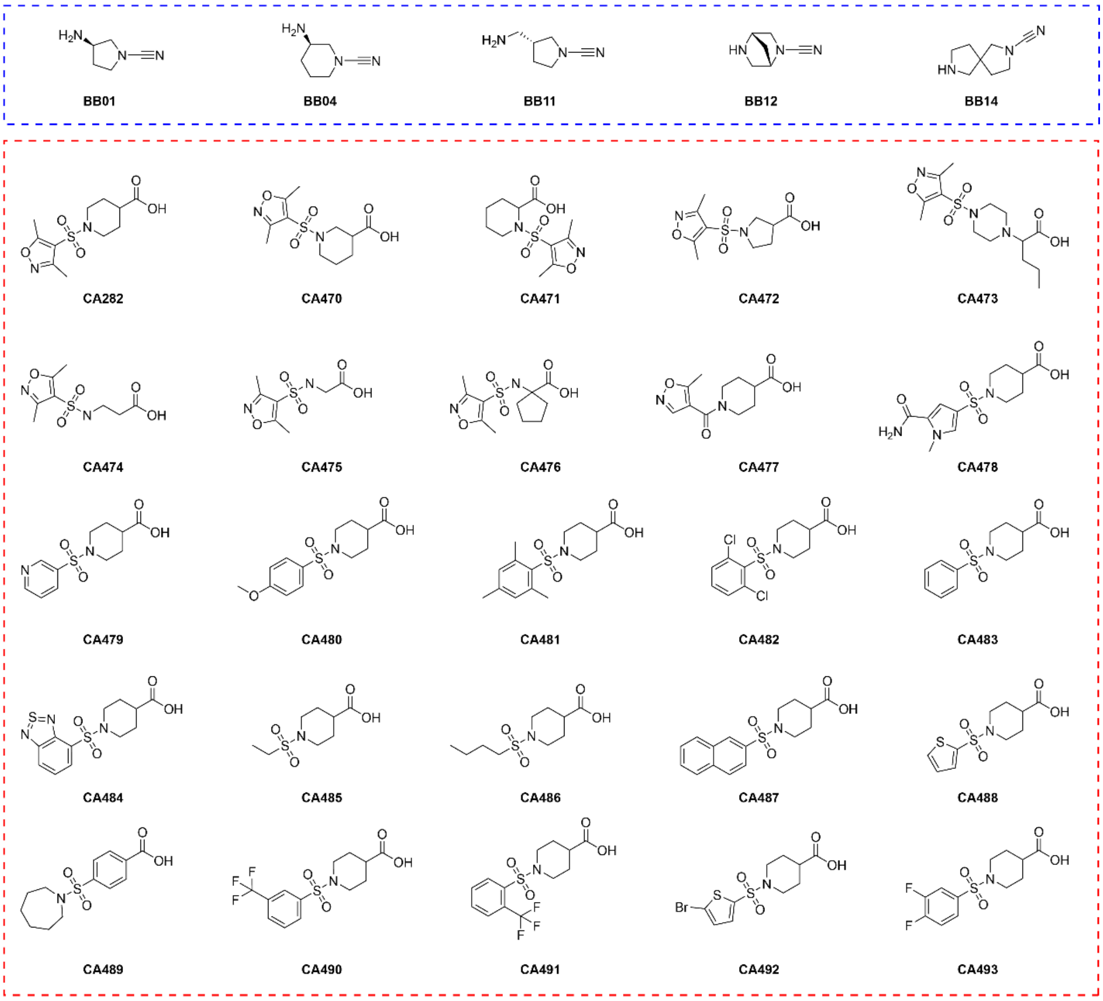

B

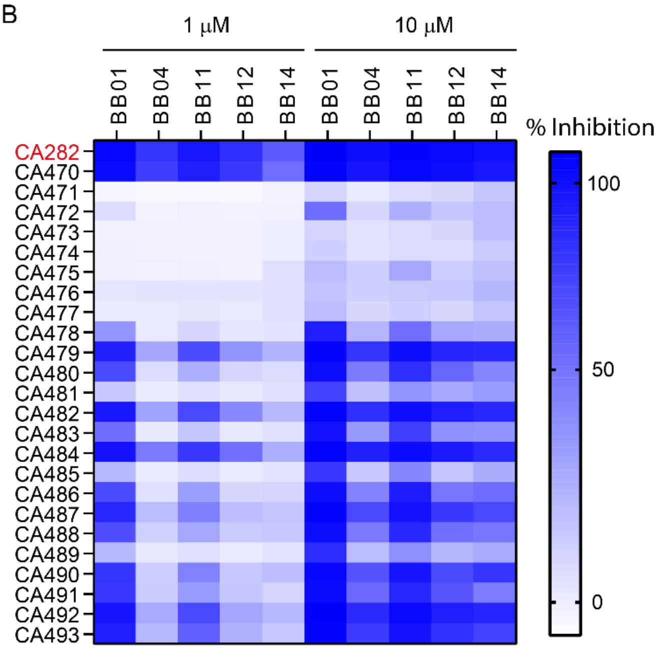

**Supplementary Figure 1. Screening of secondary library containing 125 compounds. A.** Overview of the 5 amine building blocks (BB) and 25 carboxylic acids (CA) used to create a small library containing 125 compounds designed based on carboxylic acid **CA282**. **B.** Heatmap displaying the USP32 screening results of the 125 compound library at 1 and 10 µM compound concentration using UbRhoMP-based fluorescent intensity assay. Related to Figure 1.

Supplementary Figure 2

A

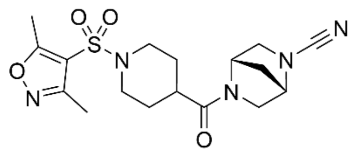

BB012CA282

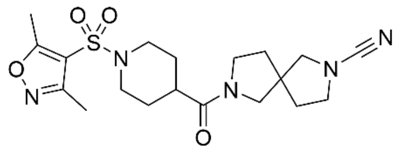

BB014CA282

B

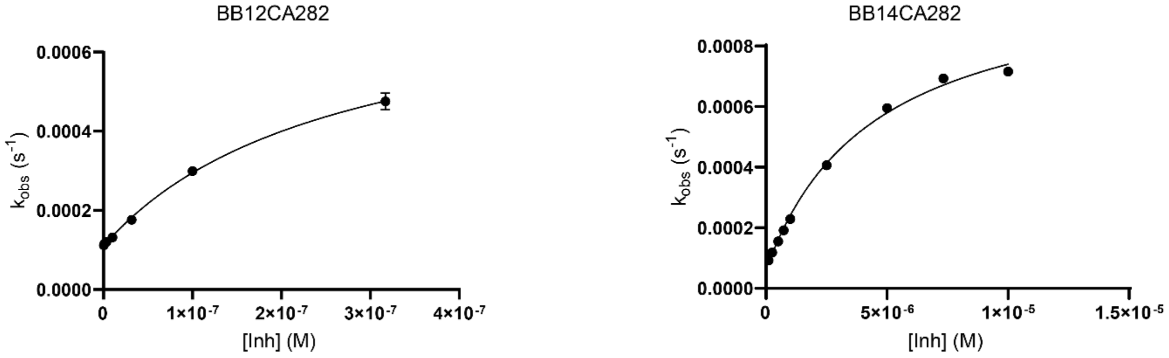

C

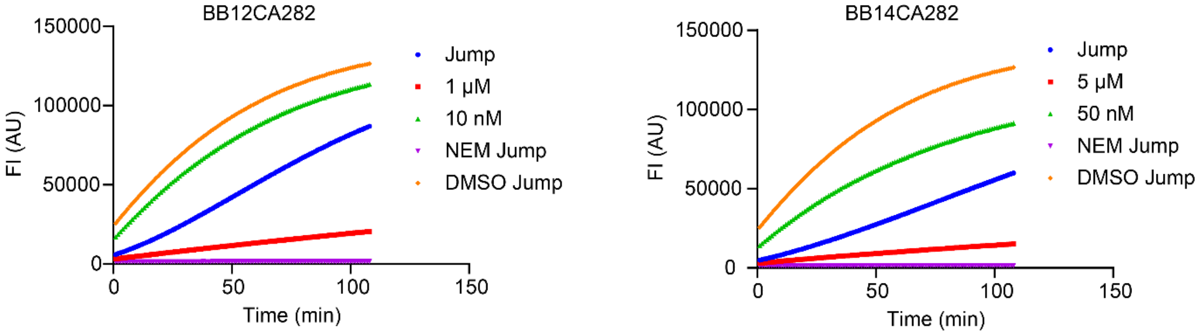

**Supplementary Figure 2. Properties of USP32 inhibitors A.** Structures of BB012CA282 and BB014CA282 compounds. **B.** Kinetics plot of $k_{obs}$ versus different concentrations of BB012CA282 and BB014CA282 compounds. **C.** Jump dilution progress curve for USP32 proteolytic activity for BB012CA282 and BB014CA282 compounds. Related to Figure 2

**Table S1.** $IC_{50}$, $k_{inact}$, $K_i$, and $k_{inact}/K_i$ values

| Compounds | $IC_{50}$ (nM) | $k_{inact}$ ($s^{-1}$) | $K_i$ (M) | $k_{inact}/K_i$ ($s^{-1}M^{-1}$) |
|---|---|---|---|---|
| **BB01CA282** | 21.2 | 0.000222 | 1.18E-08 | 18,800 |
| **BB04CA282** | 19837 | n.d. | n.d. | n.d. |
| **BB11CA282** | 1937 | n.d. | n.d. | n.d. |
| **BB12CA282** | 94 | 0.000324 | 1.52E-07 | 2,130 |
| **BB14CA282** | 518.1 | 0.000984 | 4.49E-06 | 219 |

n.d.: Not determined.

Supplementary Figure 3

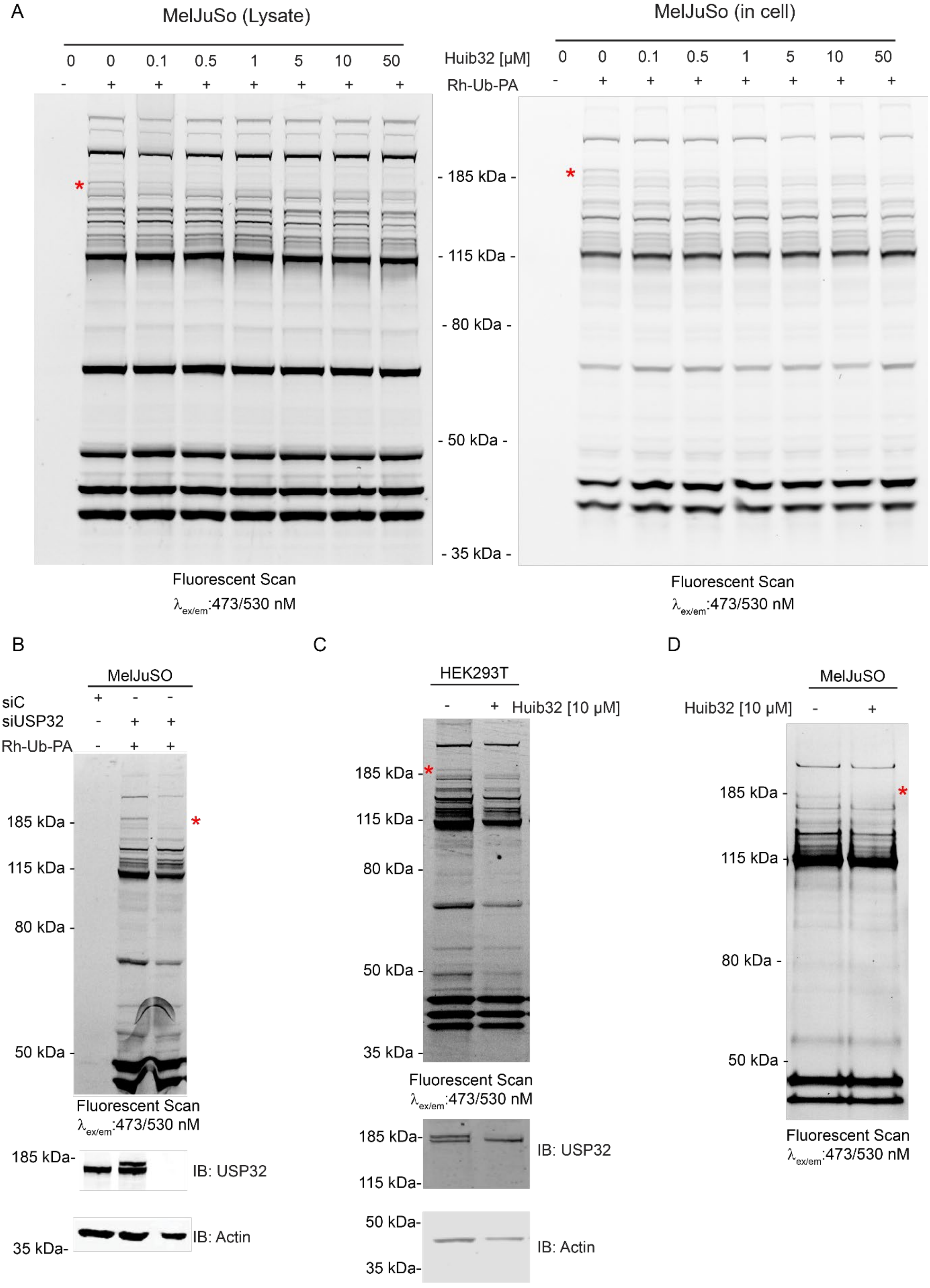

**Supplementary Figure 3. *In vitro* and cellular specificity of USP32 inhibitors. A.** Cell lysates (left panel) were incubated with the indicated concentration of Huib32 for 1 h, while intact MelJuSo cells (right panel) were incubated with Huib32 for 24 hours. After these incubations, both cell lysates were treated with the Rho-Ub-PA probe for 5 min. Samples were resolved by SDS-PAGE and gels were analyzed by fluorescence scanning. See also Figure 3B. **B.** Determination of the band corresponding to the endogenous USP32 in DUB activity-based probe labeling assay. USP32 was depleted in MelJuSo cells by transfection with siControl or siUSP32, followed by cell lysis and incubation with Rho-Ub-PA probe for 5 min. The samples were subjected to SDS-PAGE, analyzed by fluorescence scanning and immunoblotted against USP32 and β-actin. β-Actin was used as a loading control. **C.** Target engagement of **Huib32** in HEK293T cells. HEK293T cells were treated with DMSO or 10 μM final concentration of **Huib32** for 24 h, followed by cell lysis and incubation with Rho-Ub-PA probe for 5 min. The samples were subjected to SDS-PAGE, analyzed by fluorescence scanning, and immunoblotted against USP32 and β-actin. β-Actin was used as a loading control. **D.** MelJuSo cells were treated with 10 μM **Huib32** for 24 hours, lysed, and incubated with a Rho-K(Biotin)-Ub-PA probe for 5 minutes. DUBs modified by the probe were pulled down using NeutrAvidin beads, and the samples were analyzed by fluorescence scanning. See also Figures 3C and 3D. Supplementary Figure 3 is related to Figure 3.

Supplementary Figure 4

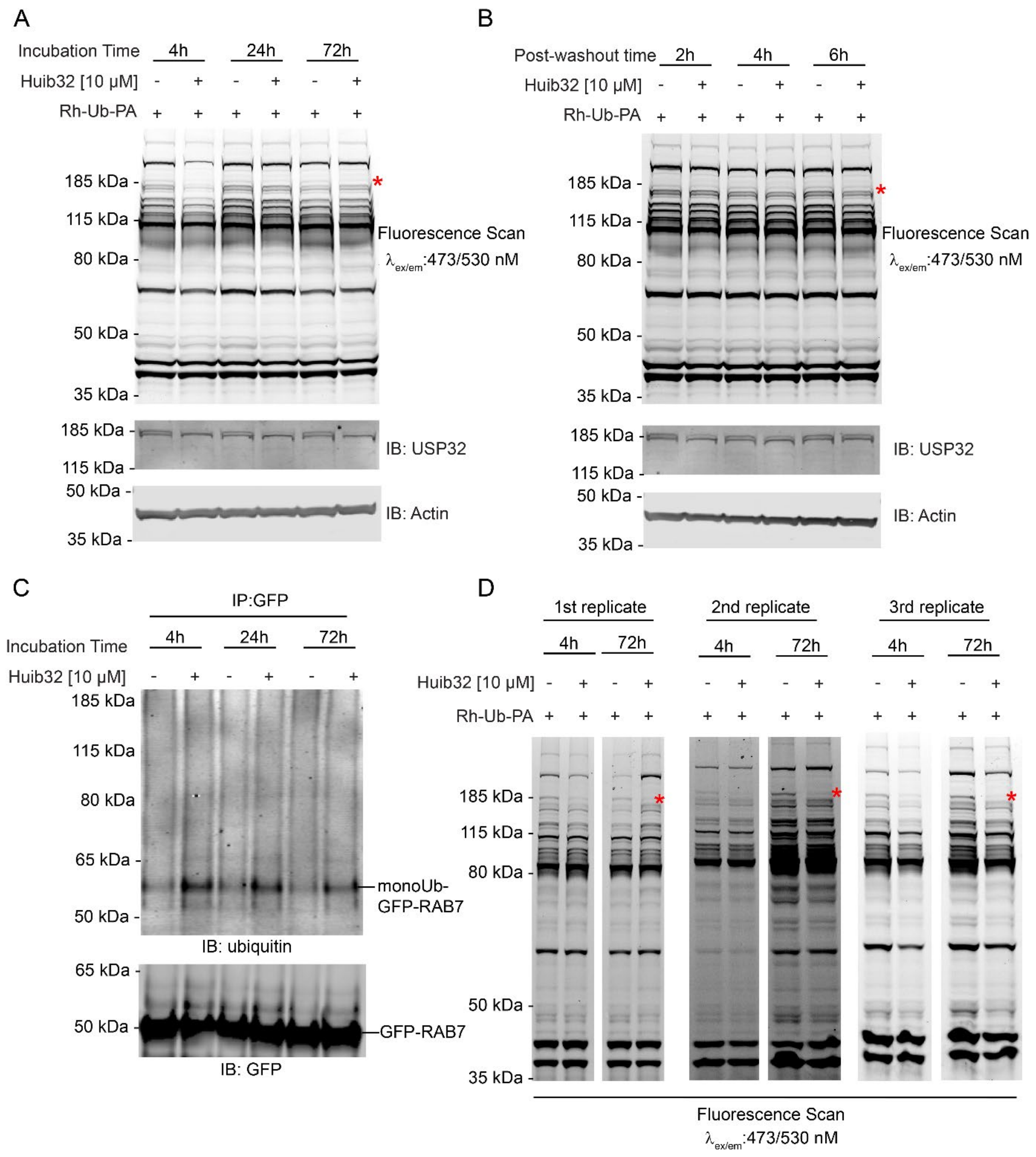

**Supplementary Figure 4. Assessment of the stability of Huib32 in cells and optimization for ubiquitomics and proteomics analysis. A.** Inhibition of USP32 by **Huib32** at various time points. MelJuSo cells were treated with 10 µM **Huib32** for the indicated times, followed by cell lysis and a 5-minute incubation with the Rho-Ub-PA probe. Samples were resolved by SDS-PAGE, scanned for fluorescence, and immunoblotted against USP32 and β-actin, with β-actin serving as a loading control. **B.** Time-dependent recovery of cellular USP32 activity following **Huib32** washout.

MelJuSo cells were incubated with 10 μM **Huib32** for 24 h. After washing out **Huib32**, cells were allowed to recover for the indicated times (post-washout). Cell lysates were then incubated with Rho-Ub-PA for 5 min. Samples were resolved by SDS-PAGE, scanned for fluorescence, and immunoblotted against USP32 and β-actin (loading control). **C.** Ubiquitination assay of GFP-RAB7 under **Huib32** treatment. MelJuSo cells stably expressing GFP-RAB7 were treated with 10 μM **Huib32** for the indicated times, followed by immunoprecipitation with GFPTrap beads and analysis by immunoblotting with anti-ubiquitin and anti-GFP antibodies. **D.** Rho-Ub-PA probe labeling and fluorescence analysis of replicates prepared for the ubiquitomics experiment in Figures 4C and 4D. Cell lysates from three replicates of MelJuSo cells treated with 10 μM **Huib32** for 4 or 24 h were incubated with the Rho-Ub-PA probe for 5 min. Samples were analyzed by SDS-PAGE and fluorescence scanning. Related to Figure 4.

Supplementary Figure 5

A

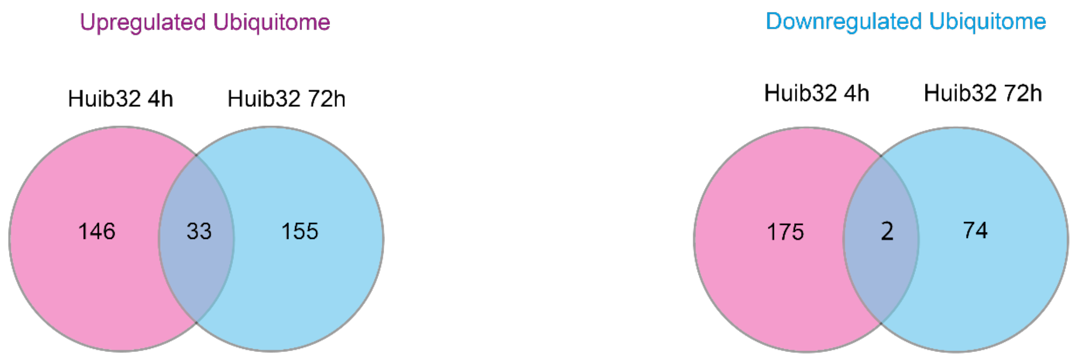

B

**Upregulated**

| | Protein ID | Protein Name | Peptide sequence (GG-K) Huib32-4h and 72h |
|---|---|---|---|
| 1 | O43934 | MFSD11 | AVDAFK(228)K |
| 2 | O75390 | CS | SMSTEGLMK(459)FVDSK |
| 3 | P06576 | ATP5F1B | VVDLLAPYAK(198)GGK_GGK(201)IGLFGGAGVGK |
| 4 | P08708 | RPS17 | EMLK(107)LLDFGSLSNLQVTQPTVGMNFK |
| 5 | P20340 | RAB6A | AK(146)ELNVMFIETSAK |
| 6 | P38646 | HSPA9 | NVPFK(143)IVR |
| 7 | P51809 | VAMP7 | LELLIDK(160)TENLVDSSVTFK_TENLVDSSVTFK(171)TTSR |
| 8 | Q8WVX9 | FAR1 | K(443)YVLNEEMSGLPAAR |
| 9 | P61106 | RAB14 | K(35)FMADCPHTIGVEFGTR |
| 10 | P61421 | ATPV0D1 | AK(343)IDNYIPIF |
| 11 | Q12893 | TMEM115 | QLALK(289)ALNER |
| 12 | Q13637 | RAB32 | IK(210)LDQETLR |
| 13 | Q587I9 | SFT2D3 | GAGLAK(211)VLPV |
| 14 | Q6DKI1 | RPL7L1 | LEVK(84)PHALELPDK |
| 15 | Q8IY95 | TMEM192 | QGDTIEYLK(246)R |
| 16 | Q8N697 | SLC15A4 | QSNGEGIGVFQQSSK(292)QSLFDSCK |
| 17 | Q8WV19 | SFT2D1 | DAVIK(152)CCSSLLS |
| 18 | Q93009 | USP7 | DLLQFFK(869PR |
| 19 | Q96BM9 | ARL8A | DLPGALDEK(141)ELIEK |
| 20 | Q96K49 | TMEM87B | MFSSEK(553)IM |
| 21 | Q96K76 | USP47 | TLK(782)AEGFFR |
| 22 | Q9BXW6 | OSBL1A | NFK(432)LEQEQEK |
| 23 | Q9BZQ8 | NIBAN1 | TAVAIEK(475)VK |
| 24 | Q9H6Y7 | RNF167 | LTK(211)EQLK_QIPTHDYQK(223)GDQYDVCAICLDEYEDGDK |
| 25 | Q9HD26 | GOPC | LYLDELEGGGNPGASCK(409)DTSGEIK |
| 26 | Q9NRX5 | SERINC1 | TSNNSQVNK(343)LTLTSDESTLIEDGGAR |
| 27 | Q9NVJ2 | ARL8b | DLPNALDEK(141)QLIEK |
| 28 | Q9P0J6 | MRPL36 | DCYLVK(83)R |
| 29 | Q9Y3E0 | GOLT1B | SFVDK(130)VGESNNMV |
| 30 | Q9Y487 | ATP6V0A2 | LGAK(172)LGFVSGLINQGK |

**Downregulated**

| | Protein ID | Protein Name | Peptide sequence (GG-K) Huib32-4h and 72h |
|---|---|---|---|
| 1 | Q15043 | SLC39A14 | LQNGDLDHMIPQHCSSELDGK(297)APMVDEK |
| 2 | Q8NFA0 | USP32 | NFPQDNQK(983)VR |

**Supplementary Figure 5**. **Overview of ubiquitomics data. A.** Venn diagrams showing the overlap of ubiquitinated proteins in MelJuSo cells treated with 10 µM **Huib32** for 4 h and 72 h compared to DMSO-treated controls at the same time points. The left panel represents proteins with increased ubiquitination (upregulated), while the right panel represents proteins with decreased ubiquitination (downregulated). **B.** Lists of proteins with altered ubiquitination at the same lysine residue(s) following 4h or 72 h **Huib32** treatment. The changes include both increases and decreases in ubiquitination levels. Source data are provided as Supplementary Data 6 and 7. Related to Figures 4C and 4D.

Supplementary Figure 6

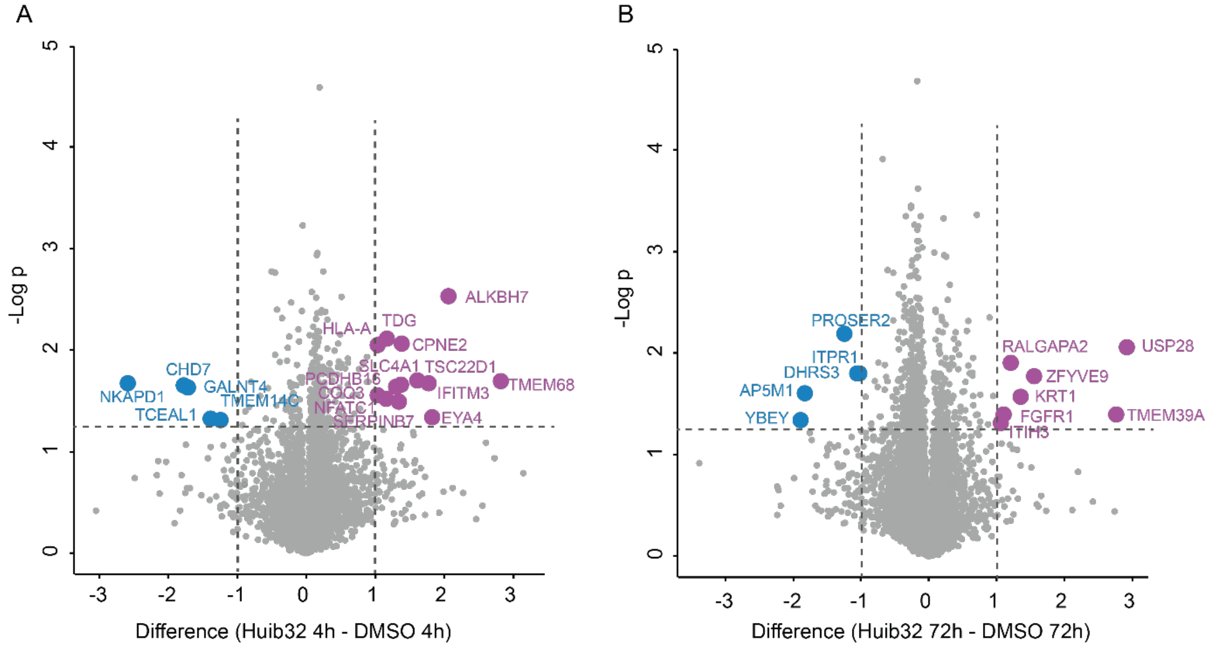

**Supplementary Figure 6**. **(A, B)** Volcano plots of the total proteome analysis of MelJuSo cells treated with DMSO or 10 µM final concentration of **Huib32**. Dashed lines indicate a significance cutoff at a p-value of 0.05 (-log10 = 1.3) in the y-axis and log2 ≥ 1 cutoff at fold change in the x-axis, n=3 independent experiments. All the proteins identified that are enriched with **Huib32** treatment are highlighted in purple, while the proteins downregulated by **Huib32** treatment are highlighted in blue. *Left volcano plot (A)*: MelJuSo cells incubated 4h with DMSO or **Huib32**. *Right volcano plot (B)*: MelJuSo cells incubated 72 h with DMSO or **Huib32**. Source data are provided as Supplementary Data S7. Related to Figure 4C.

Supplementary Figure 7

A

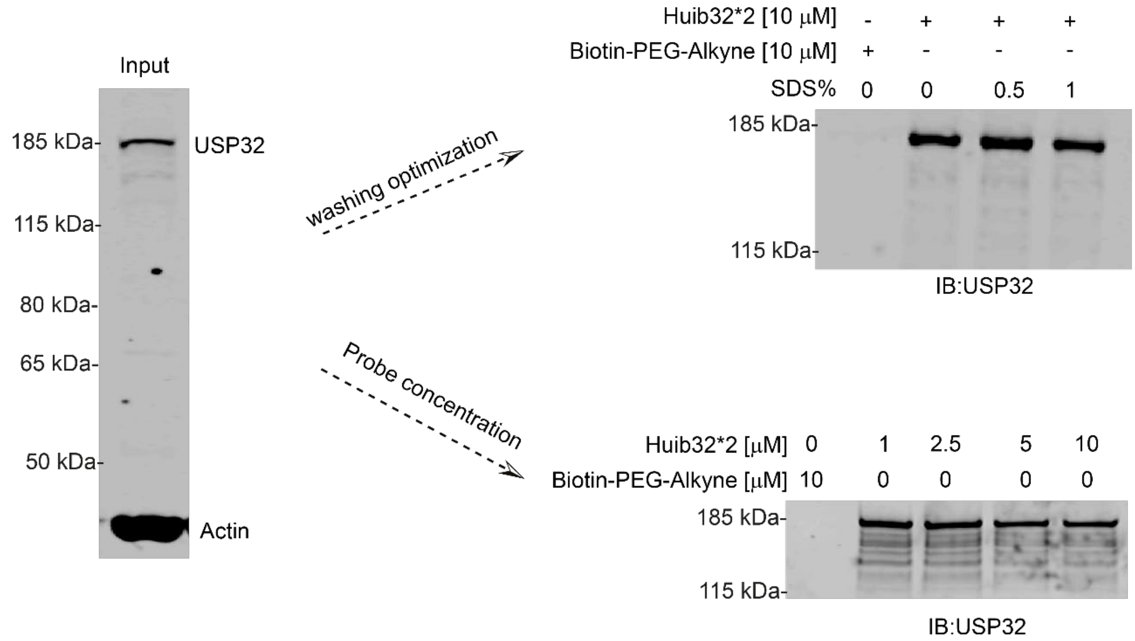

B

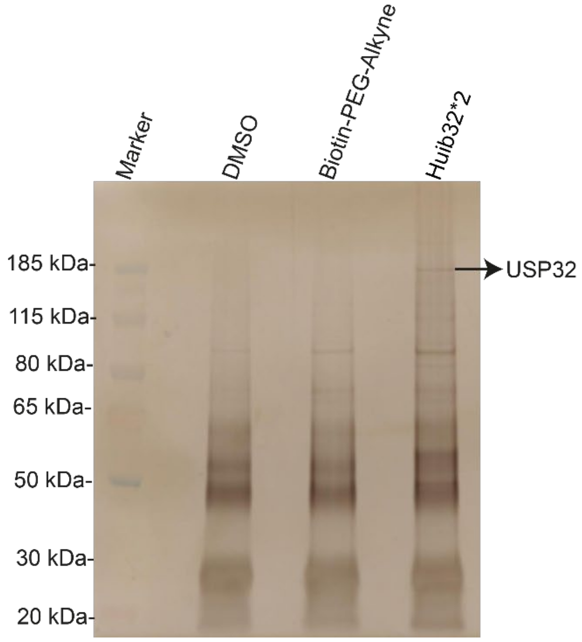

**Supplementary Figure 7**. **Optimization of washing conditions and Huib32*2 probe concentration. A.** USP32 and β-actin protein levels in MelJuSo cell lysates (input sample) was analyzed by immunoblotting against USP32 and β-actin (left panel). MelJuSo cell lysates were treated with 10 µM **Huib32*2** or Biotin-PEG-Alkyne and subjected to pulldown with NeutrAvidin beads. Beads were washed with lysis buffer containing varying SDS concentrations (0–1%). Boiled bead samples were analyzed for USP32 levels by immunoblotting. (right panel, top gel). MelJuSo cell lysates were treated with varying **Huib32*2** concentrations (0–10 μM), followed by pulldown and washing with lysis buffer containing 1% SDS. Pulldown samples were analyzed for USP32 by immunoblotting (right panel, bottom gel). **B.** Silver-stained SDS-PAGE gel showing enriched protein profiles from pulldowns performed with DMSO, Biotin-PEG4-Alkyne, and **Huib32*2** treatments. NeutrAvidin bead eluates were analyzed to compare protein enrichment across conditions. Related to Figure 6.

Supplementary Figure 8

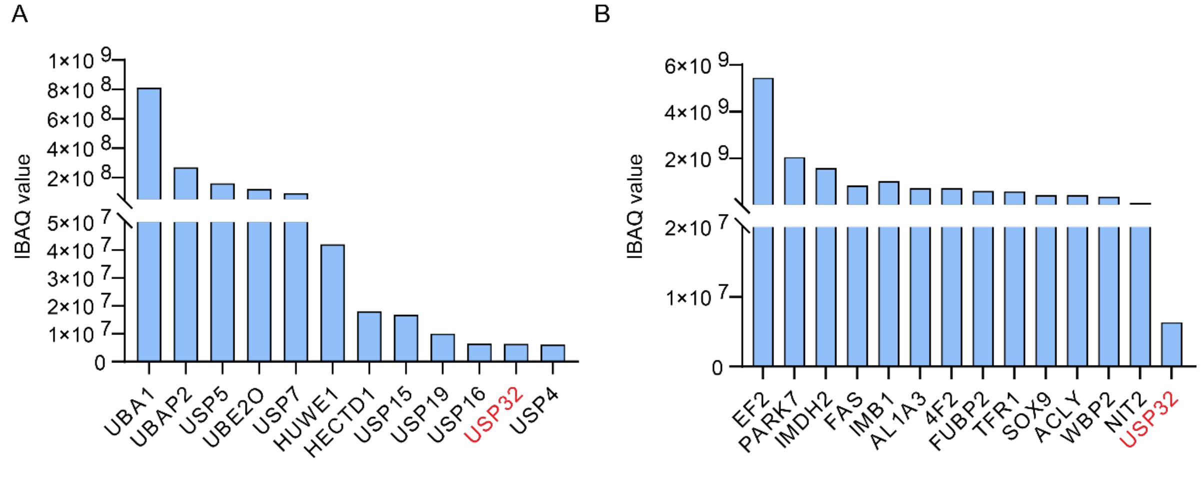

**Supplementary Figure 8. Total proteomics of MeljuSo cells.** Reference intensities (iBAQ values) of proteins shown in Figure 6 in MelJuSo cells. IBAQ values come from label-free total proteomics of MelJuSo cells used in this study. Source data are provided as Supplementary Data 9. **A.** IBAQ values of enzymes related to the Ub system identified in the pull-down LC-MSMS experiment. **B.** IBAQ values of top-15 highest-ranked proteins from the pull-down LC-MSMS experiment.

#### Synthesis

##### General materials and methods for synthesis

**Reagents, solvents and solutions.** General reagents were purchased from Sigma-Aldrich, Acros and Combi-Blocks, Inc. and were used as received. Solvents were purchased from VWR, Biosolve or Sigma-Aldrich. Thin Layer Chromatography (TLC) was performed on Merck aluminum sheets (pre-coated with silica gel 60 $F_{254}$). Compounds were visualized by UV adsorption (254 nm) and by using a solution of $KMnO_4$ (7.5 g $L^{-1}$) and $K_2CO_3$ (50 g $L^{-1}$) in $H_2O$ or a solution of ninhydrin (15 g $L^{-1}$) in 3% AcOH/EtOH v/v.

**Purification Techniques.** Compounds (unless stated otherwise) were purified on a Büchi Sepacore automatic flash chromatography system X10/X50. The Büchi Sepacore system was equipped with two Büchi pump modules C-605, a Büchi control unit C-620, Büchi fraction collector C-660 and a Büchi UV Photometer C-640. The silica columns were purchased at GraceResolv™ and were packed with a grade of Davisil® silica.

**Instrumentation for Compound Characterization**. NMR spectra ($^1H$, $^{13}C$) were recorded on a Bruker Ultrashield 300 MHz spectrometer at 298 K. Resonances are indicated with symbols 'd' (doublet), 's' (singlet), 't' (triplet) and 'm' (multiplet). Chemical shifts (δ) are given in ppm relative to $CDCl_3$, DMSO-$d_6$ or $CD_3OD$ as an internal standard and coupling constants (*J*) are quoted in hertz (Hz). LC-MS measurements were performed on an LC-MS system equipped with a Waters 2795 Separation Module (Alliance HT), a Waters 2996 Photodiode Array Detector (190–750 nm), an Xbridge C18 column (2.1 × 100 mm, 3.5 μm) and an LCT ESI-Orthogonal Acceleration Time of Flight Mass Spectrometer. Samples were run using 2 mobile phases: A = 1% $CH_3CN$ and 0.1% formic acid in $H_2O$ and B = 1% $H_2O$ and 0.1% formic acid in $CH_3CN$. Data processing was performed using Waters MassLynx Mass Spectrometry Software 4.1. LC-MS Program: Waters Xbridge C18 column (2.1 × 100 mm, 3.5 μm); flow rate = 0.4 mL $min^{-1}$, runtime = 13 min, column T = 40 °C, mass detection: 100–1500 Da. Gradient: 0–0.4 min: 5% B; 0.4–9.0 min: 5% → 95% B; 9.0–11.2 min: 95% B; 11.2–11.3 min: 95% → 5% B; 11.3–13.00 min: 5% B. Electrospray Ionization (ESI) highresolution mass spectrometry was carried on a Waters XEVO-G2 XS Q-TOF mass spectrometer equipped with an electrospray ion source in positive mode (capillary voltage: 3.0 kV, desolvation gas flow: 900 L h$^{-1}$, temperature: 60 °C) with a resolution R = 22,000 using 200 pg µL$^{-1}$ Leu-Enk (*m/z* = 556.2771) as a "lock mass". Samples were run using 2 mobile phases: A = 0.1% formic acid in $H_2O$ and B = 0.1% formic acid in $CH_3CN$ on a Waters Acquity UPLC BEH C18 column (2.1 × 50 mm, 1.7 µm); flow rate = 0.6 mL min$^{-1}$, runtime = 3.00 min, column T = 60 °C, mass detection: 50–1500 Da. Gradient: 0–0.15 min: 2% B; 0.15–1.85 min: 2% → 100% B; 1.85–2.05: 100% B; 2.05–2.10 min: 100% → 2% B; 2.10–3.00 min: 100% B. Data processing was performed using Waters MassLynx Mass Spectrometry Software 4.1.

**General procedure for amide synthesis.**

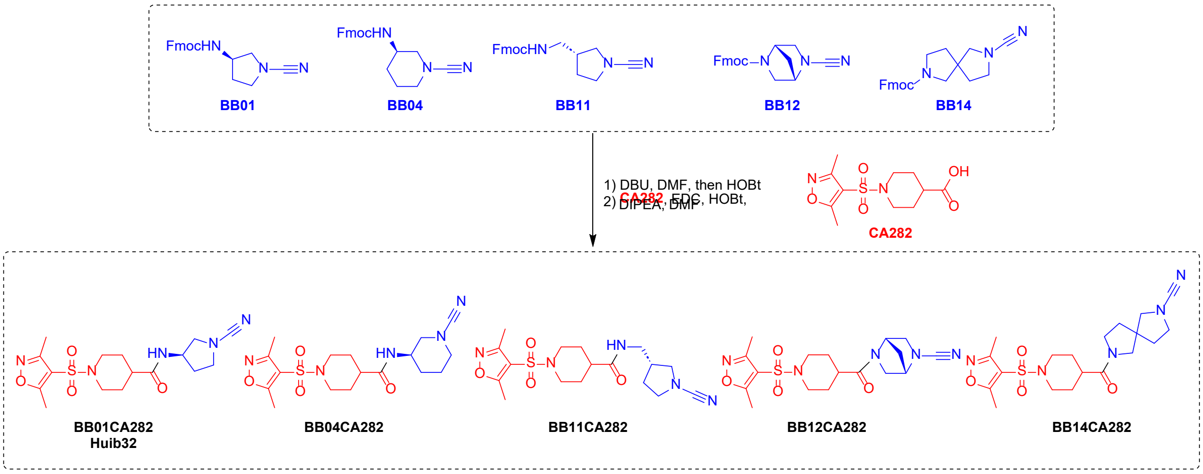

The Fmoc-protected amine was *in situ* deprotected by the addition of 1,8-Diazabicyclo[5.4.0]undec-7-ene (DBU) (3.4 µL, 22.5 µmol, 0.5 eq.) to a solution of the Fmoc-protected amine (BBx) (45 µmol, 1.0 eq.) in DMF (0.5 mL). After 30 min. at room temperature, TLC analysis revealed complete deprotection and the reaction was quenched by addition of HOBt (6 mg, 45 µmol, 1.0 eq.) A solution containing the pre-activated carboxylic acid, containing 1-((3,5-dimethylisoxazol-4-yl)sulfonyl)piperidine-4-carboxylic acid (**CA282**) (14.4 mg, 49.5 µmol, 1.1 eq.), HOBt (15.2 mg, 112.5 µmol, 2.5 eq.), EDC.HCl (13.0 mg, 67.5 µmol, 1.5 eq.) and DIPEA (23.5 µL, 49.5 µmol, 1.1 eq.) in DMF (0.5 mL), was added. The mixture was stirred for 16h at room temperature, until TLC analysis indicated complete consumption of the starting material. The mixture was concentrated *in vacuo* and the residue was dissolved in EtOAc (5 mL).

The organic layer was washed with 1M HCl (1 x 2 mL), sat. $NaHCO_3$ (1 x 2 mL) and sat. NaCl (1 x 2 mL). The organic layer was dried over $Na_2SO_4$, filtered and concentrated *in vacuo* to obtain the crude product. The compound was isolated by Büchi flash chromatography (DCM → 5% MeOH/DCM).

**(*R*)-*N*-(1-cyanopyrrolidin-3-yl)-1-((3,5-dimethylisoxazol-4-yl)sulfonyl)piperidine-4-carboxamide**(**BB01CA282, Huib32**) was synthesized according to the general procedure for amide synthesis starting from (9*H*-fluoren-9-yl)methyl (*R*)-(1-cyanopyrrolidin-3-yl)carbamate (**BB01**) (15 mg, 45 μmol, 1.0 eq.) and 1-((3,5-dimethylisoxazol-4-yl)sulfonyl)piperidine-4-carboxylic acid (**CA282**) (14.3 mg, 49.5 μmol, 1.1 eq.). The title compound was obtained as a white solid (11.7 mg, 30.6 μmol, 68%). $^1$H NMR (300 MHz, $CDCl_3$) δ 6.21 (d, J = 7.1 Hz, 1H), 4.56 – 4.42 (m, 1H), 3.74 (d, J = 11.8 Hz, 2H), 3.63 (dd, J = 10.3, 5.7 Hz, 1H), 3.52 (dd, J = 8.5, 5.8 Hz, 2H), 3.28 (dd, J = 10.3, 2.0 Hz, 1H), 2.71 – 2.56 (m, 5H), 2.40 (s, 3H), 2.25 – 2.07 (m, 2H), 1.97 – 1.77 (m, 5H). $^{13}$C NMR (75 MHz, $CDCl_3$) δ 173.85, 173.70, 158.10, 117.09, 113.84, 55.85, 49.56, 48.72, 44.88, 44.86, 41.77, 31.98, 28.19, 28.13, 13.11, 11.46. HR-MS calculated for $C_{16}H_{23}N_5O_4S$ $[M+H]^+$ 382.1549, found 382.1555.

**(*R*)-*N*-(1-cyanopiperidin-3-yl)-1-((3,5-dimethylisoxazol-4-yl)sulfonyl)piperidine-4-carboxamide** (**BB04CA282**) was synthesized according to the general procedure for amide synthesis starting from (9*H*-fluoren-9-yl)methyl (*R*)-(1-cyanopyrrolidin-3-yl)carbamate (**BB04**) (15.6 mg, 45 μmol, 1.0 eq.) and 1-((3,5-dimethylisoxazol-4-yl)sulfonyl)piperidine-4-carboxylic acid (**CA282**) (14.3 mg, 49.5 μmol, 1.1 eq.). The title compound was obtained as a white solid (13.2 mg, 33.3 μmol, 74.0%). $^1$H NMR (300 MHz, $CDCl_3$) δ 5.69 (d, J = 7.6 Hz, 1H), 4.12 – 4.02 (m, 1H), 3.80 – 3.68 (m, 2H), 3.36 (dd, J = 12.7, 3.3 Hz, 1H), 3.29 – 3.12 (m, 2H), 3.08 (dd, J = 12.7, 5.6 Hz, 1H), 2.64 (s, 5H), 2.40 (s, 3H), 2.26 – 2.10 (m, 1H), 2.04 – 1.55 (m, 8H). $^{13}$C NMR (75 MHz, $CDCl_3$) δ 173.85, 173.13, 158.10, 117.93, 113.85, 53.37, 49.81, 44.89, 44.82, 44.00, 42.03, 28.23, 28.20, 28.06, 21.30, 13.11, 11.45. HR-MS calculated $C_{17}H_{25}N_5O_4S[M+H]^+$ 396.1706, found 396.1722.

**(*R*)-*N*-((1-cyanopyrrolidin-3-yl)methyl)-1-((3,5-dimethylisoxazol-4-yl)sulfonyl)piperidine-4-carboxamide** (**BB11CA282**) was synthesized according to the general procedure for amide synthesis starting from (9*H*-fluoren-9-yl)methyl (*R*)-((1-cyanopyrrolidin-3-yl)methyl)carbamate (**BB11**) (15.6 mg, 45 µmol, 1.0 eq.) and 1-((3,5-dimethylisoxazol-4-yl)sulfonyl)piperidine-4-carboxylic acid (**CA282**) (14.3 mg, 49.5 µmol, 1.1 eq.). The title compound was obtained as a white solid (13.6 mg, 34.3 µmol, 76%). $^1$H NMR (300 MHz, $CDCl_3$) δ 5.90 (t, J = 6.1 Hz, 1H), 3.81 – 3.68 (m, 2H), 3.56 – 3.27 (m, 4H), 3.25 – 3.08 (m, 2H), 2.64 (s, 5H), 2.48 (p, J = 7.1 Hz, 1H), 2.39 (s, 3H), 2.24 – 2.09 (m, 1H), 2.09 – 1.54 (m, 6H).$^{13}$C NMR (75 MHz, $CDCl_3$) δ 174.01, 173.85, 158.07, 117.48, 113.81, 53.78, 49.95, 44.90, 42.03, 41.33, 39.41, 29.31, 28.30, 13.11, 11.45. HR-MS calculated for $C_{17}H_{25}N_5O_4S$ [M+H]$^+$ 396.1706, found 396.1697.

**(1*R*,4*R*)-5-(1-((3,5-dimethylisoxazol-4-yl)sulfonyl)piperidine-4-carbonyl)-2,5-diazabicyclo[2.2.1]heptane-2-carbonitrile** (**BB12CA282**) was synthesized according to the general procedure for amide synthesis starting from (9*H*-fluoren-9-yl)methyl (1*R*,4*R*)-5-cyano-2,5-diazabicyclo[2.2.1]heptane-2-carboxylate (**BB12**) (15.6 mg, 45 µmol, 1.0 eq.) and 1-((3,5-dimethylisoxazol-4-yl)sulfonyl)piperidine-4-carboxylic acid (**CA282**) (14.3 mg, 49.5 µmol, 1.1 eq.). The title compound was obtained as a white solid (9.3 mg, 23.6 µmol, 52%). $^1$H NMR (300 MHz, $CDCl_3$) δ 4.93 (s, 1H), 4.32 (s, 1H), 3.84 – 3.71 (m, 3H), 3.59 – 3.49 (m, 2H), 3.45 – 3.33 (m, 1H), 2.74 – 2.61 (m, 5H), 2.43 (s, 3H), 2.38 – 2.11 (m, 1H), 2.08 – 1.75 (m, 6H). $^{13}$C NMR (75 MHz, $CDCl_3$) δ 173.74, 172.15, 158.00, 141.97, 116.36, 61.99, 59.03, 54.96, 52.84, 44.65, 44.53, 38.79, 35.75, 27.36, 27.27, 12.99, 11.34. HR-MS calculated for $C_{17}H_{23}N_5O_4S$ [M+H]$^+$ 394.1549, found 394.1547.

**(*S*)-7-(1-((3,5-dimethylisoxazol-4-yl)sulfonyl)piperidine-4-carbonyl)-2,7-diazaspiro[4.4]nonane-2-carbonitrile** (**BB14CA282**) was synthesized according to the general procedure for amide synthesis starting from (9*H*-fluoren-9-yl)methyl (*S*)-7-cyano-2,7-diazaspiro[4.4]nonane-2-carboxylate (**BB14**) (15.6 mg, 45 µmol, 1.0 eq.) and 1-((3,5-dimethylisoxazol-4-yl)sulfonyl)piperidine-4-carboxylic acid (**CA282**) (14.3 mg, 49.5 µmol, 1.1 eq.). The title compound was obtained as a white solid (9.8 mg, 23.2 µmol, 52%). $^1$H NMR (300 MHz, $CDCl_3$) δ 3.67 (d, J = 11.8 Hz, 2H), 3.24 (d, J =

1.7 Hz, 8H), 2.68 – 2.54 (m, 5H), 2.33 (d, J = 1.4 Hz, 4H), 2.05 – 1.69 (m, 8H).$^{13}$C NMR (75 MHz, $CDCl_3$) δ 173.84, 172.64, 172.50, 158.14, 58.86, 58.78, 55.26, 54.15, 50.08, 49.69, 47.77, 45.48, 45.17, 44.85, 44.79, 39.37, 39.03, 35.20, 35.00, 34.82, 33.04, 27.60, 13.13, 11.48. HR-MS calculated for $C_{17}H_{25}N_5O_4S$ $[M+H]^+$ 422.1862, found 422.1862.

**Synthesis of Huib32* and probes.**

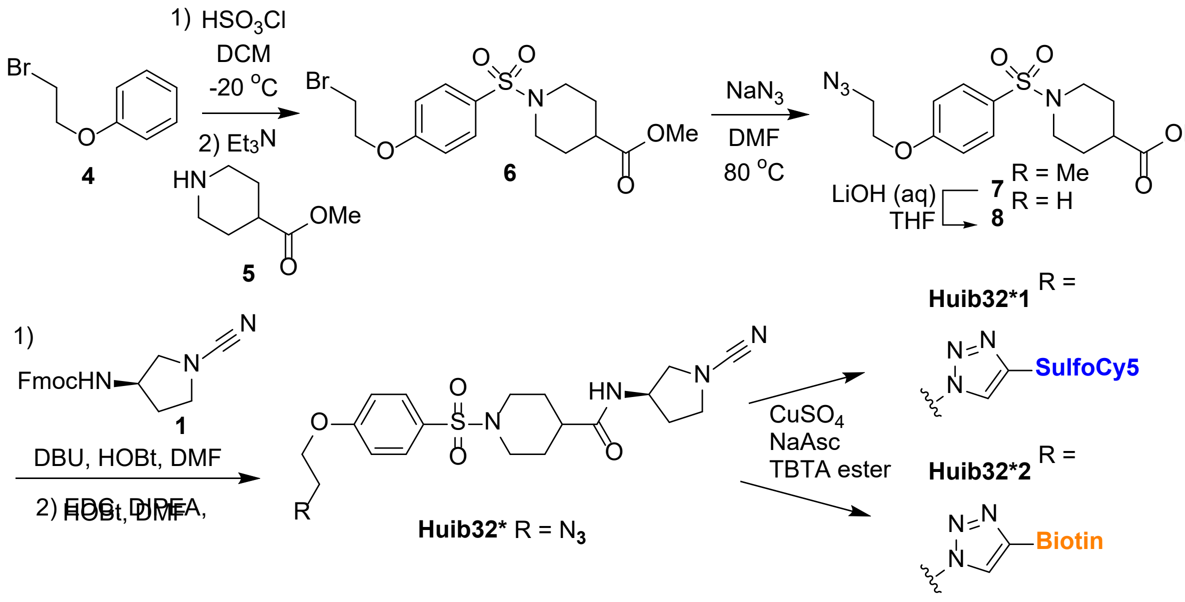

**Methyl 1-((4-(2-bromoethoxy)phenyl)sulfonyl)piperidine-4-carboxylate** (**6**). A solution of (2-bromoethoxy)benzene (**4**) (1.00 g, 4.97 mmol, 1.0 eq.) in DCM (100 mL) was cooled to -20 °C using a salted ice bath and chlorosulfuric acid (993 µL, 14.9 mmol, 3.0 eq.) was added. After stirring for 90 min. at -20 °C TLC analysis indicated complete consumption of the starting material (**4**). The reaction mixture was warmed to room temperature and quenched by pouring the mixture onto ice. The aqueous layer was quickly extracted with EtOAc (100 mL). The organic layer was dried over $Na_2SO_4$, filtered and concentrated *in vacuo* to obtain the crude intermediate as a white solid, which was used in the next reaction without further purification. The crude intermediate was dissolved in DCM (50 mL) and cooled to 0 °C. To this solution was added $Et_3N$ (2.08 mL, 14.9 mmol, 3.0 eq.) and methyl piperidine-4-carboxylate (**5**) (739 µL, 5.47 mmol, 1.1 eq.). The mixture was allowed to warm to room temperature and stirred for 5 h, until TLC analysis indicated complete consumption of the sulfonyl chloride intermediate. EtOAc (200 mL) was added to the reaction and the organic layer was washed with 1M HCl (1 x 200 mL), sat. $NaHCO_3$ (1 x 200 mL) and sat. NaCl (1 x 200 mL). The organic layer was dried over $Na_2SO_4$, filtered and concentrated *in vacuo* to obtain the crude product as an off white solid. The product was isolated by Büchi flash chromatography (Heptane → EtOAc) to obtain the title compound (**6**) as a white solid (1.18 g, 2.90 mmol, 58%). $^1$H NMR (300 MHz, $CDCl_3$) δ 7.69 (d, $J$ = 8.8 Hz, 2H), 7.00 (d, $J$ = 8.9 Hz, 2H), 4.35 (t, $J$ = 6.1 Hz, 2H), 3.71 – 3.55 (m, 4H), 3.65 (s, 3H), 2.45 (td, $J$ = 11.3, 3.0 Hz, 2H), 2.32 – 2.20 (m, 1H), 2.03 – 1.91 (m, 2H), 1.81 (qd, $J$ = 10.5, 5.3 Hz, 2H). $^{13}$C NMR (75 MHz, $CDCl_3$) δ 174.39, 161.58, 129.96, 128.65, 114.93, 68.16, 52.02, 45.52, 40.02, 27.54, 14.25.

**Methyl 1-((4-(2-azidoethoxy)phenyl)sulfonyl)piperidine-4-carboxylate** (**7**). A solution of compound (**6**) (1.18 g, 2.90 mmol, 1.0 eq.) in DMF (50 mL) was added to a flask containing $NaN_3$ (755 mg, 11.62 mmol, 4.0 eq.). The mixture was heated to 80 °C and stirred for 16 h, after which TLC analysis indicated complete consumption of the starting material (**6**). The reaction was cooled to room temperature. EtOAc (200 mL) was added to the mixture and the organic layer was washed with water (200 mL), sat. $NaHCO_3$ (200 mL) and 1M LiCl (100 mL). The organic layer was dried over $Na_2SO_4$, filtered and concentrated *in vacuo* to obtain the crude product as a yellow oil. The title compound was isolated by Büchi flash chromatography (Heptane → EtOAc) to obtain the title compound (**7**) as a white solid (724 mg, 1.96 mmol, 68%). $^1$H NMR (300 MHz, $CDCl_3$) δ 7.67 (dd, J = 8.9, 2.0 Hz, 2H), 7.00 (d, J = 8.9 Hz, 1H), 4.19 (t, J = 4.9 Hz, 2H), 3.68 – 3.51 (m, 8H), 2.43 (t, J = 11.4 Hz, 1H), 2.30 – 2.18 (m, 1H), 2.01 – 1.90 (m, 2H), 1.86 – 1.71 (m, 3H). $^{13}$C NMR (75 MHz, $CDCl_3$) δ 174.35, 161.66, 129.88, 128.47, 114.83, 67.39, 51.96, 50.03, 45.49, 39.95, 27.48.

**1-((4-(2-azidoethoxy)phenyl)sulfonyl)piperidine-4-carboxylic acid** (**8**). To a solution of compound (**7**) (724 mg, 1.96 mmol, 1.0 eq.) in THF (5 mL) was added 1.5 M $LiOH_{(aq)}$ (5 mL) and the reaction was stirred for 3 h, after which TLC analysis indicated complete consumption of the starting material. The reaction mixture was washed with DCM (50 mL) . The aqueous layer was acidified with 2M HCl until pH 2 and extracted with DCM (3 x 25 mL). The latter organic layers were combined and dried over $Na_2SO_4$, filtered and concentrated *in vacuo* to obtain the title compound as a white solid (700 mg, 1.97 mmol, quant.). $^1$H NMR (300 MHz, $CDCl_3$) δ 7.76 – 7.64

(m, 2H), 7.07 – 6.96 (m, 2H), 4.21 (t, J = 4.5 Hz, 2H), 3.69 – 3.56 (m, 4H), 2.54 – 2.39 (m, 2H), 2.30 (m, 1H), 2.07 – 1.92 (m, 1H), 1.91 – 1.72 (m, 2H). $^{13}C$ NMR (75 MHz, $CDCl_3$) δ 179.48, 161.75, 129.97, 128.59, 114.91, 67.45, 50.10, 45.46, 39.82, 27.30.

**(*R*)-1-((4-(2-azidoethoxy)phenyl)sulfonyl)-N-(1-cyanopyrrolidin-3-yl)piperidine-4-carboxamide** (**Huib32***). This compound was synthesized according to the general procedure for amide synthesis starting from (9*H*-fluoren-9-yl)methyl (*R*)-(1-cyanopyrrolidin-3-yl)carbamate (**BB01**) (15 mg, 45 µmol, 1.0 eq.) and 1-((4-(2-azidoethoxy)phenyl)sulfonyl)piperidine-4-carboxylic acid (**8**) (15.9 mg, 49.5 µmol, 1.1 eq.). The title compound was obtained as a white solid (15.1 mg, 22.4 µmol, 68%). $^{1}H$ NMR (300 MHz, $CDCl_3$) δ 7.69 (d, 2H), 7.02 (d, 2H), 6.43 (d, J = 7.6 Hz, 1H), 4.51 – 4.37 (m, 1H), 4.21 (t, J = 4.4 Hz, 2H), 3.72 (d, J = 11.7 Hz, 2H), 3.65 (t, J = 4.4 Hz, 2H), 3.59 (dd, J = 10.2, 5.8 Hz, 1H), 3.53 – 3.43 (m, 2H), 3.31 – 3.20 (m, 1H), 2.34 (dt, J = 11.3, 3.6 Hz, 2H), 2.20 – 1.97 (m, 2H), 1.96 – 1.69 (m, 5H). $^{13}C$ NMR (75 MHz, $CDCl_3$) δ 174.10, 161.79, 129.91, 128.51, 117.21, 114.95, 87.83, 67.46, 55.65, 50.08, 49.48, 48.76, 45.59, 41.97, 31.78, 28.08. HR-MS calculated for $C_{19}H_{25}N_7O_4S$ $[M+H]^+$ 448.1767, found 448.1823.

**General procedure for click chemistry.**

The azide (10 mg, 1.0 eq.) and alkyne (1.1 eq.) were dissolved DMSO (2 mL) and the mixture was purged with argon. To this solution was added 60 µL of a premade mixture (“click mix”) of 1:1:1 v/v/v $CuSO_4 \cdot 5H_2O$ 100 mM in $H_2O$, sodium ascorbate 600 mM in $H_2O$ and TBTA ester 100 mM in ACN. The reaction mixture was stirred and monitored by LC-MS every 30 min. In case the starting compound (azide) was not fully consumed, another 60 µL “click mix” was added until full conversion was achieved. Sat. NaCl (10 mL) was added and the mixture was extracted with DCM (3 x 20 mL). The combined organic layers were concentrated and the product was purified by Büchi flash chromatography (DCM → 5% MeOH/DCM).

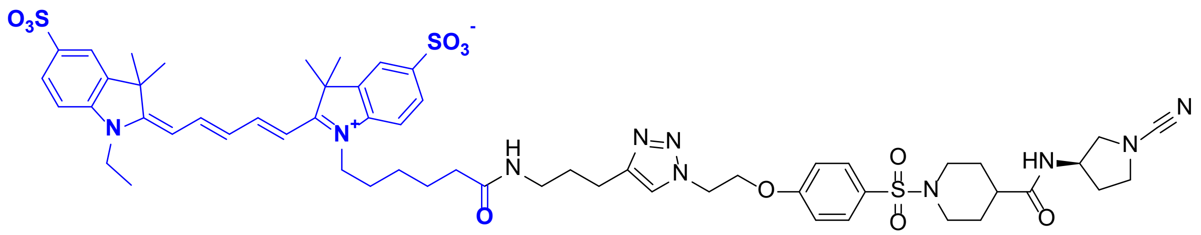

Huib32*1

**SulfoCy5 USP32 probe (Huib32*1).** This compound was synthesized according to the general procedure for click chemistry starting from compound (**Huib32***) (10 mg, 22.4 µmol, 1.0 eq.) and 3,3-dimethyl-1-(6-oxo-6-(pent-4-yn-1-ylamino)hexyl)-2-((1*E*,3*E*)-5-((*E*)-1,3,3-trimethyl-5-sulfoindolin-2-ylidene)penta-1,3-dien-1-yl)-3*H*-indol-1-ium-5-sulfonate (SulfoCy5-alkyne) (17.4 mg, 24.6 µmol, 1.1 eq.). The title compound was obtained as a blue solid (14.4 mg, 12.3 µmol, 55%). $^{1}$H NMR (300 MHz, DMSO) δ 8.36 (t, J = 13.1 Hz, 2H), 8.05 (d, J = 6.6 Hz, 1H), 7.97 (s, 1H), 7.87 – 7.77 (m, 3H), 7.69 – 7.58 (m, 4H), 7.37 – 7.25 (m, 2H), 7.14 (t, 2H), 6.57 (t, J = 12.3 Hz, 1H), 6.30 (d, J = 2.2 Hz, 2H), 4.75 (t, J = 4.9 Hz, 2H), 4.49 (t, J = 5.1 Hz, 2H), 4.22 – 4.06 (m, 5H), 3.60 – 3.41 (m, 4H), 3.12 – 3.00 (m, 3H), 2.57 (t, 2H), 2.31 – 2.13 (m, 3H), 2.12 – 1.87 (m, 5H), 1.68 (s, 14H), 1.58 – 1.45 (m, 5H), 1.43 – 1.17 (m, 10H). $^{13}$C NMR (75 MHz, DMSO) δ 179.80, 178.44, 177.77, 177.65, 165.74, 158.80, 151.01, 147.55, 146.98, 145.91, 145.79, 145.33, 145.12, 133.57, 132.05, 130.65, 130.38, 126.91, 123.94, 118.65, 114.24, 114.03, 107.78, 107.59, 70.59, 58.91, 53.36, 53.28, 53.19, 53.08, 49.37, 47.54, 45.19, 42.85, 42.20, 39.22, 34.45, 32.70, 31.76, 30.72, 30.43, 30.29, 29.82, 29.04, 26.16, 15.11. HR-MS calculated for $C_{57}H_{73}N_{10}O_{11}S_3$ [M+H]$^+$ 1170.4701, found 1170.4663.

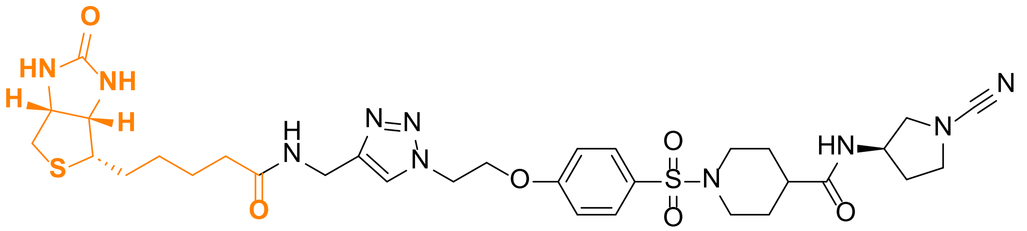

Huib32*2

**Biotin USP32 probe (Huib32*2).** This compound was synthesized according to the general procedure for click chemistry starting from compound (**Huib32***) (9 mg, 20 µmol, 1.0 eq.) and 5-((3a*S*,4*S*,6a*R*)-2-oxohexahydro-1*H*-thieno[3,4-d]imidazol-4-yl)-

*N*-(prop-2-yn-1-yl)pentanamide (Biotin-alkyne) (20 mg, 70 µmol, 3.5 eq.). The title compound was obtained as a white solid (0.46 mg, 0.6 µmol, 3%).

$^1$H NMR (300 MHz, DMSO) δ 8.36 (t, J = 13.1 Hz, 2H), 8.05 (d, J = 6.6 Hz, 1H), 7.97 (s, 1H), 7.87 – 7.77 (m, 3H), 7.69 – 7.58 (m, 4H), 7.37 – 7.25 (m, 2H), 7.14 (t, 2H), 6.57 (t, J = 12.3 Hz, 1H), 6.30 (d, J = 2.2 Hz, 2H), 4.75 (t, J = 4.9 Hz, 2H), 4.49 (t, J = 5.1 Hz, 2H), 4.22 – 4.06 (m, 5H), 3.60 – 3.41 (m, 4H), 3.12 – 3.00 (m, 3H), 2.57 (t, 2H), 2.31 – 2.13 (m, 3H), 2.12 – 1.87 (m, 5H), 1.68 (s, 14H), 1.58 – 1.45 (m, 5H), 1.43 – 1.17 (m, 10H). $^{13}$C NMR (75 MHz, DMSO) δ 173.61, 172.01, 162.72, 161.22, 145.19, 129.66, 127.66, 125.54, 117.28, 115.06, 107.53, 105.49, 66.64, 61.02, 59.19, 55.41, 54.91, 52.17, 48.72, 48.46, 45.27, 34.98, 34.09, 30.80, 28.20, 28.01, 27.63, 25.19. HR-MS calculated for $C_{32}H_{44}N_{10}O_6S_2$ $[M+H]^+$ 729.2965, found 729.2957.

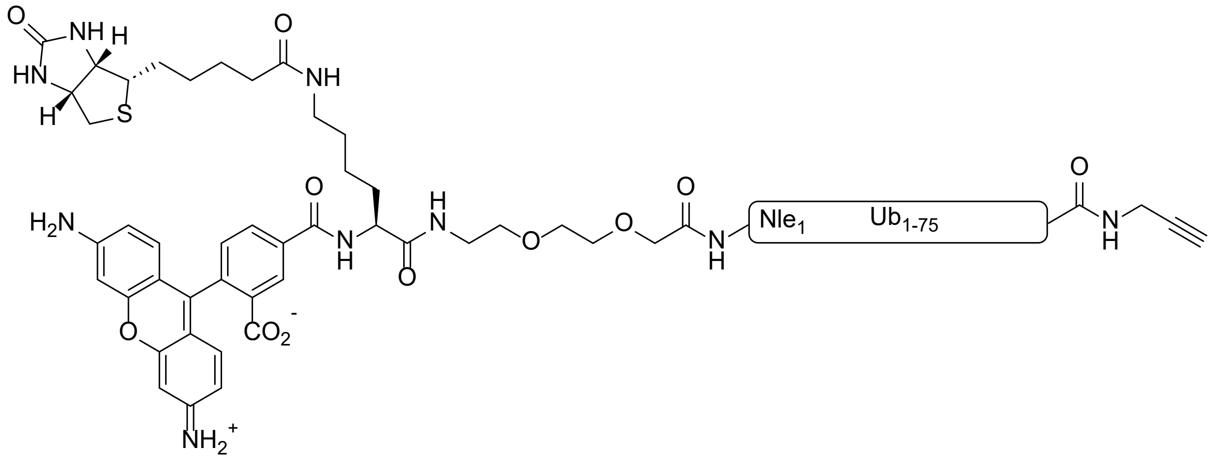

**RhoK(Biotin)UbPA probe.** The probe was synthesized according to the reported procedure for the SPPS of Ub.[1] In brief, SPPS was performed on a Syro II MultiSyntech Automated Peptide synthesizer using standard 9-fluorenylmethoxycarbonyl (Fmoc) based solid phase peptide chemistry. All amino acids were used in excess and dipeptides were used in positions determined previously. The K(Biotin)-PEG-Ub(1-75) was synthesized by SPPS on a TentaGel Trt R resin (Sigma-Aldrich) preloaded with Fmoc-Glycine on a 20 µmol scale. The Fmoc-L-Lys(biotin)-OH used in SPPS was obtained from Combi Blocks (Cat.ID SS-0990, CAS 146987-10-2). The N-terminal Met in the Ub sequence was changed to Norleucine (Nle) to avoid oxidation adducts. DiBoc-Rhodamine was coupled on-resin to the N-terminus of the K(Biotin)-PEG-Ub(1-75) using standard coupling conditions.[2]

After mild-acidic removal of the peptide from the resin using 20 vol% HFIP/DCE, the C-terminal propargyl group was coupled, followed by a global deprotection.[3]

The crude material was first dissolved in DMSO. This solution was slowly added to MQ water containing 0.05% TFA and filtered through a GfxO/0.45μm GHP membrane Acrodisc® Premium 25mm syringe filter. The sample was then purified by HPLC on a Waters XBridge™ Prep C18 Column (30 x 150 mm, 5μm OBD™) at a flow rate of 37.5 mL/min using the gradient outlined below with water containing 0.05% TFA (Solvent A) and acetonitrile containing 0.05% TFA (Solvent B) as eluents.

| Time (min.) | Eluent B (%) |
|---|---|
| 0 → 5 | 5 |
| 5 → 7 | 5 → 25 |
| 7 → 22 | 25 → 55 |
| 22 → 24 | 55 → 95 |
| 24 → 27 | 95 |
| 27 → 27.5 | 95 → 5 |
| 27.5 → 30 | 5 |

All fractions were analyzed using LC/MS and pure fractions were pooled and lyophilized. The purity of the protein was confirmed by LC-MS analysis. Data was processed using Waters MassLynx Mass Spectrometry Software v4.1. Mass was inferred by charge deconvolution (MaxEnt1 function) from mass over charge (m/z) measurements. Calculated MW 9400.3 Da.

#### $^{1}H$ NMR: BB01CA282, Huib32

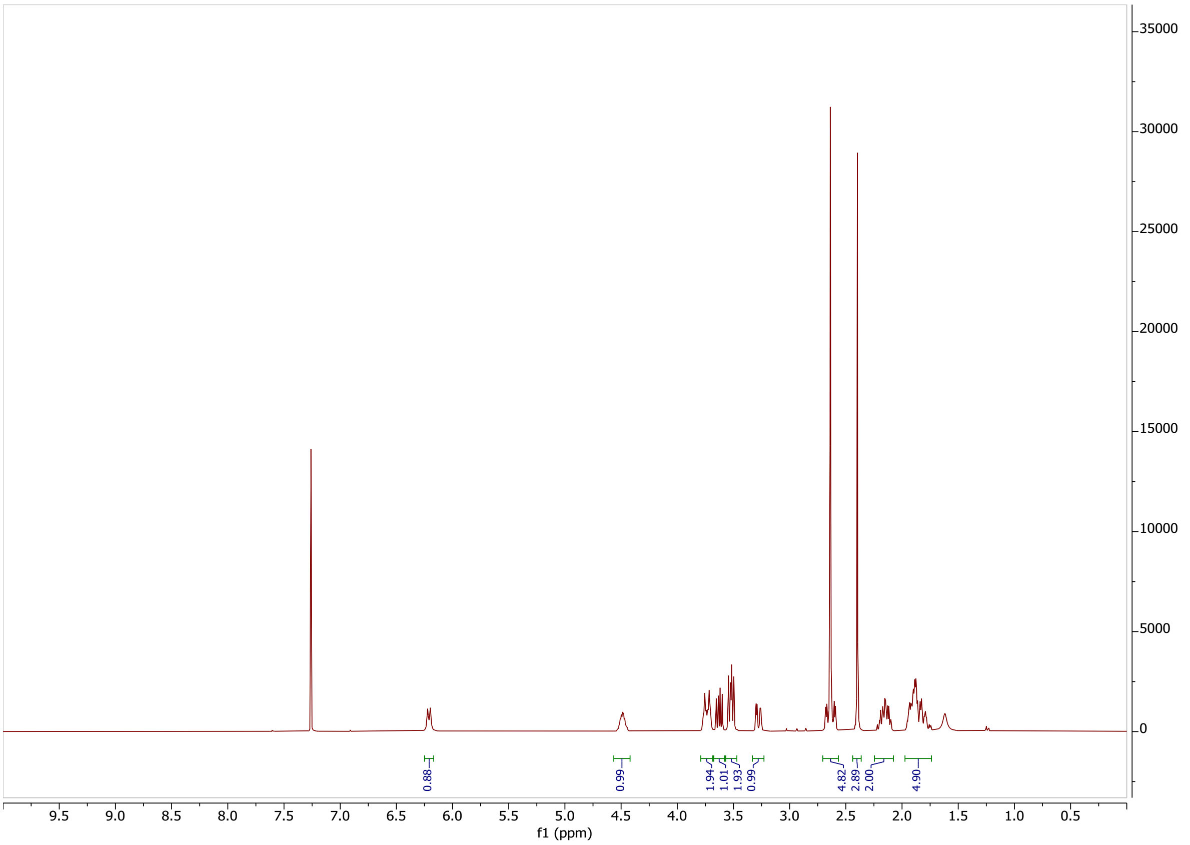

#### $^{13}C$ NMR: BB01CA282, Huib32

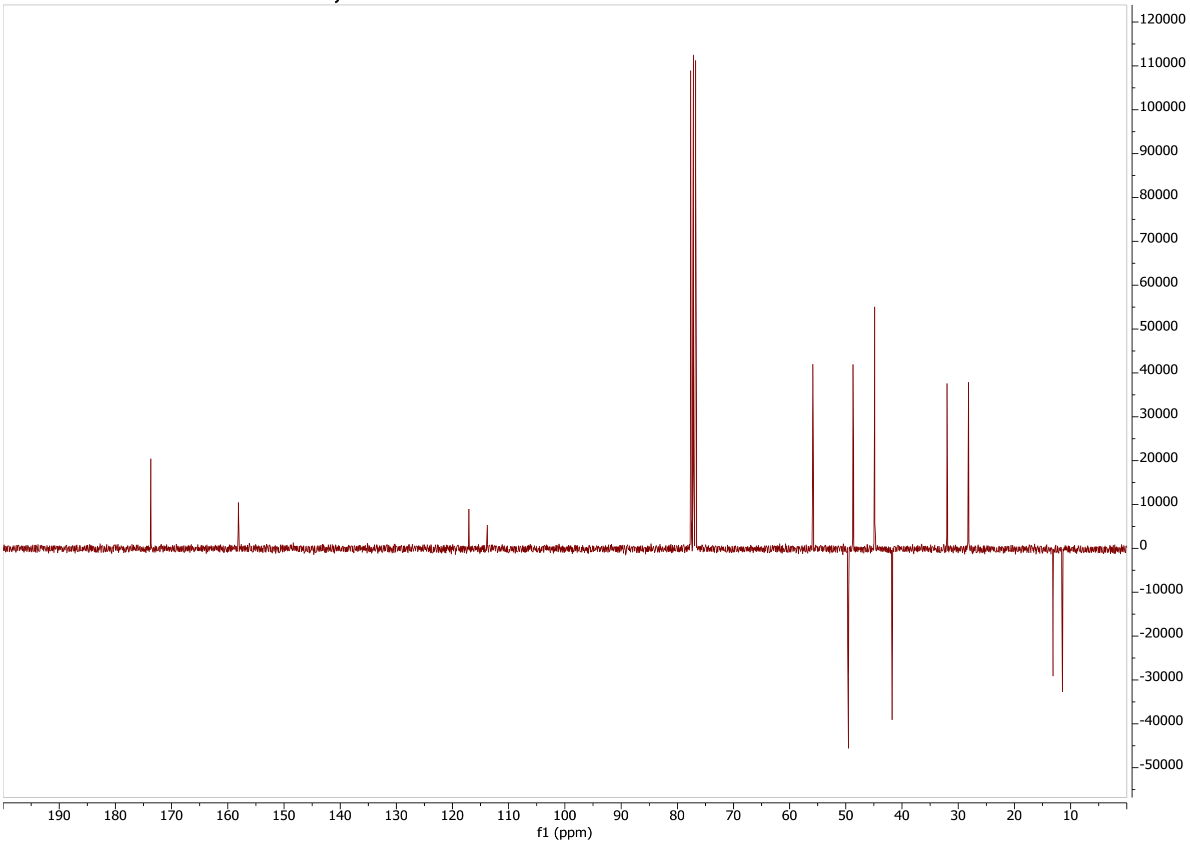

#### LC-MS: BB01CA282, Huib32

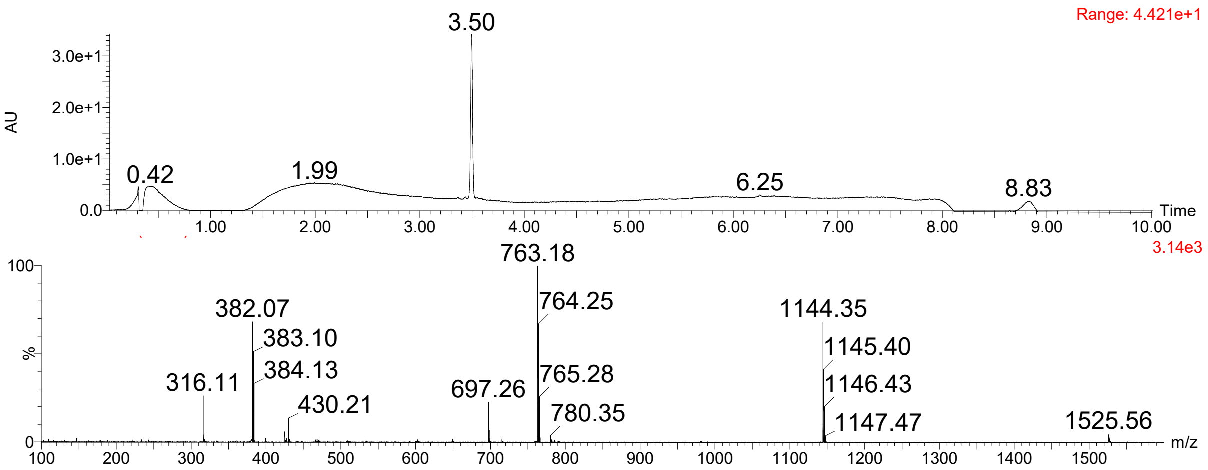

#### $^{1}$H-NMR: BB04CA282

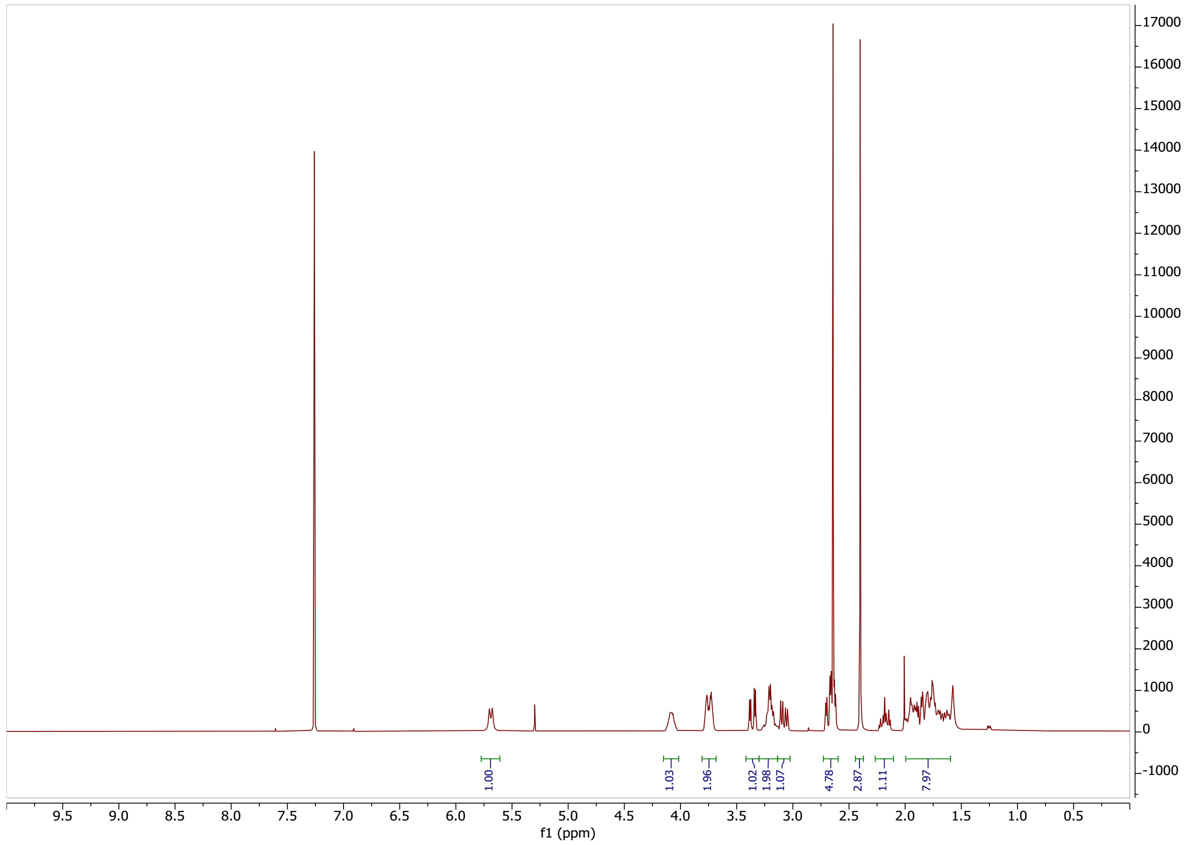

#### $^{13}$C-NMR: BB04CA282

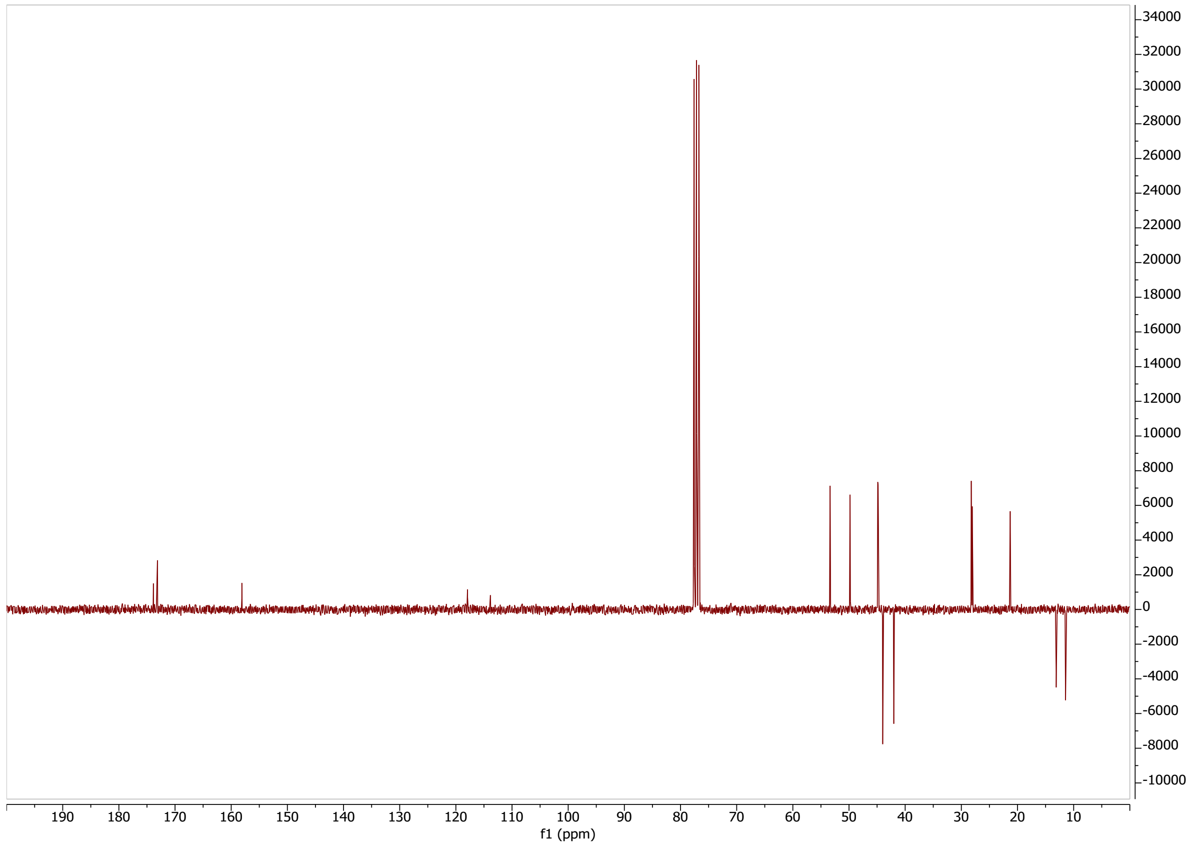

## LC-MS: BB04CA282

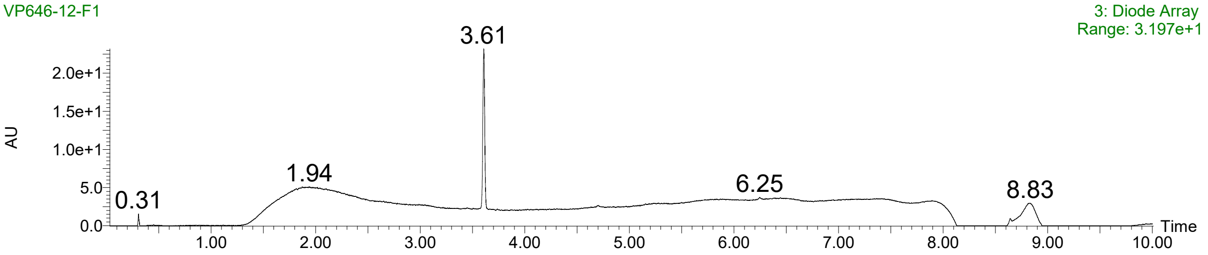

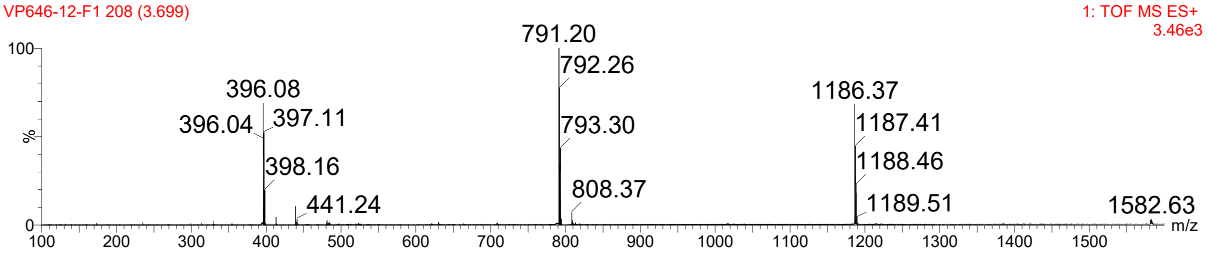

#### $^{1}$H-NMR: BB11CA282

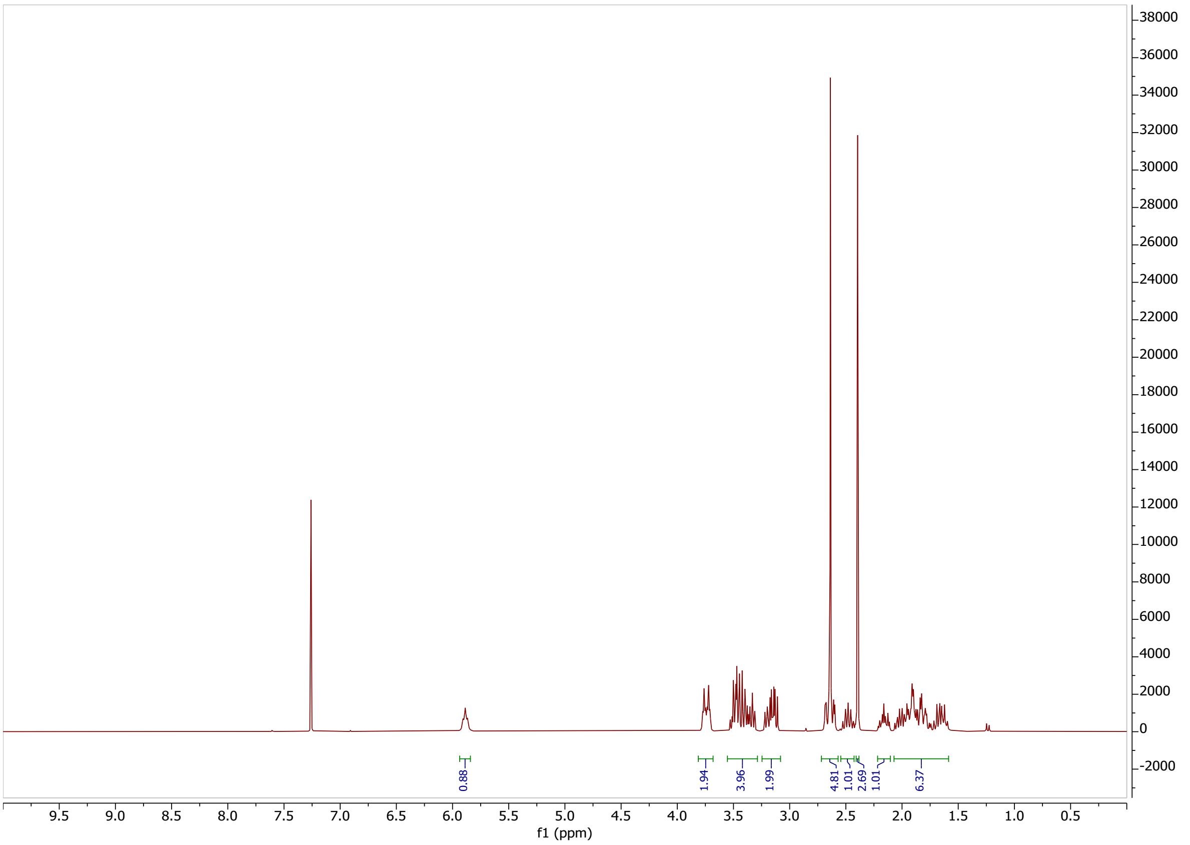

#### $^{13}$C-NMR: BB11CA282

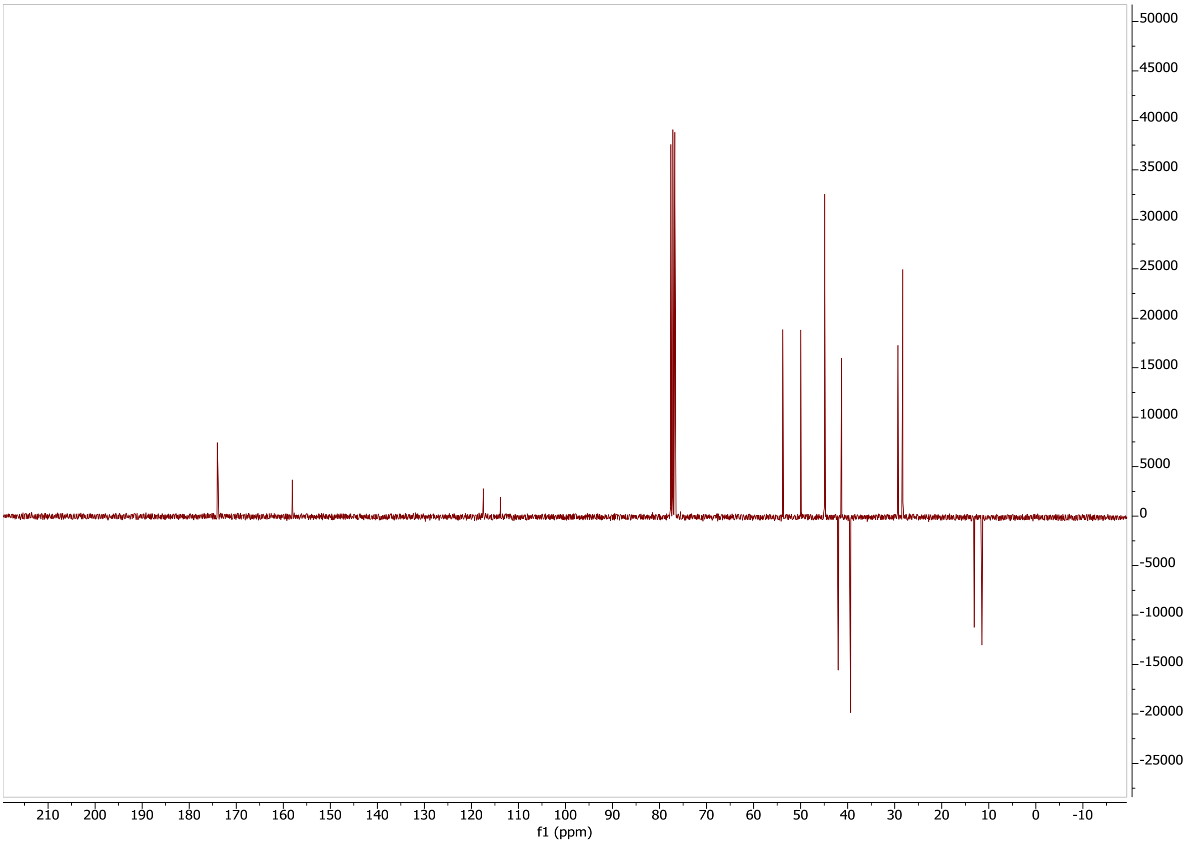

## LC-MS: BB11CA282

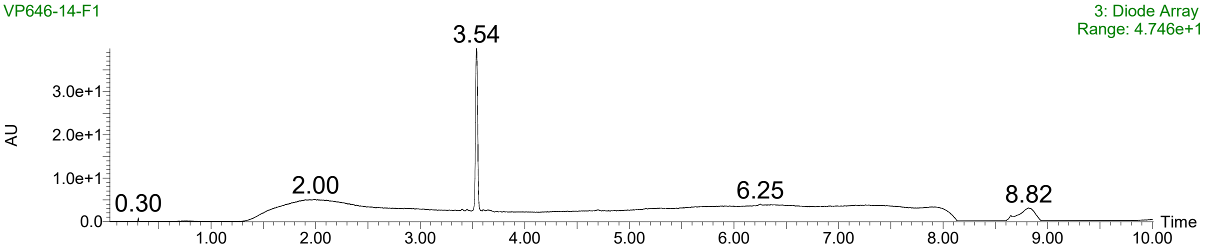

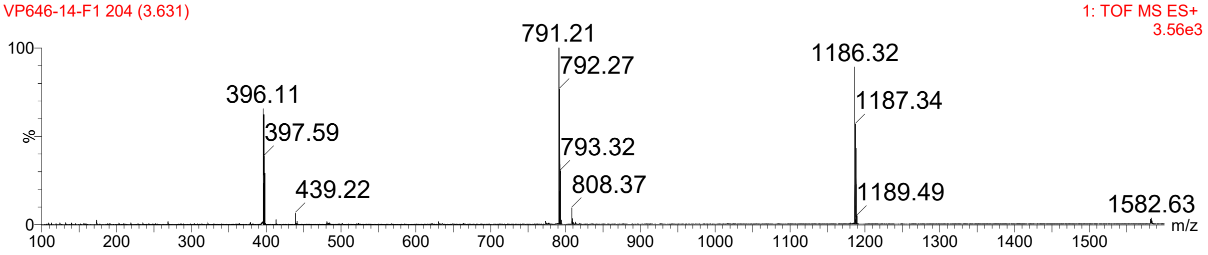

**[1]H-NMR: BB12CA282**

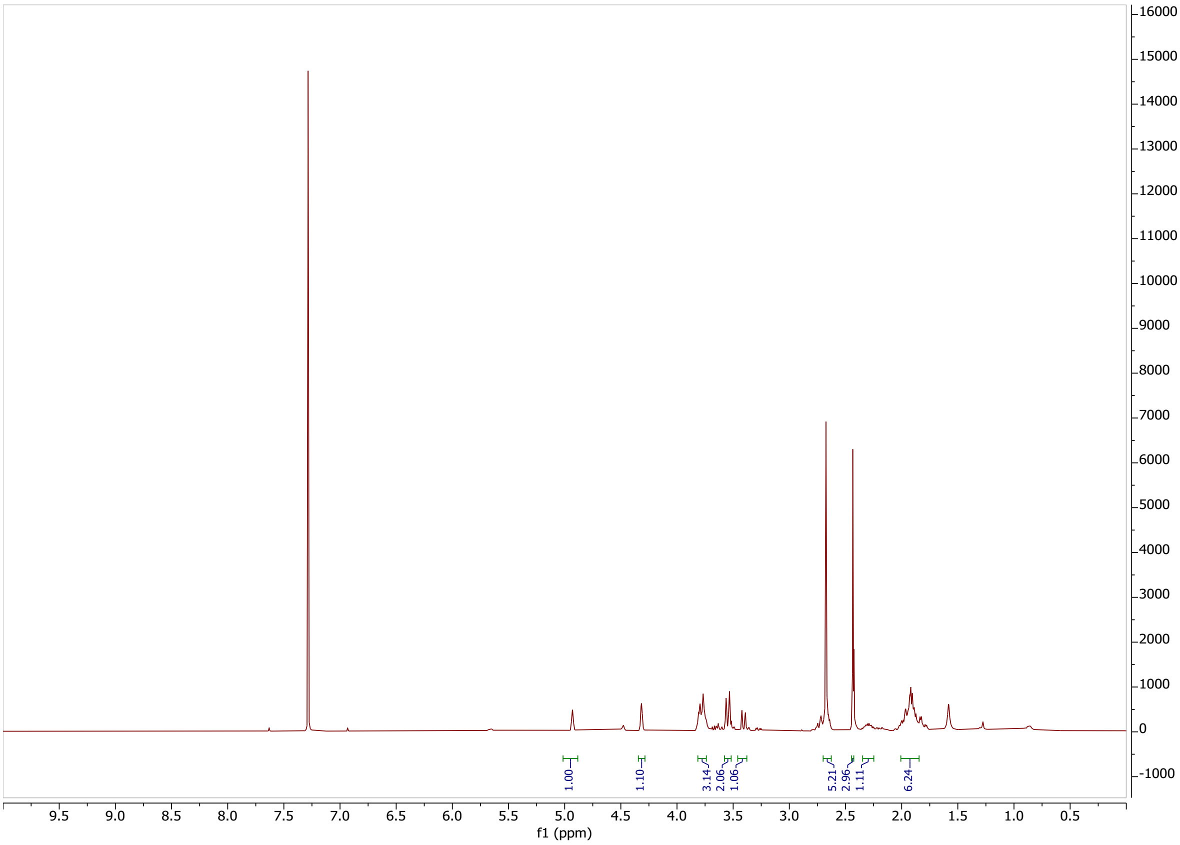

**[13]C-NMR: BB12CA282**

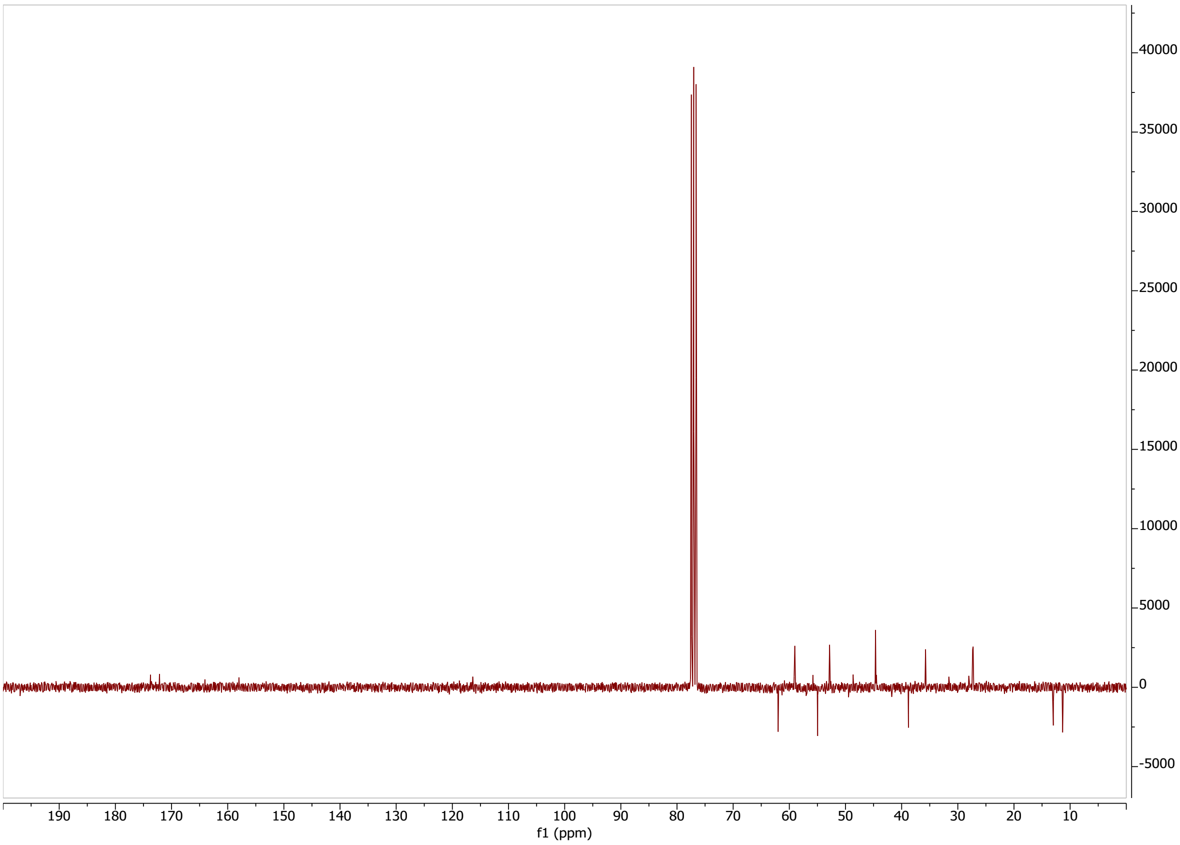

## LC-MS: BB12CA282

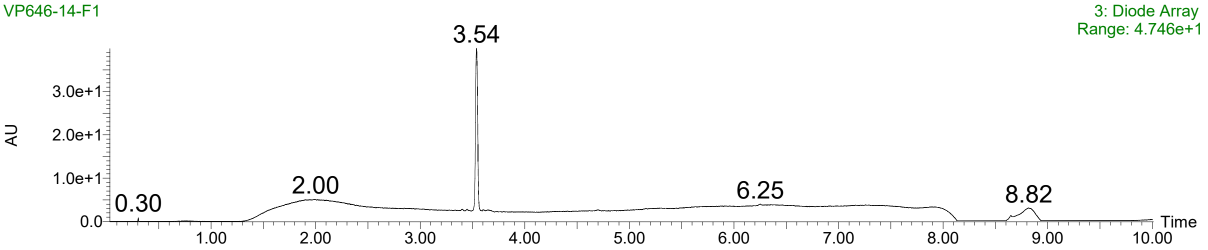

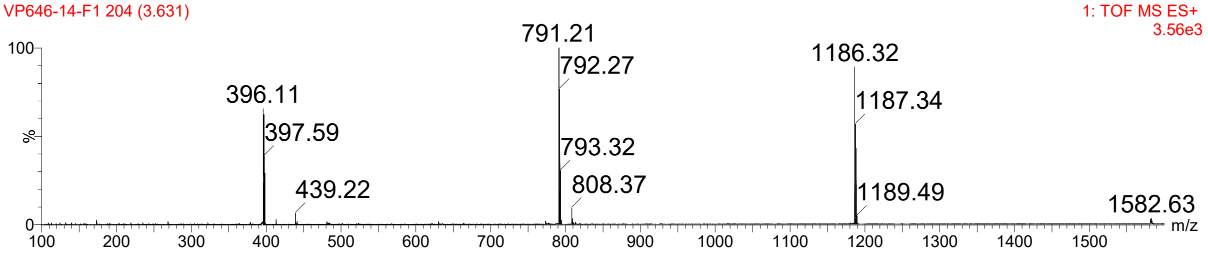

#### [1]H-NMR: BB14CA282

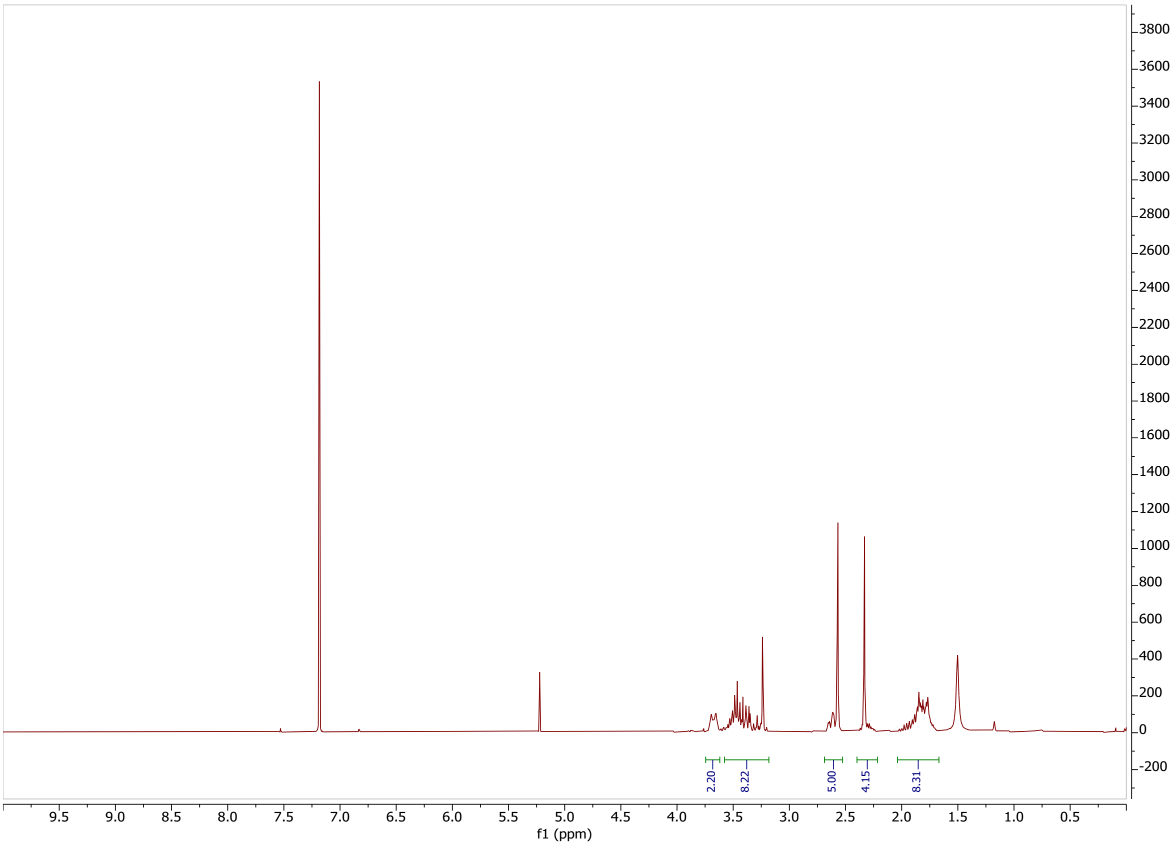

#### [13]C-NMR: BB14CA282

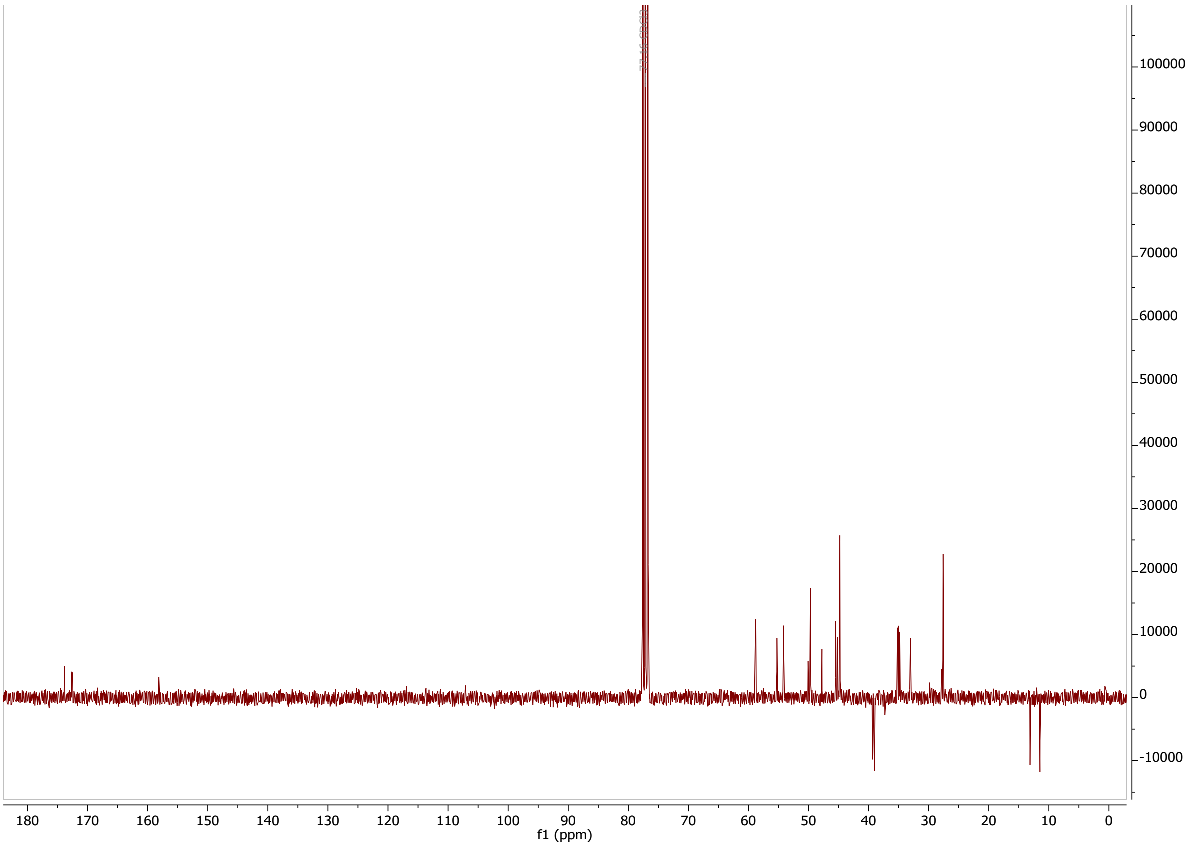

#### LCMS: BB14CA282

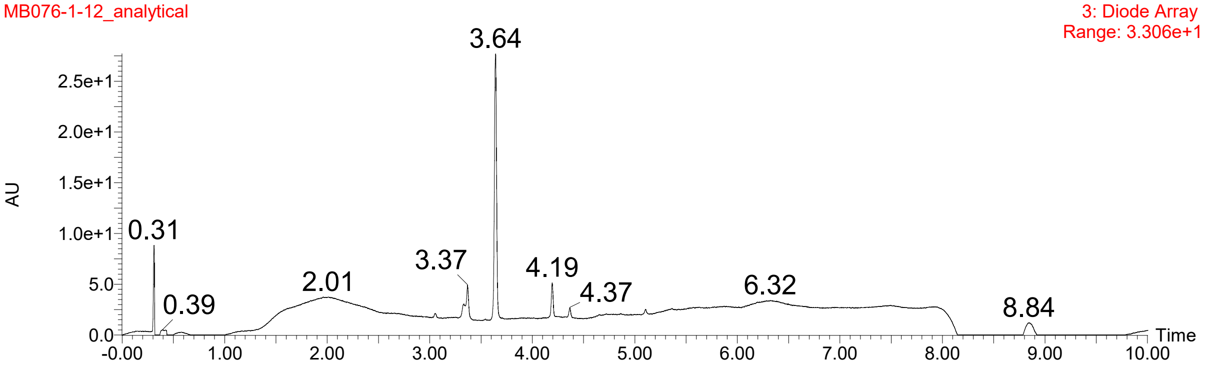

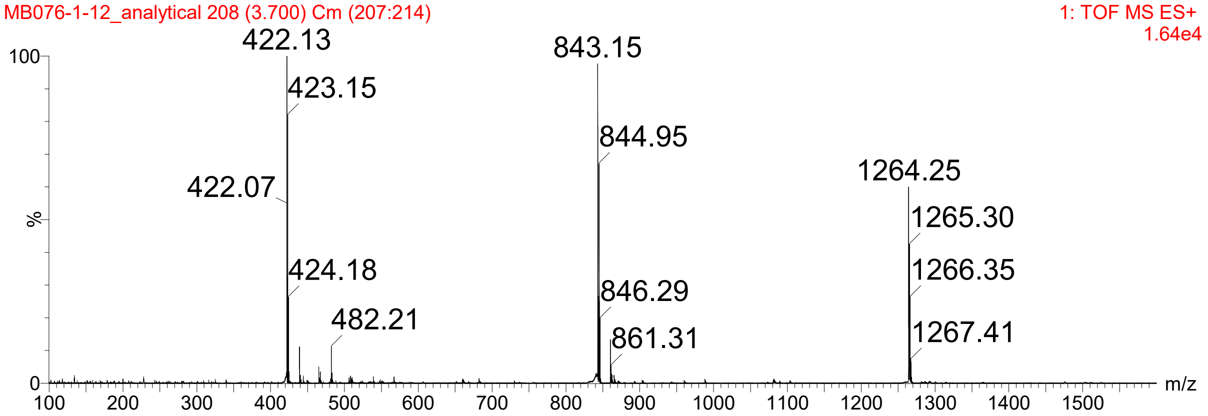

#### [1]H-NMR: Compound 6

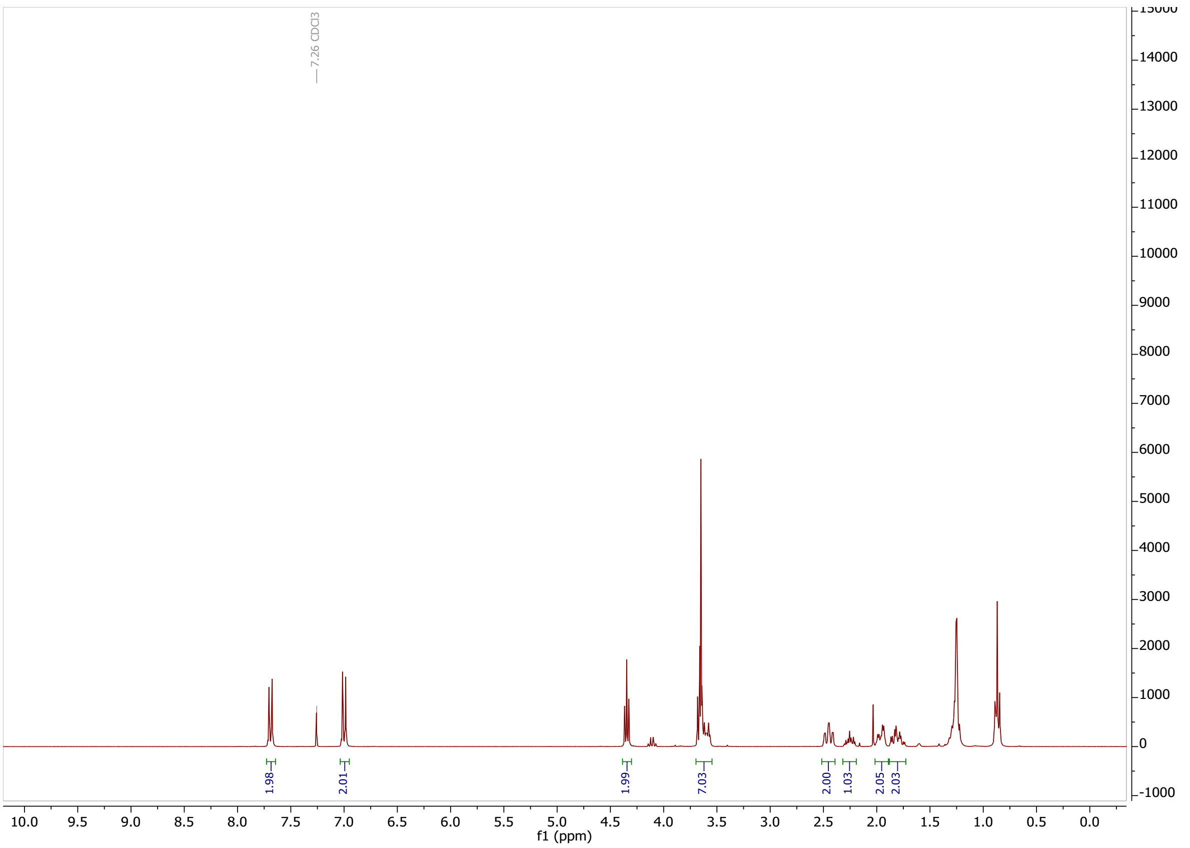

#### [13]C-NMR: Compound 6

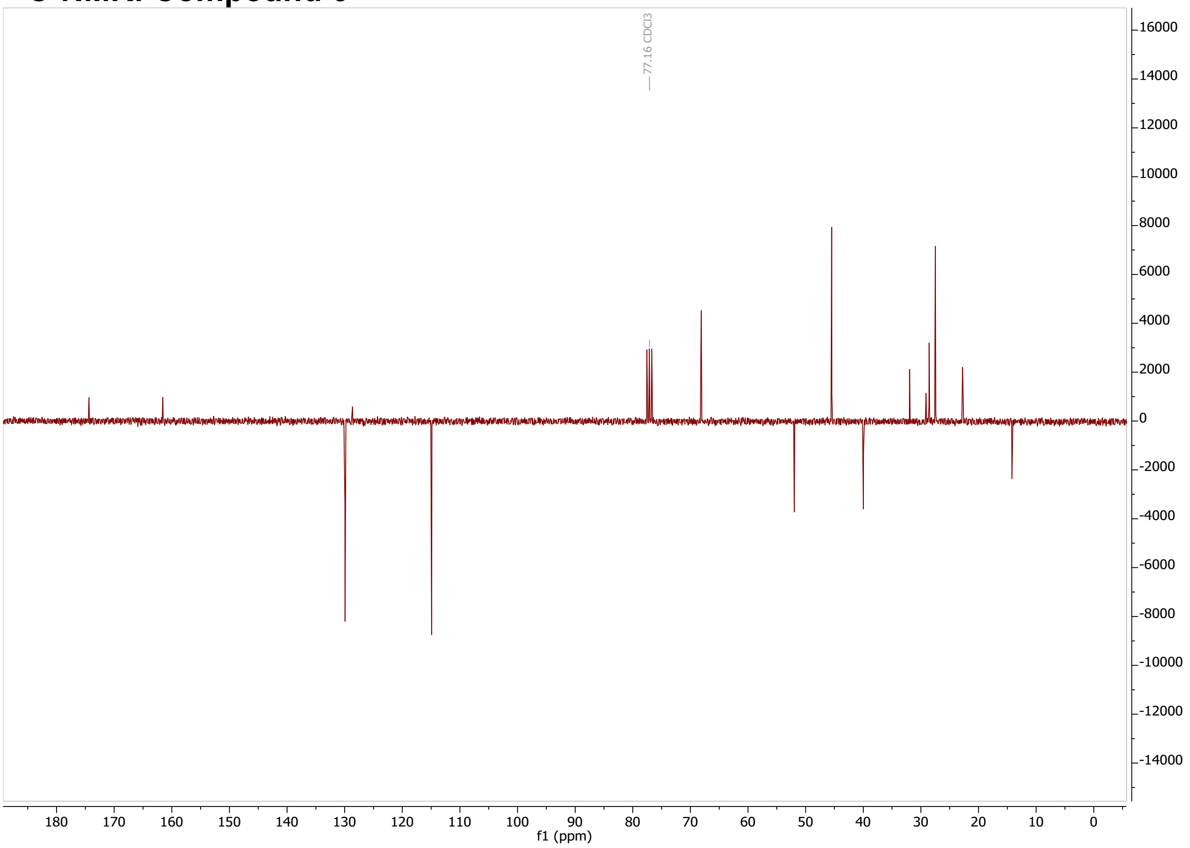

#### [1]H-NMR: Compound 7

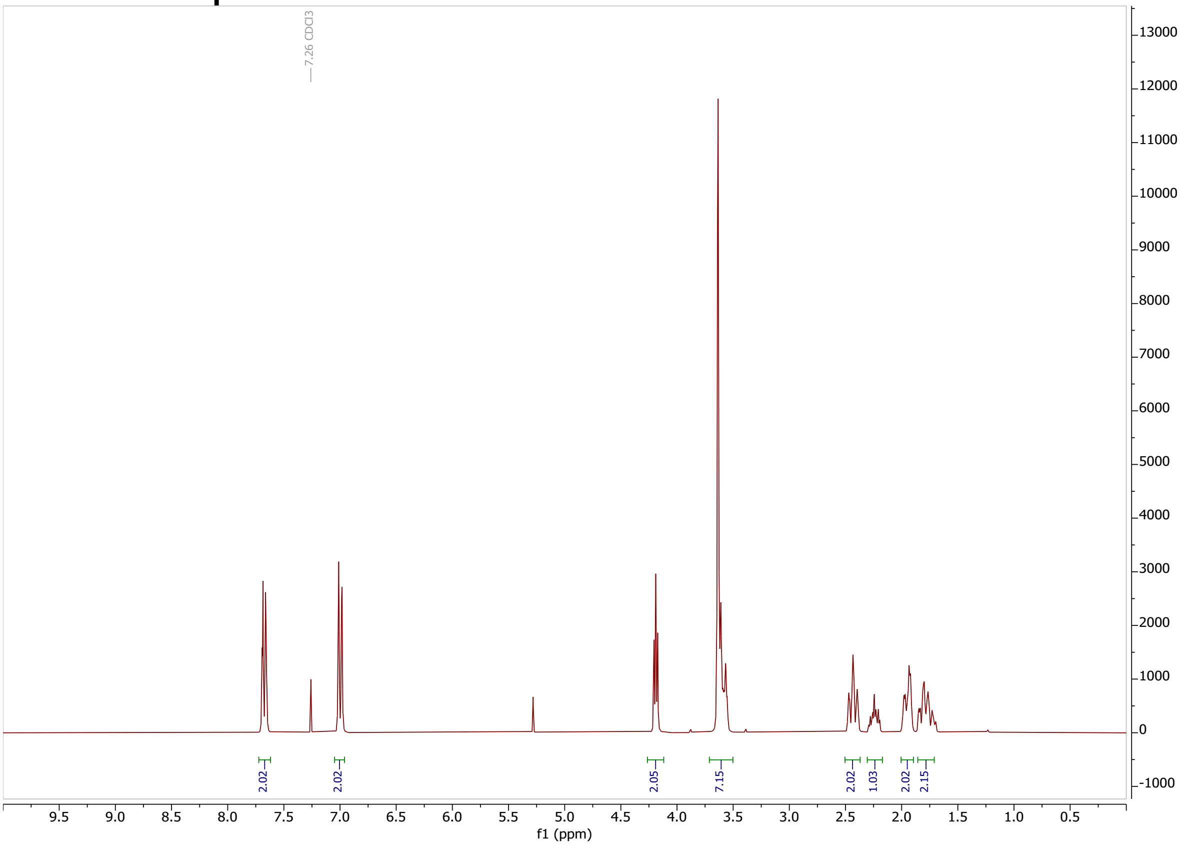

#### [13]C-NMR: Compound 7

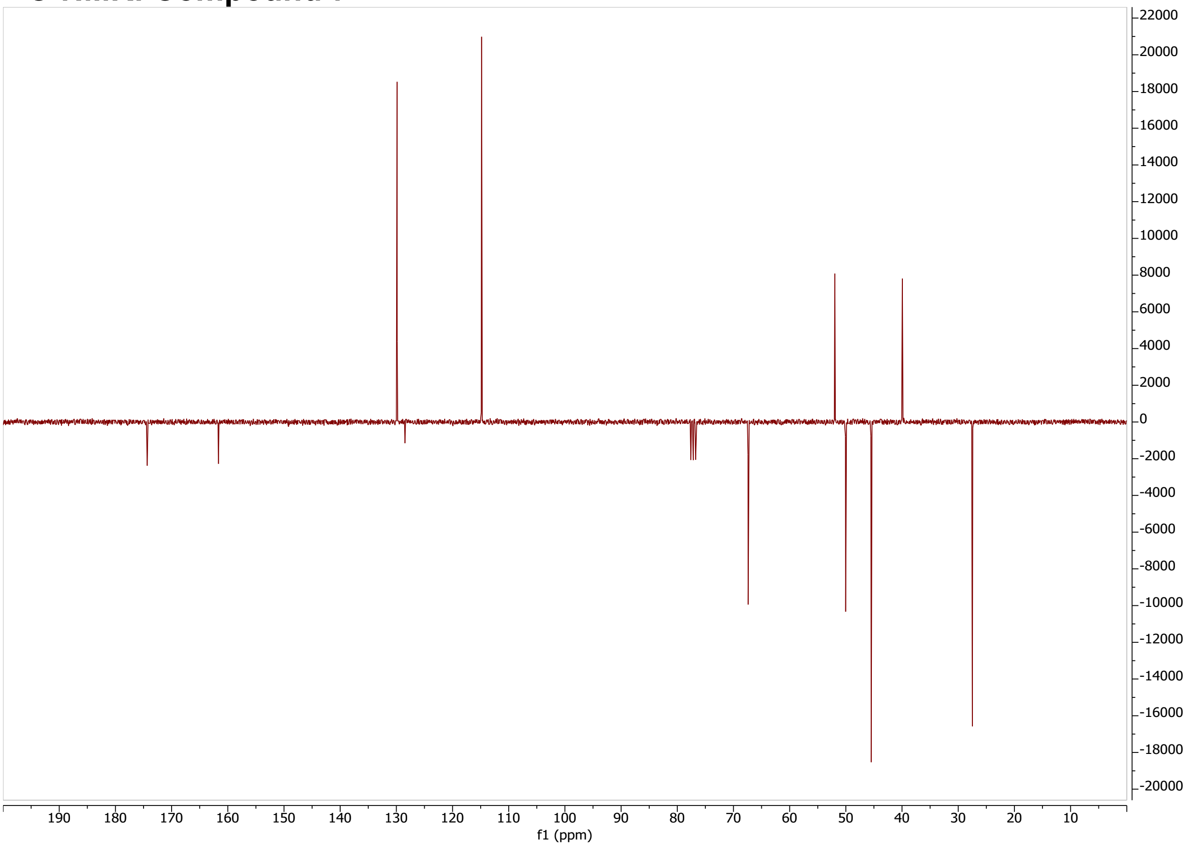

#### $^{1}$H-NMR: Compound 8

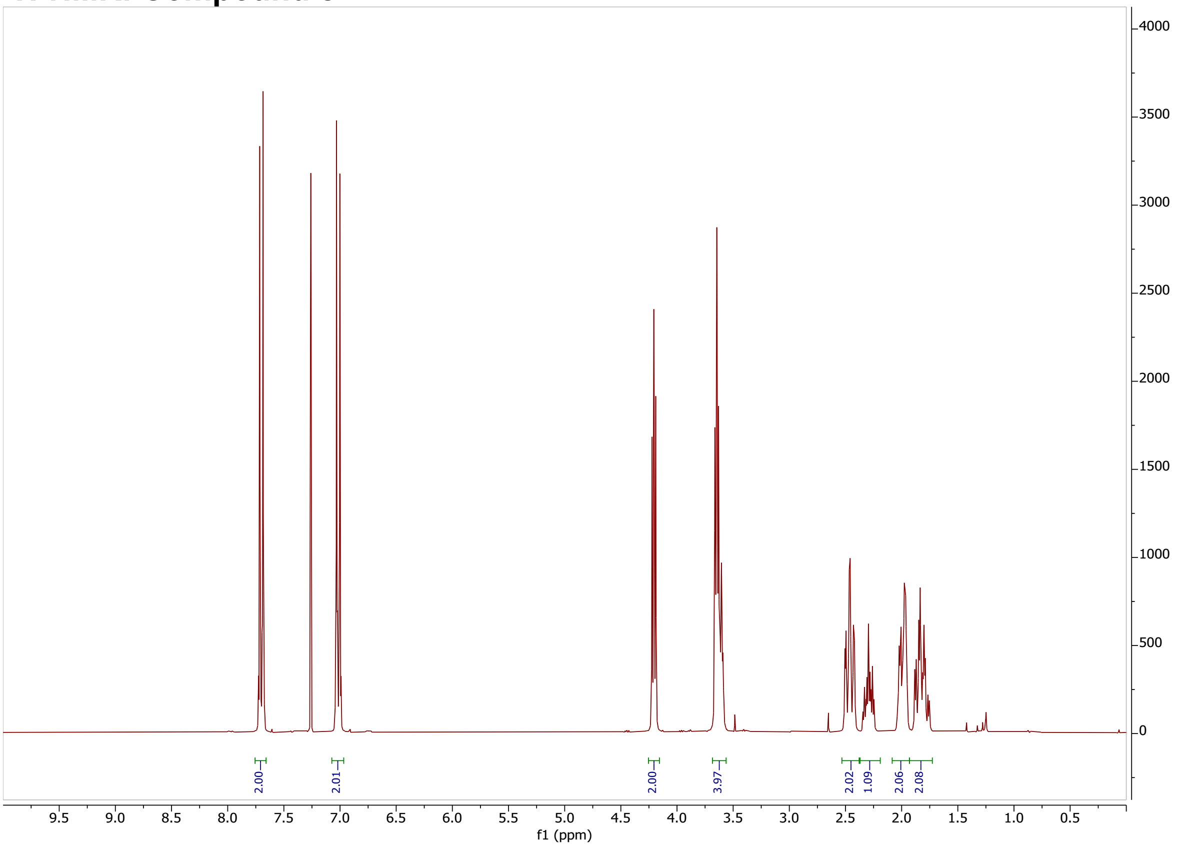

#### $^{13}$C-NMR: Compound 8

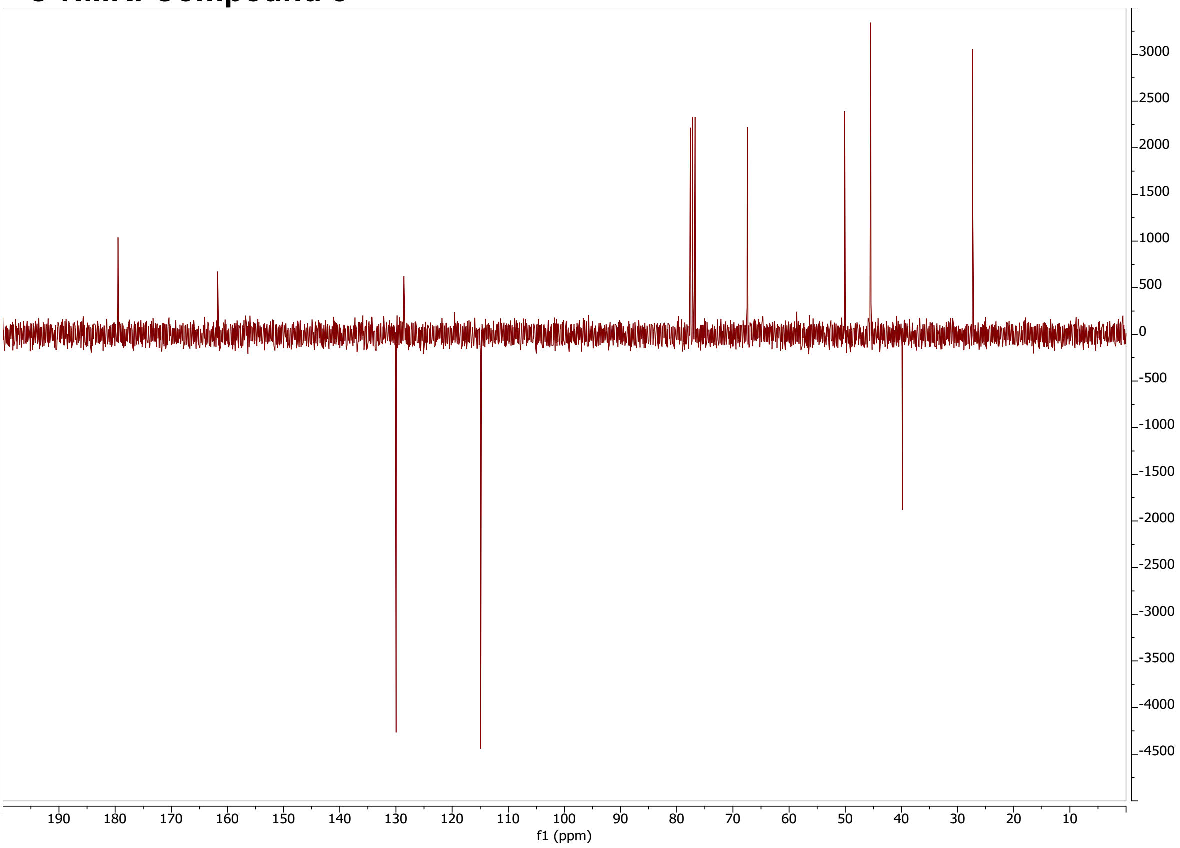

**$^1$H-NMR: Huib32***

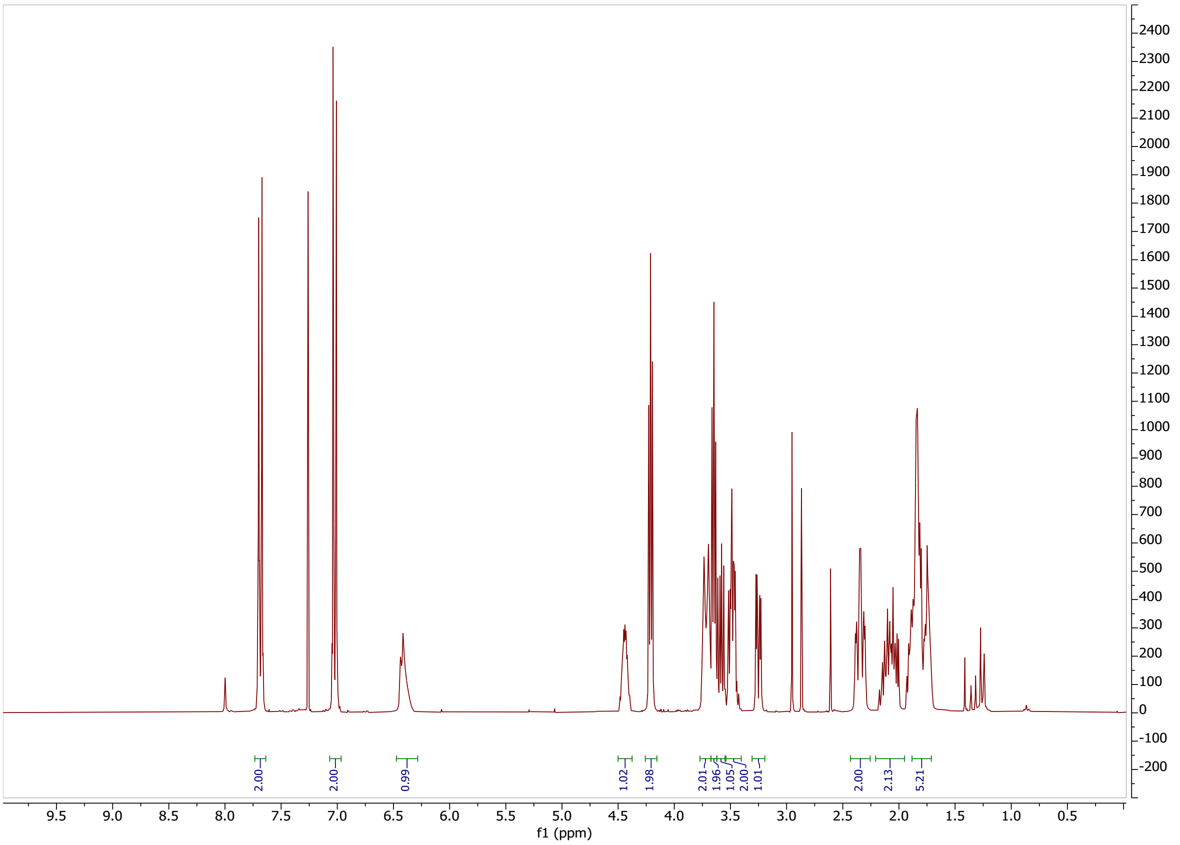

**$^{13}$C-NMR: Huib32***

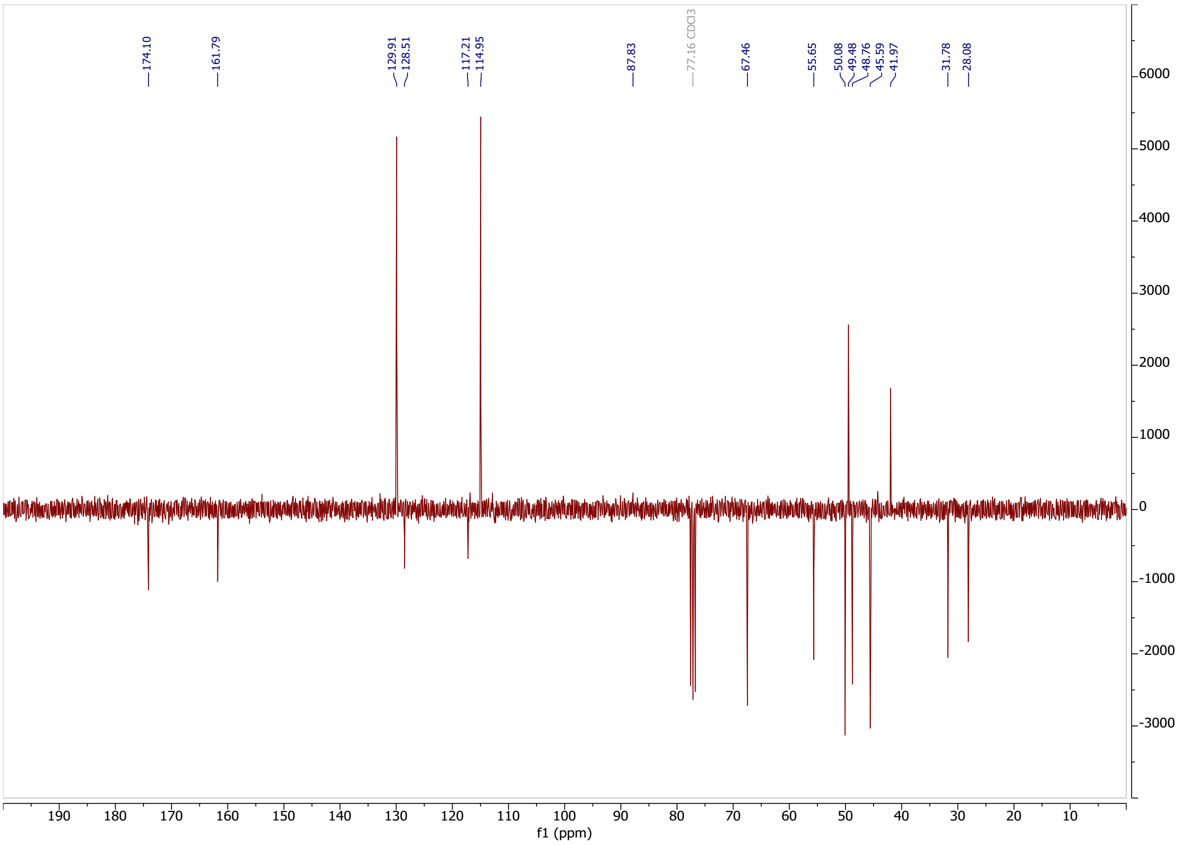

#### LCMS: Huib32*

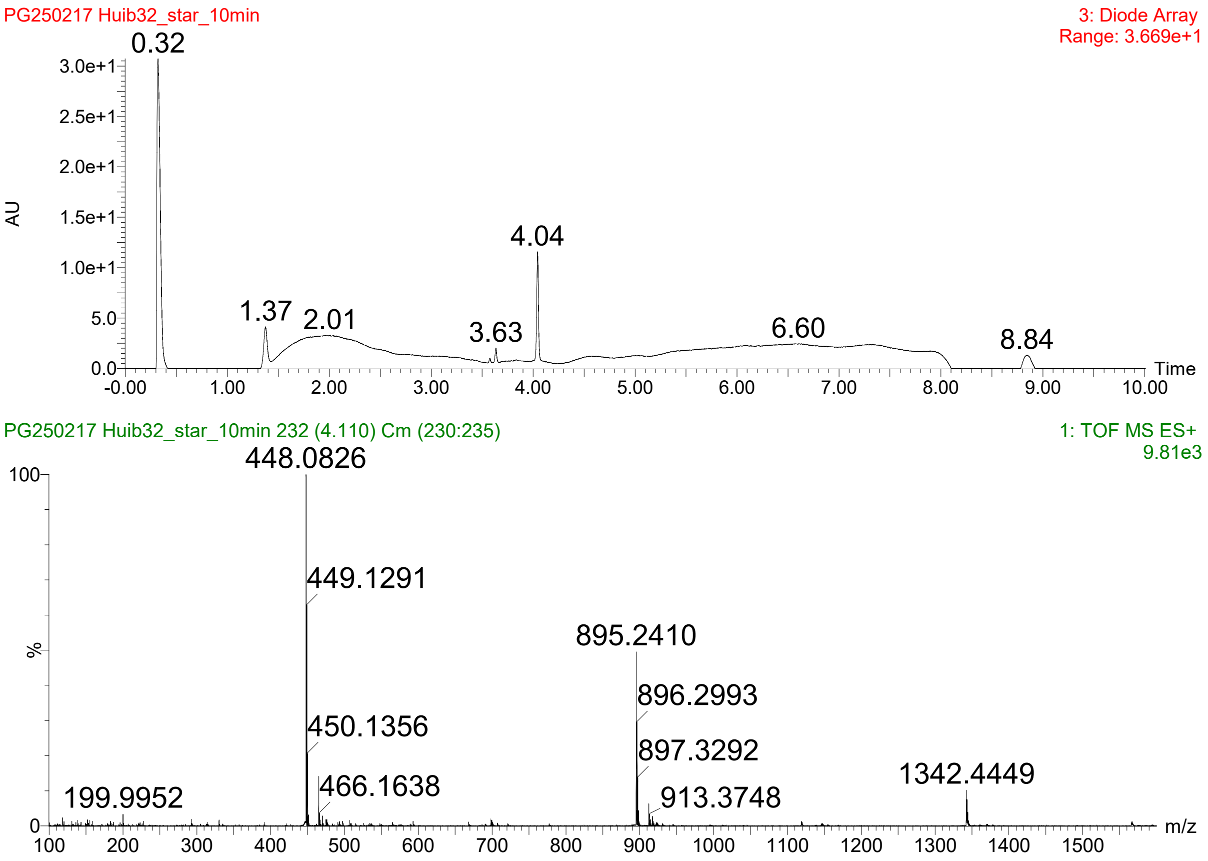
PG250217 Huib32_star_10min
3: Diode Array
Range: 3.669e+1
AU
Time
0.32
1.37
2.01
3.63
4.04
6.60
8.84
PG250217 Huib32_star_10min 232 (4.110) Cm (230:235)
1: TOF MS ES+
9.81e3
%
m/z
448.0826
449.1291
450.1356
466.1638
199.9952
895.2410
896.2993
897.3292
913.3748
1342.4449

#### $^{1}$H-NMR: Huib32*1

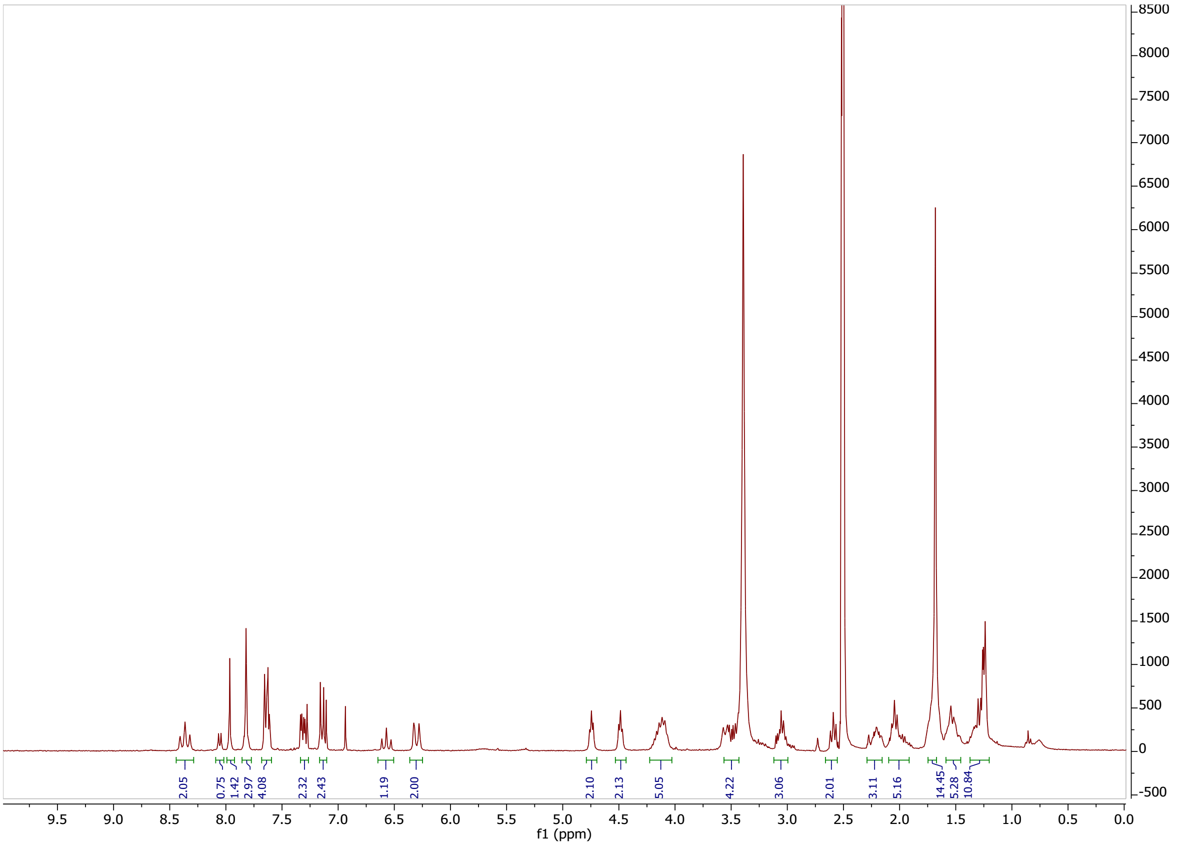

#### $^{13}$C-NMR: Huib32*1

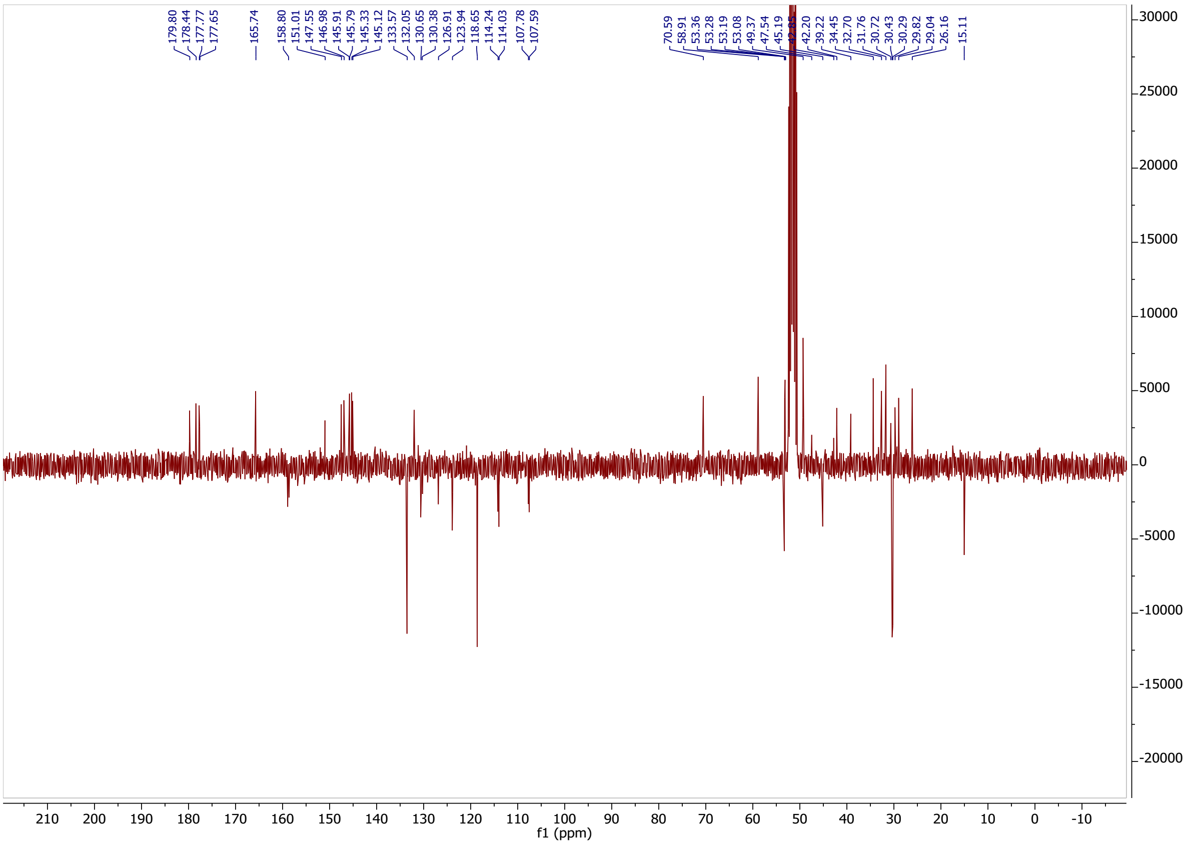

#### LCMS: Huib32*1

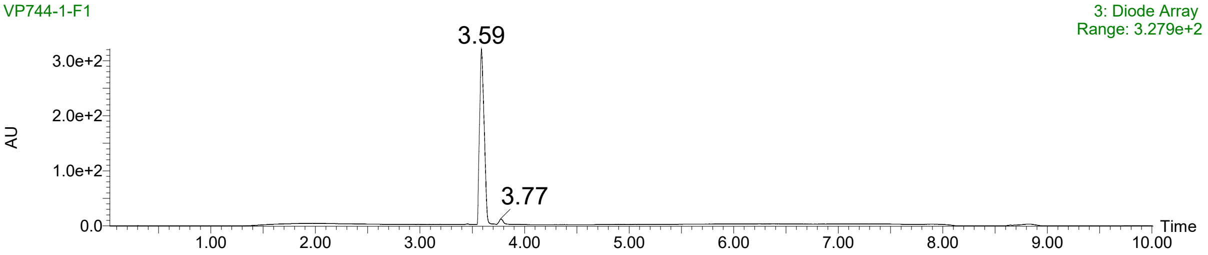

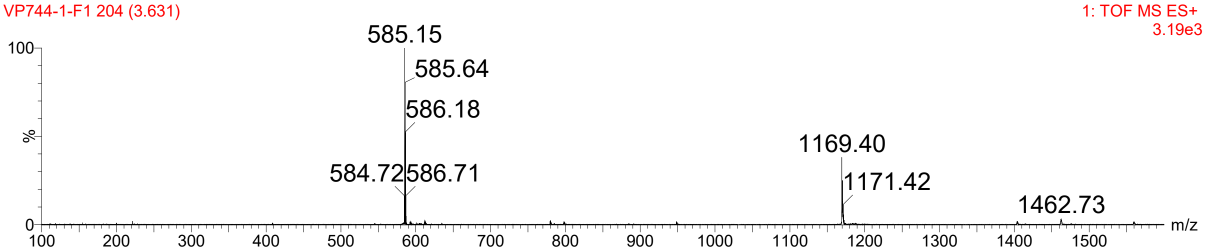

#### $^{1}$H-NMR: Huib32*2

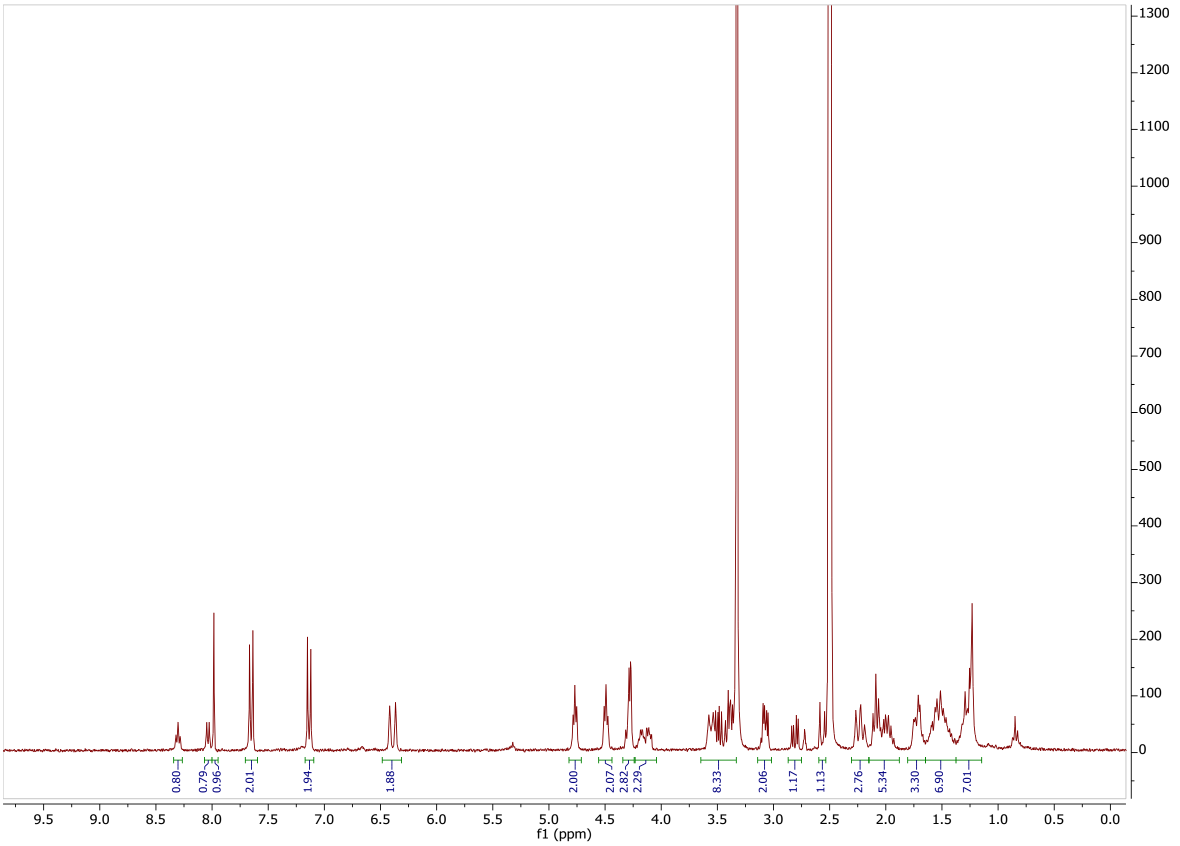

#### $^{13}$C-NMR: Huib32*2

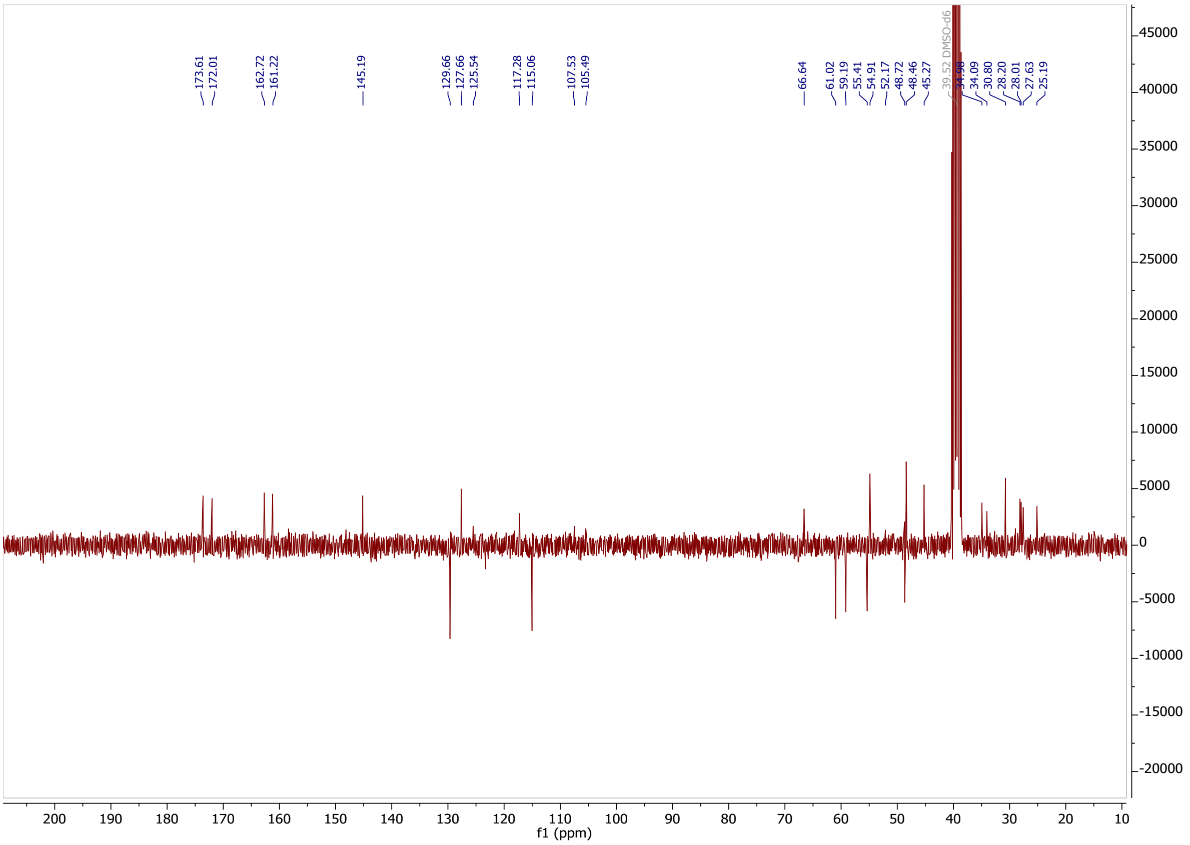

### LCMS: Huib32*2

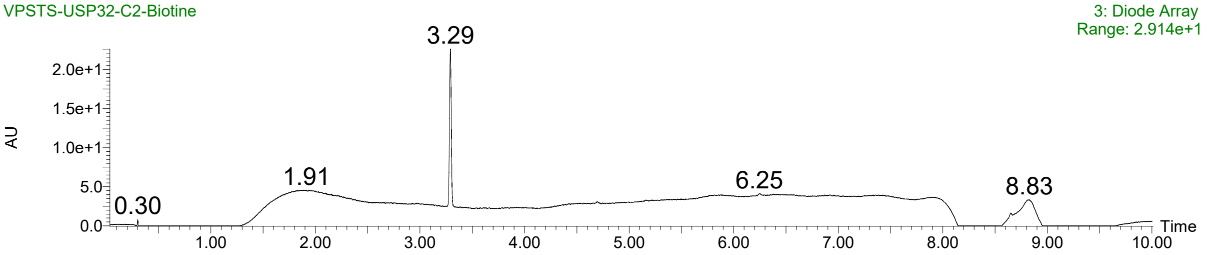

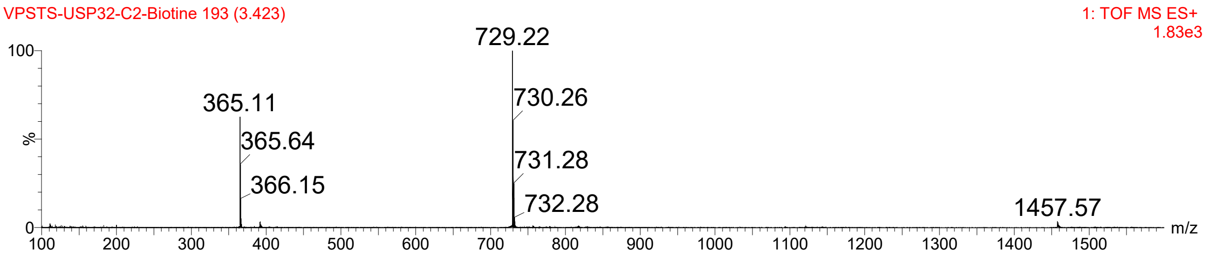

**RhoK(Biotin)UbPA probe.**

**LCMS**

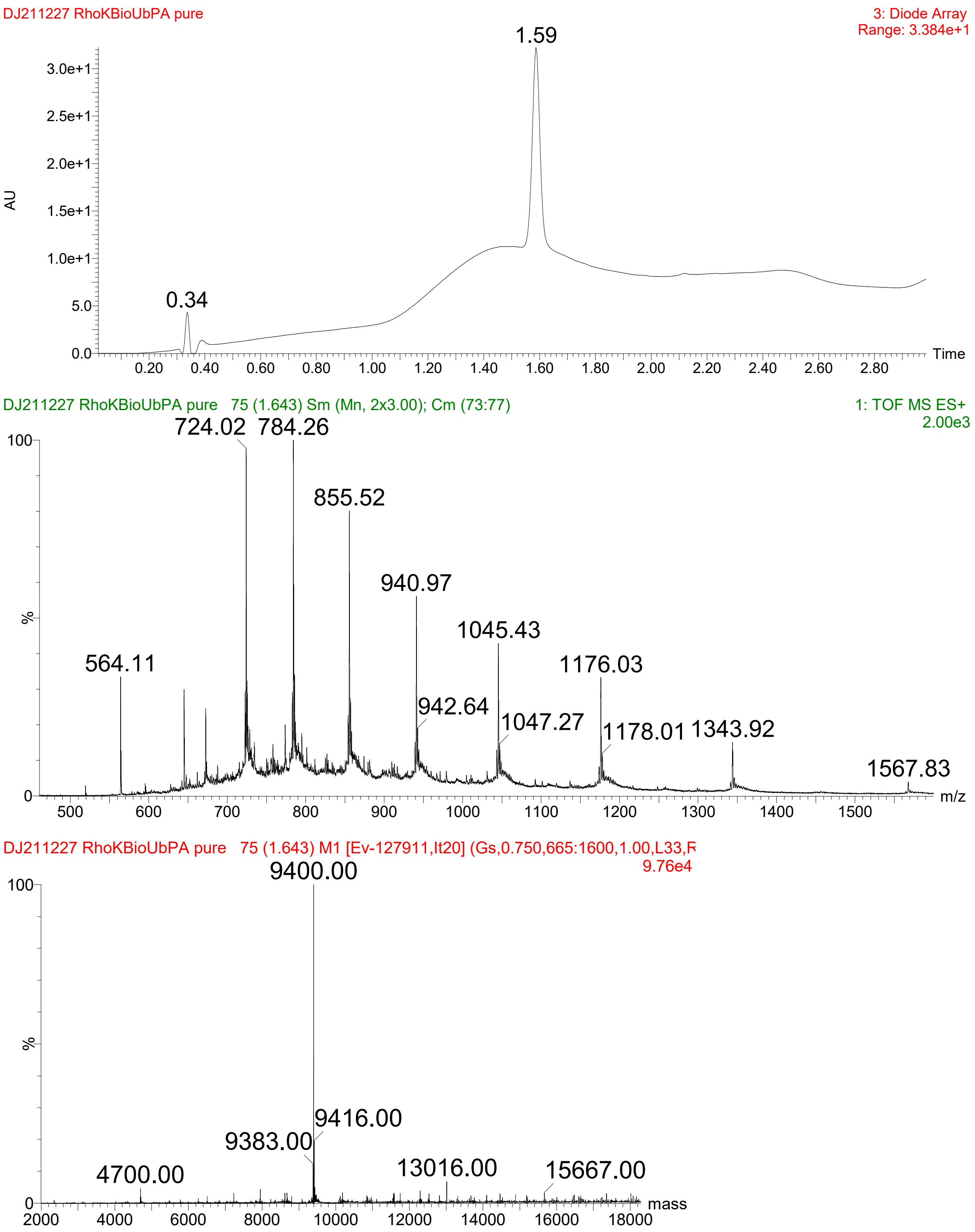
